# A gene-based model of fitness and its implications for genetic variation: Linkage disequilibrium

**DOI:** 10.1101/2024.09.12.612686

**Authors:** Parul Johri, Brian Charlesworth

## Abstract

A widely used model of the effects of mutations on fitness (the “sites” model) assumes that heterozygous recessive or partially recessive deleterious mutations at different sites in a gene complement each other, similarly to mutations in different genes. However, the general lack of complementation between major effect allelic mutations suggests an alternative possibility, which we term the “gene” model. This assumes that a pair of heterozygous deleterious mutations in *trans* behave effectively as homozygotes, so that the fitnesses of *trans* heterozygotes are lower than those of *cis* heterozygotes. We examine the properties of the two different models, using both analytical and simulation methods. We show that the gene model predicts positive linkage disequilibrium (LD) between deleterious variants within the coding sequence, under conditions when the sites model predicts zero or slightly negative LD. We also show that focussing on rare variants when examining patterns of LD, especially with Lewontin’s *D*’ measure, is likely to produce misleading results with respect to inferences concerning the causes of the sign of LD. Synergistic epistasis between pairs of mutations was also modeled; it is less likely to produce negative LD under the gene model than the sites model. The theoretical results are discussed in relation to patterns of LD in natural populations of several species.

## Introduction

There is much interest in the effects of deleterious mutations on properties of populations such as their genetic load, inbreeding depression, and variance in fitness, tracing back to the pioneering work of Haldane (1937), Muller (1950), Morton et al. (1956) and Crow (1958), and to more recent studies of the genetics of *Drosophila* fitness components (reviewed by Crow 1993; Charlesworth 2015). In addition, there have been a number of recent theoretical studies of measures of linkage disequilibrium (LD) among deleterious mutations (e.g., Garcia and Lohmueller 2021; Roze 2021; Good 2022; Ragsdale 2022; Lyulina et al. 2004), stimulated by population genomic studies where the sign of LD among rare, putatively deleterious variants has been used to make inferences concerning features such as epistatic fitness interactions and Hill-Robertson interference (Pool et al. 2012; Sohail et al. 2017; Sandler et al. 2021; Garcia and Lohmueller 2021; Ragsdale 2022; Stolyarova et al. 2022).

The theoretical predictions are mostly based on population genetics models of diploid organisms that tacitly assume that nucleotide sites subject to mutation and selection can be treated as independent of each other with respect to their fitness effects, and of whether they are in the same or different genes (reviewed by Kyriazis et al. 2023). Models of this type were first introduced by Haldane (1937), who assumed that fitnesses were multiplicative across loci. Consider individuals that are heterozygous at two different nucleotide sites. Assume that the fitnesses of the two types of double heterozygotes are the same, regardless of whether or not the two mutations are carried on the same haplotype (*cis*), or on the maternal and paternal haplotypes (*trans*) – the “sites” model (see Figure 1A). With the same selection coefficients against homozygotes (*s)* and dominance coefficients (*h*) at each site, the fitnesses of these individuals are both (1 – *hs*)^2^ if effects are multiplicative across sites and 1– 2*hs* if effects are additive. If selection is sufficiently weak that *s*^2^ << *s*, as is assumed here, the additive and multiplicative models are nearly equivalent. For mathematical convenience, we will use the additive model from now on.

**Figure 1:**
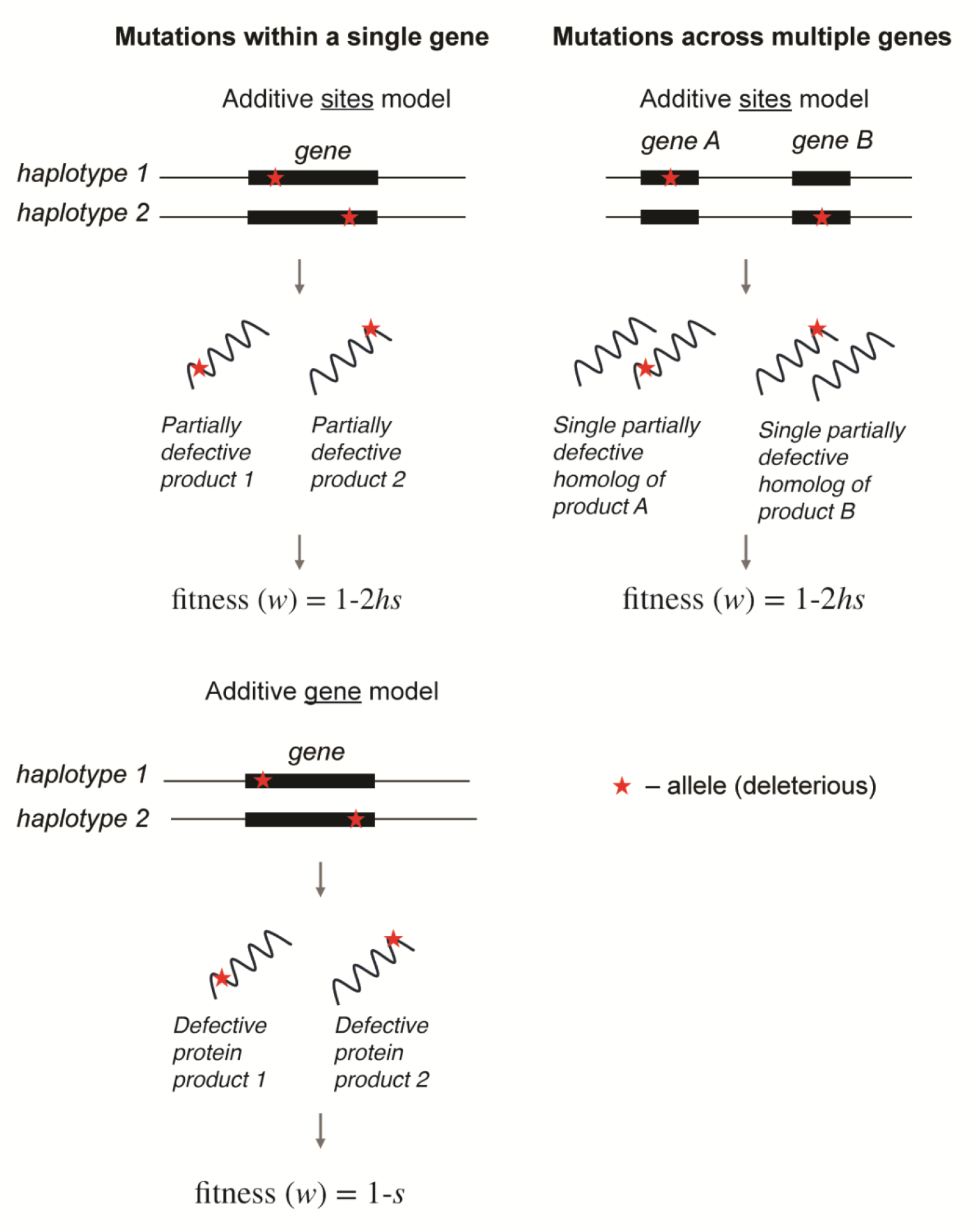
A schematic of the additive gene versus additive sites models. Note that the additive sites model is approximately equivalent to the standard multiplicative fitness model.

In contrast, if the gene is treated as a single entity that is subject to mutation, and there is no allelic complementation between mutations within the same gene (more accurately, a cistron as defined by Benzer [1957]), the fitness of an individual that is heterozygous for two mutations depends on whether or not they are in *cis* or *trans* (Figure 1B) – the “gene model”. The fact that double heterozygotes with mutations in *cis* under the gene model have higher fitnesses than those with mutations in *trans* implies an intragenic position effect with respect to fitness when *h* < ½. Other things being equal, this is likely to favor positive linkage disequilibrium (LD), if the sign of the coefficient of linkage disequilibrium (*D*) is defined as positive when the frequencies of haplotypes carrying pairs of deleterious mutant alleles and pairs of wild-type alleles are in excess of random expectation. This is because the lower fitness of *trans* compared with *cis* double heterozygotes when *h* < ½ means that + + and – – haplotypes will be transmitted more frequently to the next generation relative to + – and – + haplotypes than when the two genotypes have the same fitness.

Position effects of this kind have rarely been included in population genetics models, with the exception of a study of two-locus polymorphisms by Nordborg et al. (1995), and the models of mutation and selection at multiple sites within a gene by Clark (1998), Thornton et al. (2013) and Sanjak et al. (2015), and at two sites by Ragsdale (2022), who all used somewhat different fitness models from ours. Such gene-based fitness models are likely to result in patterns of LD between deleterious mutations that are different from those expected under the sites model (Ragsdale 2022).

The purpose of the present paper is to examine the consequences of the differences between the gene and sites models for patterns of linkage disequilibrium (LD) among variants at sites under selection, using a model of a single coding sequence subject to deleterious mutations. We do not attempt to present a completely general model of a multi-site system, but instead analyze a simple model that brings out the main differences between the two different representations of mutational effects on fitness. An analytical treatment of a model with two sites under selection in an infinite, randomly mating population is given first, followed by stochastic simulations of a multi-site sequence that is intended to provide an approximate description of a coding sequence. We show that the difference between the two models has significant consequences for the patterns of LD among such mutations, with the gene model leading to positive LD among pairs of deleterious variants under conditions where the additive model is associated with zero or slightly negative LD. Interpretations of such patterns in natural populations need to take into account the possibility that the gene model is likely to be a more accurate representation of the dominance properties of deleterious mutations. We also show that focusing on rare variants, as advocated by Garcia and Lohmueller (2021) and Good (2022), is likely to produce estimates of positive LD among deleterious variants in agreement with Good (2022); the opposite finding of Garcia and Lohmueller (2021) is due to their use of Lewontin’s *D*’ statistic (Lewontin 1964), which is biased towards positive values when rare variants are used. The consequences of the differences between the two models for the genetic load, inbreeding load, and genetic variance in fitness are described in the companion paper (Johri and Charlesworth 2025).

## Methods

### Multi-locus simulations

Forward simulations were performed using SLiM 4.0 (Haller and Messer 2023), with a Wright-Fisher population of *N* =1000 diploid individuals, a mutation rate of *u* = 0.5 × 10^−5^ per nucleotide site/generation, and a rate of crossing over of 10^−5^ per nucleotide site/generation. A gene with 1000 selected sites was simulated, assuming a dinucleotide model, *i*.*e*, there were two alleles at each site: a wildtype (denoted by +) and a selectively deleterious mutant (denoted by –) allele. Note that the two alleles were equally likely to mutate to each other, i.e., the + allele could mutate to the – allele at rate *u* and vice versa. This model contrasts with similar previous studies (Garcia and Lohmueller 2021; Good 2022; Ragsdale 2022), where mutation was unidirectional from + to –. This procedure has the advantage that it allows a true stationary distribution of allele frequencies to be reached (Wright 1931), in contrast to models with one-way mutation, where all sites will eventually become fixed for mutant alleles. It thus provides a more realistic description of evolution resulting from mutations involving base substitutions. With sufficiently strong selection (*N*_*e*_*s* >> 1), where *N*_*e*_ is the effective population size and *s* is the selective disadvantage to a mutant homozygote (see Figure 1), there is little effect of back-mutation.

Simulations were run either assuming the same selection coefficient at each site or a distribution of fitness effects (DFE). In the first case, the selective disadvantage of mutant homozygotes scaled by twice the population size was fixed at *γ* = 2*Ns* = 2, 20, 100, or 1000. When simulating a distribution of fitness effects (DFE), a gamma distribution was assumed for most of the results presented here, with shape parameter *α* = 0.3 and mean scaled selection coefficients of 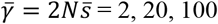, or 1000. In this case, the homozygous selective disadvantage of a – allele at a particular site remained constant throughout a given simulation, although it varied across simulation replicates. The dominance coefficient (*h*) of the mutant allele (see Figure 1) was fixed at 0.0, 0.2, or 0.5. The fitness of a diploid individual was calculated using either the gene or the sites model, as described below. All sites (polymorphic and fixed) were included when calculating the fitnesses of individuals.

Simulations were run for 10*N* generations from an initial population of *N* identical diploid individuals with no variation. For each scenario, 1000 replicates were simulated, and used to determine the means, standard deviations and standard errors across replicates.

#### *Relating the simulations to* Drosophila melanogaster *populations*

These simulation parameters can be placed in the context of the *D. melanogaster* genome, for which there are on average about 5 exons per gene with a mean length of ∼300 bp (Misra et al. 2002). As only about two-thirds of coding sites are nonsynonymous, there are expected to be ∼1000 (= 300 × 5× 2/3) selected sites per gene, corresponding to our simulations. Intron lengths, and hence gene lengths, vary across the genome, with a mean and median length of 5 kb and 2 kb respectively; we have not included introns or synonymous sites in our simulations. Assuming *N*_*e*_ = 10^6^ for *D. melanogaster* and a sex-averaged rate of crossing over of 10^−8^ per site/generation (Johri et al. 2020), *N*_*e*_*Gr ≈* 20 − 50 over a genic region, where *G* is the mean length of a gene. With our simulation parameters, the 1000 selected sites have *N*_*e*_ *Gr* = 10 when rescaled to a population size of 10^6^, and therefore approximately represents a genic region of *D. melanogaster*. Silent site polymorphism data imply that *N*_*e*_ *u ≈* 0.003 − 0.005 in *D. melanogaster* East African populations (Johri et al. 2020), consistent with the *Nu* = 0.005 used in our simulations. This rescaling uses the principle that, when all evolutionary forces are weak, the outcome of evolution is determined by the products of the effective population size and the magnitudes of the deterministic forces (e.g., Ewens 2004, Chap. 5).

Because our simulations were computationally very expensive, it was not feasible to perform them with a large population size. Rescaling by a large scaling factor in forward-in-time simulations can affect population genetic statistics (Dabi and Schrider 2024; Ferrari et al. 2025); rescaling has, however, been found to produce accurate results in the absence of recombination (Kaiser and Charlesworth 2010). As we only simulated short regions of 1000 selected sites, rescaling is likely to produce accurate results in our case as well, since the frequency of crossing over between the most widely separated sites is nearly proportional to the distance between them, so that the recombination rates can be scaled by population size when using a fixed rate of crossing over per basepair. This would not be the case for large distances between sites, for which double crossovers cause a non-linear relation between distance and recombination rate. This means that multiplication of the rate of crossing over per basepair by a given factor does not result in a proportional change in the frequency of recombination. To test the effects of rescaling, additional simulations were conducted with a larger population size of 5000 diploid individuals, while maintaining the same values of *Nu* (*u* = 2 × 10^−6^ per site/generation) and *Nr* (*r* = 2 × 10^−6^ per site/generation) as the simulations above. Selection coefficients followed a gamma distribution with an expected value of 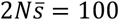 and a shape parameter of 0.3.

#### Simulations of human populations

A Wright-Fisher population of *N* = 1000 diploid individuals was simulated with a mutation rate *u* = 2.5 × 10^−7^per nucleotide site/generation and crossing over rate (*r*) of 2 × 10^−7^per nucleotide site/generation. This resulted in *Nu* = 0.00025 and *Nr* = 0.0002 in our simulations, which are consistent with the estimates of these parameters for human populations: *N*_e_ of 20,000, mutation rate of 1.25 × 10^−8^ per site/generation (Kong et al. 2012), and recombination rate of 1.0 × 10^−8^ per site/generation (Altshuler et al. 2005). Assuming that there are 8 exons per gene on average (Johri et al. 2021), that each exon is about 350 bp, and that 70% of all coding sites in humans are nonsynonymous (Kryukov et al. 2007), there should be about 1960 selected sites in a gene. We therefore simulated 2000 selected sites. We again ignored introns; because genes tend to be coldspots for recombination in many species (e.g., Myers et al. 2010; Joseph et al. 2024), the presence of introns should not greatly affect recombination rates within genes. The DFE was assumed to be a gamma distribution with shape parameter *α* = 0.23 and 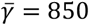, as estimated by Eyre-Walker et al. (2006) and Kim et al. (2017).

### Modeling the fitnesses of individuals

As mentioned in the Introduction, two different models of main effects on fitness (the sites and gene models) are used to calculate the fitnesses of diploid genotypes. The sites model assumes that heterozygous mutations at all sites complement each other fully. The gene model assumes a complete absence of complementation between heterozygous mutations present in *trans*. The difference in fitness between *cis*-and *trans*-heterozygotes can be regarded mathematically as a form of epistasis, although its biological basis is different from the type of epistasis usually considered in population genetics models.

In both cases, additivity of the main effects across sites was assumed in most of our numerical work, as was also done by Good (2022) and Ragsdale (2022), in contrast to the multiplicative fitness model that is often used in multi-locus studies. The difference between additive and multiplicative fitness models involves second-order terms in the selection coefficients. For a single gene with a modest number of segregating sites, as simulated here, there should be little difference between additive and multiplicative models, unless selection is much stronger than is generally assumed here. As just described, our simulations use rescaling of the deterministic parameters, so that their products with *N*_*e*_ match those for much larger natural populations. The second-order terms in the selection coefficients involved in multiplicative fitness models makes such rescaling problematical; this difficulty is avoided when the additive model used.

In addition to the main effects of each site, a simple model of pairwise epistasis of the standard type was used, in which a single parameter (*ϵ*) describes a pairwise interaction effect on fitness. To avoid confusion with the effect of non-complementation under the gene model, we will refer to this as either antagonistic or synergistic epistasis. The total number of sites within a gene is denoted by *G*. The parameter *ϵ* is positive if there is synergistic epistasis, so that double mutants reduce fitness by more than is expected from the effects of each mutation on its own, and negative if there is antagonistic (diminishing returns) epistasis, where double mutants behave in the opposite fashion (Crow 1970; Kondrashov 1988).

Analytical results are derived in sections 1-3 of the Appendix and in Supplementary File S1 for the case of *G* = 2, with the same selection coefficient *s* at each site. In general, in addition to a possible difference in fitness between *cis*-and *trans*-heterozygotes, there are three possible epistatic coefficients that measure departure from additivity in a symmetric fitness model, denoted by the coefficients *e*_1_, *e*_2_ and *e*_3_ in Table 1. In order to avoid multiple epistatic parameters, we assume that the magnitude of such epistasis scales linearly with the sizes of the selective effects at each site with a constant of proportionality *ϵ*, so that the fitness of a mutant double homozygote is 1 – 2*s –* 2*ϵs*. The fitnesses of all possible genotypes under the sites model are defined in the upper section of Table 1, where the assumption of linear scaling and complete complementation of mutations at different sites in double heterozygotes gives *e*_3_ = 2*ϵs, e*_1_ = 2*ϵhs* and *e*_2_ = *ϵ*(1 + *h*)*s*. For the gene model, a double heterozygote with mutations in *cis* has fitness 1 – 2*hs* – 2*ϵhs*, whereas the absence of complementation in a *trans* double heterozygote implies that its fitness is 1 – *s*.

**Table 1.** Models of the fitness effects of two sites with a fixed selection coefficient.

|  | ++ | + - | - + | - - |
| --- | --- | --- | --- | --- |
| ++ | 1 | $1 - hs$ | $1 - hs$ | $1 - 2hs - e_1$ |
| + - | $1 - hs$ | $1 - s$ | $1 - 2hs - e_1$ | $1 - s - hs - e_2$ |
| - + | $1 - hs$ | $1 - 2hs - e_1$ | $1 - s$ | $1 - s - hs - e_2$ |
| - - | $1 - 2hs - e_1$ | $1 - s - hs - e_2$ | $1 - s - hs - e_2$ | $1 - 2s - e_3$ |

$$e_1 = 2\epsilon hs; e_2 = \epsilon(1 + h)s; e_3 = 2\epsilon s.$$ The “gene” model
|  | ++ | + - | - + | - - |
| --- | --- | --- | --- | --- |
| ++ | 1 | $1 - hs$ | $1 - hs$ | $1 - 2hs - e_1$ |
| + - | $1 - hs$ | $1 - s$ | $1 - s$ | $1 - (1+k)s - e_2$ |
| - + | $1 - hs$ | $1 - s$ | $1 - s$ | $1 - (1+k)s - e_2$ |
| - - | $1 - 2hs - e_1$ | $1 - (1+k)s - e_2$ | $1 - (1+k)s - e_2$ | $1 - 2s - e_3$ |

It is less obvious how to assign the fitness of a genotype for which one site is homozygous for a mutation and the other is heterozygous, e.g., + –/– –. The representation in Table 1 assigns a fitness reduction of (1 + *k*)(1 + *ϵ*)*s* to this genotype. For most of our numerical results, we assume that a lack of complementation implies that the effects of each mutation on the fitness of this genotype combine additively, with the heterozygous site contributing a fitness reduction of ½*s* and the homozygous site a reduction of *s*, so that *k* = 0.5. This giving a net additive effect of 1.5*s* and an epistatic effect of 1.5*ϵs*. With *e*_3_ = 2*ϵs*, we thus have *e*_1_= *ϵhs* and *e*_2_ = 1.5*ϵs*.

An extreme alternative is to assume that *k* = *h*, as for the sites model, so that the single heterozygous site has the same effect as if the other site were wild-type. There are no data available to distinguish between these alternatives. It is shown in section 3 of the Appendix and section 5 of Supplementary File S1 that the second alternative is likely to give a substantially larger positive LD measure for pairs of deleterious mutations, so that our use of the additive assumption is conservative from the point of view of predicting positive LD.

#### Modeling fitness with multiple sites and no epistasis

For describing fitness (*w*) when there are *G* selected sites within a gene, as in the simulations, consider the two haplotypes (1 and 2) of a diploid individual, where *w* = 1 when all sites are wild-type. With the same selection coefficient at all sites, fitness for the gene model is calculated as follows.

If there is at least one – allele in 1 or 2, but not both, we have:

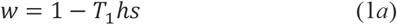

where *T*_1_ is the total number of – alleles over all *G* sites.

If there is at least one – allele in both 1 and 2, we have

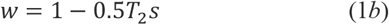

where *T*_2_ is the total number of – alleles present.

In the case of a distribution of fitness effects (DFE), with variable selection coefficient *s*_*i*_ at each site, these equations become:

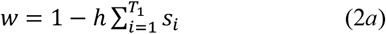

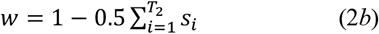

For the sites model with a fixed selection coefficient, we have:

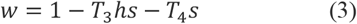

where *T*_3_ is the total number of heterozygous (+/–) sites and *T*_4_ is the total number of mutant homozygous (–/–) sites. With a DFE, Equation (3) becomes:

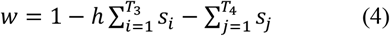

#### The gene model with multiple sites

For the gene model, an extension of the analytical treatment of the two sites model to multiple sites subject to the same selection regime (in the absence of epistasis) is described in section 4 of the Appendix. This is based on the assumptions that LD is sufficiently weak that the distribution of numbers of mutations per haplotype follows a Poisson distribution, that the number of selected sites *G* is large, and that selection is sufficiently strong that the equilibrium frequencies of deleterious mutations at each site are very small. Since significant levels of synergistic or antagonistic epistasis generate LD, we have not included epistasis in this approach. We consider only the gene model, since LD for the sites model without epistasis is expected to be negligible.

#### Modeling epistasis with multiple sites

All epistatic effects are assumed to be between pairs of sites that are either homozygous or heterozygous for mutant alleles, *i*.*e*, there are no higher-order epistatic effects. Each pairwise interaction involves the sum of a joint measure of the fitness effects of the alleles at the pair of sites in question. The simplest model, which applies to both the sites and gene models, is to assume that the epistatic contribution from a pair of homozygous sites *i* and *j* is *ϵ*(*s*_*i*_ + *s*_*j*_); this is summed over all pairs of homozygous sites, and the net fitness reduction of the genotype where all sites are homozygotes for mutations is given by this term and the sum of the *s*_*i*_ over all homozygous mutant sites.

For the sites model, a heterozygous site *i* can be considered as contributing a main effect of *hs*_*i*_ to the fitness reduction of a genotype, and *ϵhs*_*i*_ to each of the interactions with other sites. Its interaction with a homozygous site *j* thus contributes *ϵ*(*hs*_*i*_ + *s*_*j*_) and its interaction with another heterozygous site *k* contributes *ϵh*(*s*_*i*_ + *s*_*k*_). Similarly, a pair of homozygous sites *i* and *j* contributes a fitness reduction of *ϵ*(*s*_*i*_ + *s*_*j*_). The net epistatic contribution is obtained by summing over all pairs of sites, and the net fitness reduction is given by this sum plus the sum of the fitness reductions from each heterozygous and homozygous mutant site. Note that for both the sites and the gene model, if the fitness of a diploid individual was found to be negative in the simulations, it was changed to a value of zero.

For the gene model, a heterozygous individual whose mutations are all in *cis* would have an epistatic component given by the sum over all pairs of *i* and *j* of *ϵh*(*s*_*i*_ + *s*_*j*_). The net fitness reduction is the sum of this quantity plus the sum of the *hs*_*i*_ over all the heterozygous sites. For an individual who is homozygous for a mutation, or heterozygous in *trans*, at more than one site, the sum of the epistatic contributions over all pairs of sites is obtained by weighting *ϵs*_*i*_ for heterozygous sites by ½ instead of by *h*, and *ϵs*_*i*_ for homozygous sites by 1. The net fitness reduction is given by the sum of this quantity and the sum of the main effects across sites, weighting *s*_*i*_ at homozygous sites by 1 and heterozygous sites by ½. This is slightly different from the symmetric two-site model of Table 1, where a heterozygote for two mutations in *trans* behaves like a homozygote.

### Estimating the linkage disequilibrium statistics from the simulations

Summaries of linkage disequilibrium (LD) between sites from simulated data were calculated using the mean of the coefficient of linkage disequilibrium *D* across all pairs of segregating sites:

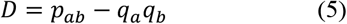

and the two normalized measures of LD:

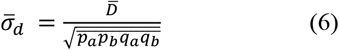

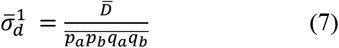

where *q*_*a*_ is the selected (deleterious) or minor allele frequency at the first site, *q*_*b*_ is the deleterious/minor allele frequency at the second site, *p*_*a*_ = 1 − *q*_*a*_, *p*_*b*_ = 1 − *q*_*b*_, and *p*_*ab*_ is the frequency of the haplotype “*a b*” which corresponds to the presence of either the deleterious or the minor alleles at both sites. The overbars represent the mean of the respective quantities over all SNP pairs and simulation replicates. Here, the mean of *D* and the mean of the product *p*_*a*_*p*_*b*_*q*_*a*_*q*_*b*_ was calculated across all SNPs for each simulated replicate. For each replicate, a single value of 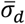 and 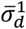 was obtained by taking the ratio of the mean *D* and the mean product of allele frequencies. The means and SEs reported in tables and figures are those across the 1000 replicate simulations. The fluctuations across replicates in the values of the three statistics (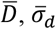 and 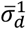) will thus be somewhat different in magnitude, so there is no guarantee that their overall means will have the same sign. However, this problem did not greatly affect our results, because it only happened when 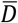 was not significantly different from 0.

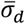 corresponds approximately to the mean correlation coefficient between the two loci (Ohta and Kimura 1969); the denominator of the statistic 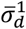, used by Good (2022), and Ragsdale (2022) greatly magnifies the apparent magnitude of LD compared with 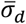. However, the denominator of 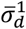 (the mean crossproduct of the four allele frequencies at a pair of segregating sites, rather than its square root) means that its range is not confined to (0,1), in contrast to the correlation coefficient.

Note that in our case, a *deleterious* allele may not be equivalent to a *derived* allele, since back mutations from – to + are possible. The distinction is important, since many empirical and theoretical studies of LD have relied on classifying variants as derived versus ancestral; neither of these are necessarily equivalent to *minor* alleles (for a discussion, see Potapova and Kondrashov 2023) We have used combinations of deleterious alleles as the basis for calculations of the direction of *D*; with sufficiently strong selection (*N*_*e*_*s* > 4), there will be little difference between these two classifications, since deleterious alleles are almost always derived. We also show simulation results for LD statistics where combinations of minor alleles were used as the criterion for positive LD, as has often been done in empirical studies.

### Two-locus simulations

We also investigated the effect of weighting LD statistics toward low frequency derived variants, a procedure suggested by Garcia and Lohmueller (2021) and Good (2022). We used the approach of Good (2022) for simulating a two-site haploid model without recombination. This procedure assumes that a mutation from A_1_ to A_2_ at the first site (A) is present at a frequency *q*, whose probability distribution is given by the standard formula for drift and selection (Equations S1.18 of section 6 of Supplementary File S1). A second mutation, from B_1_ to B_2_, was introduced at the second site, either in *cis* with A_1_ (case 1, probability 1 – *q*) or in *cis* with A_2_ (case 2, probability *q*). The fitness model for determining the post-selection frequencies of the haplotypes in a given generation assumed that the relative fitnesses of A_1_B_1_, A_1_B_2_, A_2_B_1_ and A_2_B_2_ are 1, 1 – *s*, 1 – *s* and 1 − 2*s*(1 + *ϵ*), respectively, where *ϵ* is a measure of epistasis of the same form as that used in the two-site diploid model. As described in section 6 of Supplementary File S1, multinomial sampling of *N* haplotypes was then carried out each generation, until one of the two mutations was fixed or lost, and the mean values over all generations and replicates of *D, D*^2^ and the allele frequency cross-product *P* = *p*_*A*_*q*_*A*_*p*_*B*_*q*_*B*_ (where subscripts *A* and *B* denote sites A and B) were obtained. The means of *D, D*^2^ and *P*, denoted by 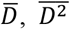 and 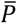, were found by dividing the respective sums of these statistics by the sum of the durations of the replicate simulations.

## Data availability

The scripts used to determine the properties of two-site equilibrium populations, perform all the simulations, and calculate statistics are provided at https://github.com/paruljohri/Gene_vs_sites_model/tree/main.

## Results

### Deterministic results for two selected sites

Here we examine the properties of equilibria under mutation-selection balance at two selected sites within a coding sequence. The fitness matrices in Table 1 are used for this purpose. The mutation rate from + to – at both sites is *u*. The symmetry of the fitness models means that the equilibrium allele frequencies are the same at both sites, so that the system can be completely described by the frequency *q* of the – allele at each site and the coefficient of linkage disequilibrium *D* between them. The correlation coefficient between the two selected sites is then given by *R* = *D*/*q*(1 − *q*). An algebraic treatment is given in sections 1-3 of the Appendix and sections 1-5 of Supplementary File S1. We focus on the case when the recombination frequency, *r*, is zero. This gives insights into the behavior of the system with low frequencies of recombination, such as those between pairs of sites within a gene. The complexity of the equilibrium equations, even when *r* = 0, means that exact results for the equilibrium haplotype frequencies can only be obtained numerically, as described in section 1 of the Appendix.

It is, however, possible to obtain approximate analytical results for the sign and magnitude of *D* and *R* at equilibrium without full solutions for the equilibrium haplotype frequencies. As might be expected from results in the earlier literature (e.g., Felsenstein 1965; Kimura and Maruyama 1966; Kondrashov 1988; Charlesworth 1990; Shnol and Kondrashov 1993; Roze 2021), a multiplicative version of the sites model with no synergistic or antagonistic epistasis results in equilibria with 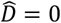 (where the carat indicates equilibrium). In contrast, the additive version results in equilibria with negative values of 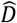, reflecting the weak synergistic epistasis on the multiplicative scale that is associated with additive fitnesses (see sections 2 and 3 of File S1). Explicit approximate expressions for the equilibrium values of 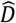 and 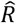 for the additive sites model with no epistasis and a low frequency of recombination are given by Equations (A9*a*), (A9*b*) and (A9*c*); the effect of epistasis on 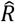 is described by Equation (A9*d*). Equations (A9) show that, if the ratio of recombination rate to selection coefficient (*r*/*s*) is 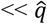 or << *h*, recombination has only a small effect on the magnitude of *D* and *R*.

In contrast, for the additive gene model with no synergistic epistasis, Equation (A10) shows that 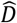 and 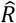 are positive when *h* is sufficiently small relative to ½ and *k* ≤ ½, selection is sufficiently weak, and 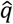 is sufficiently large. 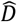 and 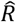 become slightly negative when these conditions are violated, even in the absence of synergistic epistasis, but are always positive for the corresponding multiplicative model when *h* ≤ *k* ≤ ½ (Equation A11). Equations (A10) and (A11) show that a smaller value of *k* is associated with larger LD values; in the absence of synergistic epistasis, the values of 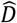 and 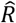 with *k* = *h* < ½ are approximately twice those with *h* < *k* = ½, reflecting the weaker selection against – – haplotypes in the former case (see Equation A3*b*). Unsurprisingly, with sufficiently strong synergistic epistasis, 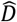 and 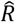 become negative (and vice-versa for antagonistic epistasis).

These conclusions are borne out by the results of iterating the exact recursion relations (Equations A1 and A2), which are shown in Figure 2. The upper panels in Figure 2 show the results for the sites model, and the lower panels those for the gene model with *k* = ½. In both cases, *u* = 0.0001 and *s* = 0.01. These are clearly not realistic values of the mutation and selection parameters but produce relatively large values of 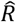, enabling the gene and sites models to be compared more convincingly than when 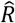 is very small. The left-hand panels plot 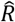 (in percent) against the epistasis coefficient, *ϵ*, for three different dominance coefficients, *h*. The right-hand panels plot 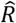 against *h* for three different values of *ϵ*. Since antagonistic epistasis (*ϵ* < 0) is well known to induce positive LD (Kondrashov 1988; Charlesworth 1990; Shnol and Kondrashov 1993), we have focused here on synergistic epistasis (*ϵ* > 0), as well as the zero epistasis case. As is shown below, the general properties of the deterministic case mirror those of the multi-site simulations, although the results are quantitatively very different.

**Figure 2.**
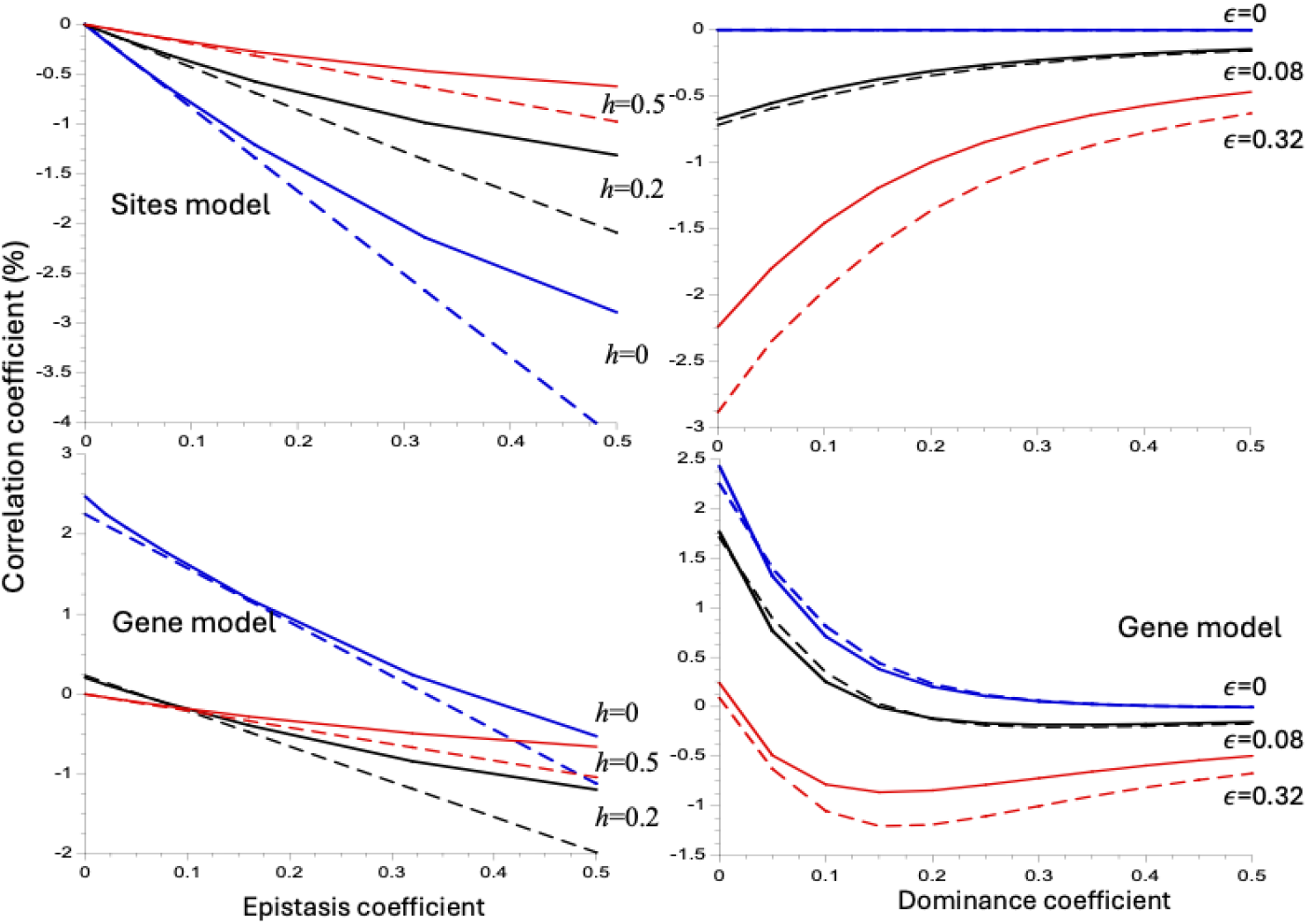
Linkage disequilibrium (LD) as a function of the synergistic epistasis coefficient (*ϵ*; left-hand panels) and dominance coefficient (*h*; right hand panels) for the two-sites deterministic equilibrium model, with *u* = 0.0001, *s* = 0.01, and *r* = 0. The upper panels are the results for the additive sites model; the lower panels are for the additive gene model with *k* = ½. LD is measured by the correlation coefficient (*R*) multiplied by 100. The solid curves are the exact solutions to the recursion equations for *D* (Equations S1.2 of Supplementary File S1). The dashed curves are the analytic approximations for *R;* see sections 2-3 of the Appendix, Equations (A.9) and (A.10). The horizontal lines in the lower panels indicate zero values of *R*.

Several points of interest emerge. As expected from the theoretical results, the sites model shows consistently negative but very small values of 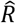, which are very close to the negative of one-half of the mutation rate (– 5 × 10^−5^) when *ϵ* = 0, independently of *h* (the difference from 0 is too small to be detectable in the figure, but can be seen in the results of the exact iterations provided in Supplementary File S2). The dominance coefficient has larger effects under the sites model, the larger *ϵ*; 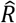 declines almost linearly with *ϵ*, becoming negative when *ϵ* is sufficiently large. In contrast, for the gene model with no epistasis, 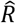 remains positive until *h* becomes close to ½, decreasing monotonically as *h* increases. With synergistic epistasis, the gene model is associated with larger values of 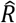 compared with the sites model when *h* < ½, and the difference is more marked, the smaller *h*, with small *h* being associated with larger magnitudes of 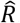. In addition, with the strongest value of epistasis (*ϵ* = 0.32), 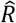 first declines with *h* and then increases again as it approaches ½ but remains negative. The approximations describe the general patterns quite well. In the absence of epistasis, 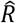 for the multiplicative gene model is always positive for *h* < 0.5, with a maximal value at *h* = 0, when the very small value of the denominator in Equation (A11) overcomes the fact that the numerator is proportional to 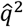 and is thus very small.

Sets of results for other values of *u* and *s* are shown in Supplementary File S2. The general features described above can be seen in these cases. However, as might be expected, the smaller the mean deleterious allele frequency, the smaller the differences between the gene and sites models, and the smaller the magnitudes of 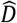 and 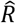. While 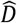 and 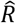 are still positive for the gene model in the absence of synergistic epistasis and sufficiently small *hs*, under the sites model they become negative at much smaller *ϵ* and 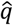 values. Thus, in order to determine whether there are any biologically important consequences of the features just described, it is necessary to determine what happens with multiple selected sites, where larger effects are to be expected.

### Deterministic results for the gene model with multiple sites

As described in section 4 of the Appendix, the results from the two-site gene model without epistasis can be extended to an arbitrary number of selected sites, all subject to the same selection regime, using the assumptions described in the above section *The gene model with multiple sites*. Equations (A19) provide approximate expressions for 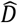 and 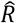 for the additive gene model without epistasis, given a value of 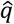, on the assumption that 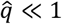 and that number of selected sites (*G*) is large. Equations (A21) show how to determine 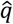 for given values of *G, u, s* and *h*. The results were derived for the case *k* = ½, since this was used in the simulations. The conclusion that positive LD between pairs of deleterious mutations is expected under the gene model, provided that *h* < ½, is confirmed by this analysis. In contrast to the two-site model, where 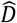 and 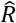 depend on 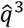 and 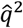 respectively (see Equation A11), 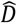 is proportional to 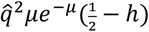 and 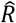 to 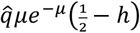 where 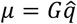 is the mean number of deleterious mutations per haplotype; this reflects the cumulative effects of LD across multiple sites. The next section presents comparisons of the predictions of Equations (A19) with LD statistics obtained in the simulations.

### Simulation results for multiple sites

The analytical results presented above were supplemented with forward simulations of a single gene with multiple selected sites, with an equal probability of mutations between the + and – alleles and vice versa (see the section *Methods, Multi-locus simulations*). We first describe the results with no synergistic or antagonistic epistasis.

#### Linkage disequilibrium with no epistasis

As discussed in the section *Deterministic results for two selected sites*, deleterious alleles are more likely to be associated on the same haplotypes in the gene model than in the sites model, resulting in larger positive mean *D* values for the gene model with suitable parameter values, as can be seen from the results in Figure 2. Table 2 shows the results of simulations without epistasis in which *s* was fixed, and LD statistics were calculated by assigning positive values to cases when + + and – – haplotypes were in excess of random expectation (entries under “selected allele”). In these examples, the gene model results in higher mean LD than the sites model for closely linked sites whenever there are significantly non-zero values. The difference between the two models increases with increasing recessivity and strength of selection, with mean positive values of 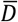 and 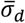 for the gene model versus negative values with the sites model when *h* < 0.5 and *γ* ≥ 20, except when *h* = 0 and *γ* = 100, when small but significantly positive values of 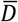 and 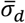 were found with the sites model, consistent with the results of Roze (2021) for small *h* values. For *γ* = 2, however, negative values were found with both models, consistent with the operation of Hill-Robertson interference (HRI) effects generated by the interaction between drift and selection with multiple linked selected sites (Hill and Robertson 1966; Garcia and Lohmueller 2021; Roze 2021), which are not allowed for in the two-site deterministic model.

**Table 2.**
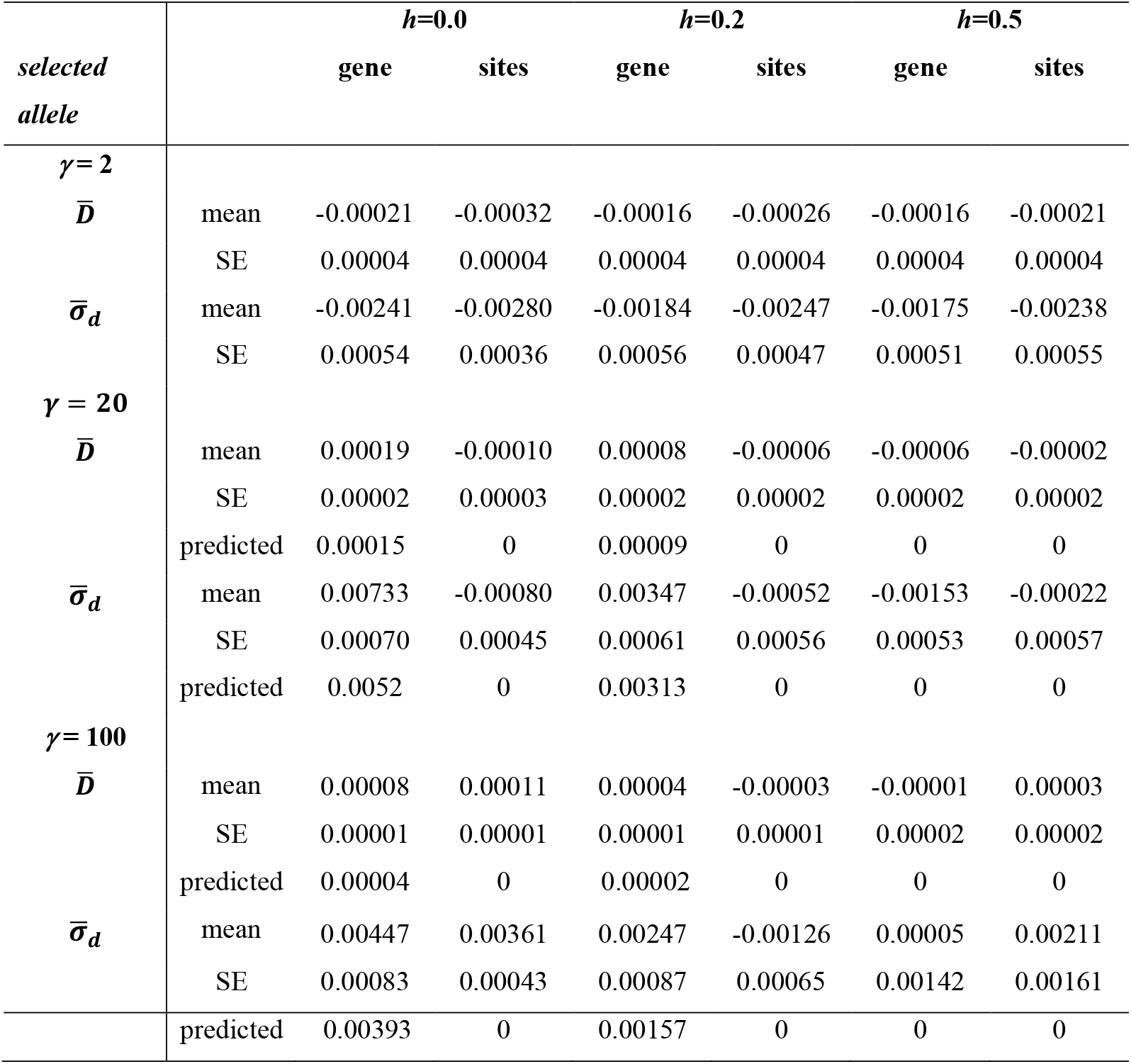
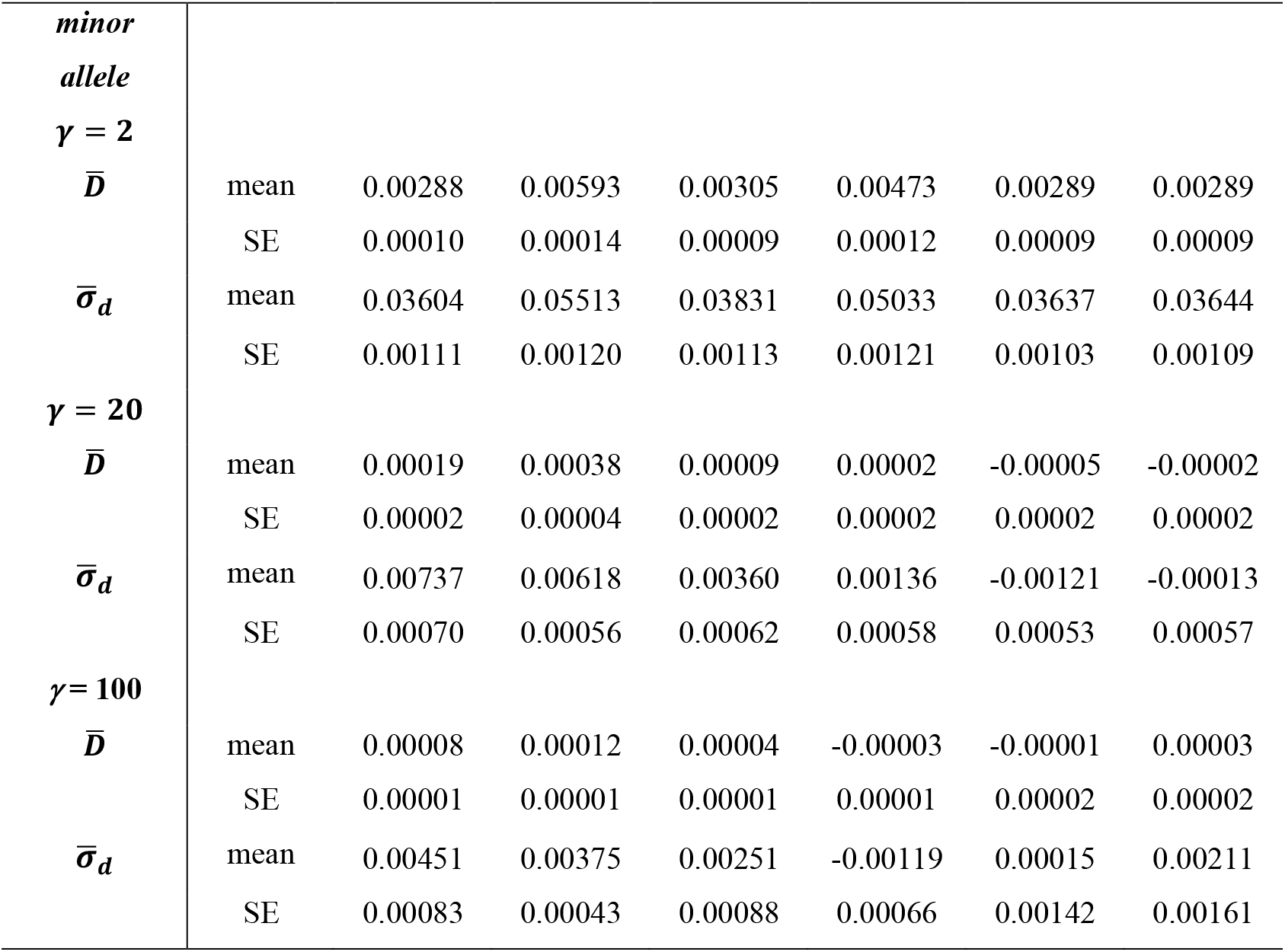
Statistical summaries of linkage disequilibrium between selected alleles for the gene and sites models, with fixed scaled selection coefficient *γ* = 2*Ns*. 1000 selected sites were simulated, with *Nu* = 0.005 and *Nr* = 0.01, where *N* is the population size (*N* = 1000), and *u* and *r* are the mutation and recombination rate per site/generation respectively. 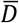 and 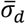 were calculated for alleles that were less than 100 sites apart. The sign of the LD statistic 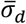 was assigned either by giving a positive sign to cases with excesses over random combinations of pairs of selectively deleterious alleles, or to cases with excesses of combinations of minor alleles. The means and standard errors (SEs) for 1000 replicate simulations are shown. The predicted values of 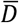 for the gene model with *γ* = 20 and 100 are also shown; these were obtained as described in the text; the corresponding values for the sites model are expected to be close to zero.

It is of interest to examine whether these results are consistent with the theoretical predictions for the multi-site gene model outlined in the previous section. The simulation results for *γ* = 20 and *γ* = 100 were used, since the results for *γ* = 2 must be heavily influenced by drift, whereas the theoretical predictions are purely deterministic. A complication in making this comparison is that the LD statistics for the simulations are presented for pairs of segregating sites, since *D* = 0 for invariant sites and 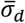 is defined only for segregating sites. As expected from general theory on strongly selected deleterious mutations (e.g. Charlesworth and Jain 2014), the proportion of sites that are segregating at a given time with *γ* = 20 or 100 is very small (see Table S5 in File S3).

The deterministic results for mean *D* are expected to correspond to the expected value of *D* over all pairs of sites. For this reason, the predictions for 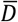 in Table 2 were obtained by dividing the predictions of 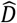 from Equation (A19*a*) by the squares of the mean proportions of segregating sites for the appropriate values of *γ* and *h*, obtained from Table S5. The values of 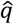 were obtained from Table 3 of Johri and Charlesworth (2025); these agree closely with the predictions from Equations (A21) (see Johri and Charlesworth (2025).

**Table 3:** Statistical summaries of linkage disequilibrium under the gene and sites models in human-like populations, using a gamma distribution of scaled selection coefficients with shape parameter 0.23 and mean 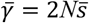. 2000 selected sites were simulated, with *Nu* = 0.00025 and *Nr* = 0.0002, where *N* is the population size (*N* = 1000), *u* is the mutation rate per site/generation, and *r* is the rate of crossing over between adjacent sites. The signs of 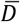 and 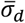 were assigned either by giving a positive sign to cases with excesses over random combinations of pairs of selectively deleterious alleles, or to cases with excesses of combinations of minor alleles. The means and standard errors (SEs) for 1000 replicate simulations are shown.

| *selected*<br>*alleles* | *h=0.0* |  |  | *h=0.2* |  | *h=0.5* |  |
| --- | --- | --- | --- | --- | --- | --- | --- |
|  |  | gene | sites | gene | sites | gene | sites |
| $\bar{\gamma} = 850$ | | | | | | | |
| $\bar{D}$ | mean | 0.00015 | 0.00031 | 0.00101 | 0.00030 | -0.00096 | -0.00105 |
|  | SE | 0.00045 | 0.00035 | 0.00062 | 0.00066 | 0.00063 | 0.00074 |
| $\bar{\sigma}_d$ | mean | 0.01130 | 0.00650 | 0.01587 | 0.00345 | -0.00706 | -0.00345 |
|  | SE | 0.00472 | 0.00353 | 0.00591 | 0.00547 | 0.00525 | 0.00600 |
| $\bar{\sigma}_d^1$ | mean | 0.46777 | 0.25534 | 0.61505 | 0.14143 | -0.12457 | 0.20649 |
|  | SE | 0.17992 | 0.08900 | 0.21642 | 0.11249 | 0.09229 | 0.20202 |
| *minor*<br>*alleles* |  |  |  |  |  |  |  |
| $\bar{\gamma} = 850$ | | | | | | | |
| $\bar{D}$ | mean | 0.00123 | 0.00137 | 0.00319 | 0.00319 | 0.00244 | 0.00285 |
|  | SE | 0.00046 | 0.00036 | 0.00063 | 0.00065 | 0.00071 | 0.00078 |
| $\bar{\sigma}_d$ | mean | 0.01924 | 0.02183 | 0.03972 | 0.03556 | 0.02356 | 0.03175 |
|  | SE | 0.00473 | 0.00373 | 0.00601 | 0.00548 | 0.00560 | 0.00625 |
| $\bar{\sigma}_d^1$ | mean | 0.54411 | 0.51751 | 0.94606 | 0.55386 | 0.20582 | 0.60968 |
|  | SE | 0.17966 | 0.09122 | 0.21699 | 0.11278 | 0.09449 | 0.20269 |

The predicted value of 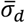 for a given parameter set can be obtained by dividing the predicted value of 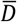 by one half of the nucleotide site diversity at selected sites (*π*), normalized by the proportion of segregating sites. The values of *π* for the parameters in Table 2 were obtained as described by Jain and Charlesworth (2014), equating the selection coefficient in their formulae to *s*[1 + (*μ* − 1)*e*^−*μ*^(1 − 2*h*)] in Equation (A20*b*). Overall, the agreement between observed and predicted values is remarkably good, considering the approximations involved, suggesting that the multi-site gene model provides an accurate description of the population genetics of LD when selection is strong and *G* is large.

Similar results to those in Table 2 were found with a gamma distribution of selection coefficients, with the gene model yielding positive values of 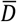 and 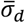 with *h* < 0.5 and 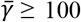, as opposed to slightly negative (but not significantly non-zero) values for the sites model (Figure 3 and Figure S1 in Supplementary File S3). The theoretical results for the two-sites model predict that 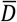 and 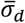 values are expected to be slightly negative for the sites model. Consistent with this prediction, we observed small negative values when *h* < ½. These effects could also be caused by HRI.

**Figure 3:**
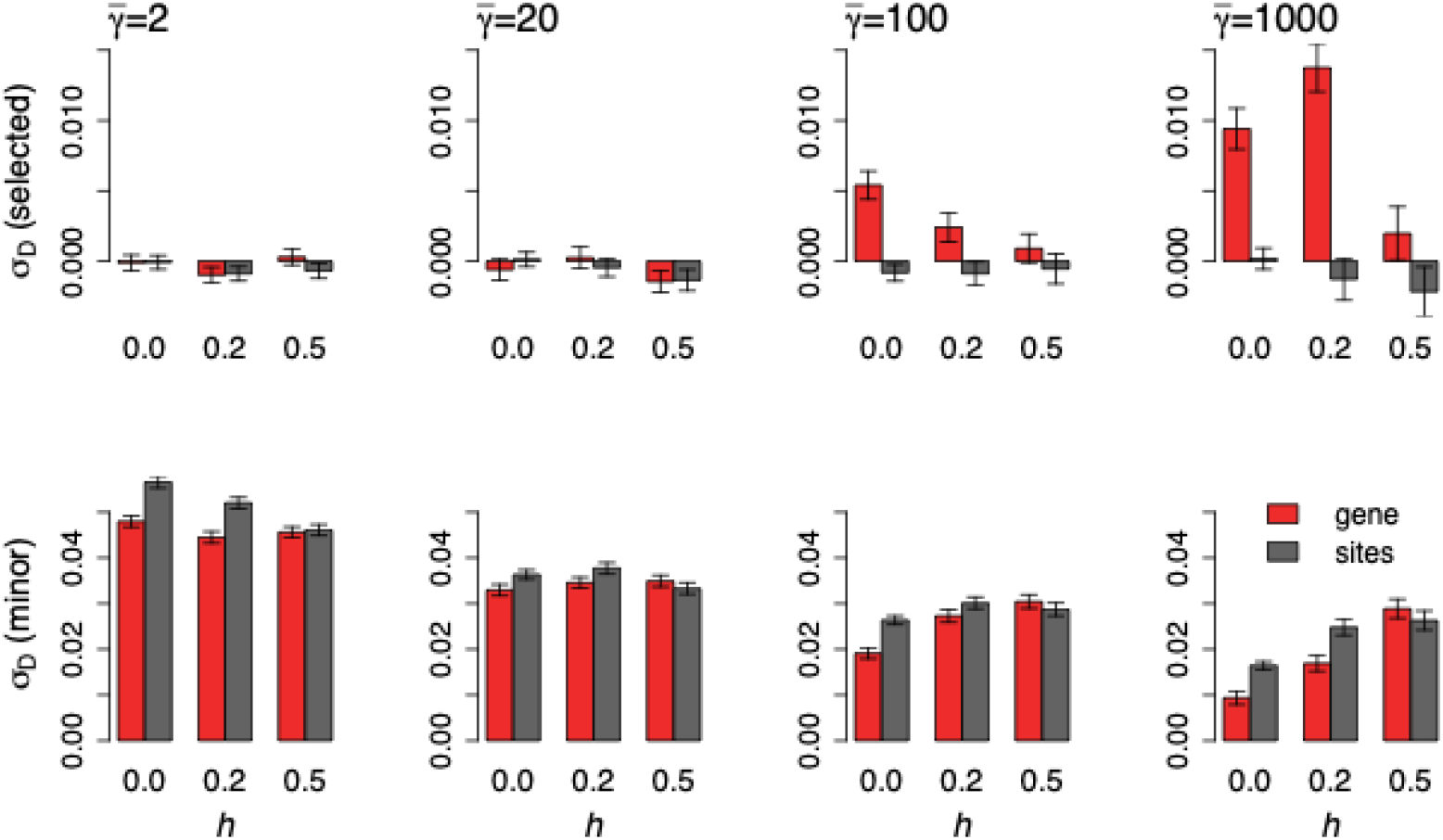
Statistical summaries of linkage disequilibrium as measured by 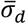 under the gene and sites models for a gamma distribution of scaled selection coefficients with shape parameter 0.3 and a mean of 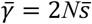. 1000 selected sites were simulated, with *Nu* = 0.005 and *Nr* = 0.01, where *N* is the population size (*N* = 1000), *u* is the mutation rate per site/generation, and *r* is the rate of crossing over between adjacent sites. 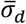 was calculated for alleles that were less than 100 basepairs apart. The sign of 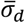 was assigned either by giving a positive sign to cases with excesses over random combinations of pairs of selectively deleterious alleles, or to cases with excesses of combinations of minor alleles (see the Methods section). The means and standard errors (SEs) for 1000 replicate simulations are shown. Similar results for 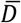 are shown in Figure S1 of Supplementary File S3.

To test this possibility, we analyzed the results with a DFE for variants separated by a greater-than-average distance (800-100 base pairs; Table S1 in Supplementary File S3). The 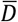 values for selected variants under the sites model then approach zero, consistent with reduced effects of HRI with higher frequencies of recombination, whereas the deterministic theory suggests only a weak effect of recombination with the low values of *r*/*s* for pairs of sites within a gene (see the previous section *Deterministic results for two selected sites*). In contrast, for the gene model under a gamma distribution with 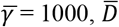 and 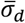 are still substantially greater than zero, suggesting significantly positive LD even at larger distances (Table S1 in Supplementary File S3). In addition, the magnitude of negative LD is greatest with *h* = 0.5, whereas the deterministic sites model predicts zero LD (see Figure 2).

Furthermore, as discussed in the previous section, the multiplicative fitness sites model is not expected to generate any LD, in contrast to the additive model. When fitnesses were assigned to be multiplicative in our multi-locus simulations, 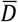 and 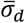 values were sometimes significantly negative when sites were less than 100 bases apart, especially with *h* = 0 (Table S2 in Supplementary File S3). For alleles between 800-1000 bases apart, 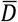 and 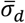 values were close to zero in all cases (Table S2 in Supplementary File S3). This suggests that HRI was probably partly responsible for the slightly negative values observed in our multi-site simulations of the sites model.

Interestingly, when an excess frequency of haplotypes with pairs of minor alleles was used as the criterion for positive *D*, there is not much difference in 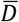 and 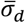 between the gene and sites models when selection is weak, with mainly positive values for both (Table 2 and Figure 3). The sign of 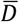 is expected to be positive for minor alleles in cases where the overall value of 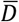 for derived alleles is zero. This comes about because there is a reasonable chance that, when derived variants in a pairwise comparison on different haplotypes, one of the derived alleles at one site in a pairwise comparison is present at a frequency greater than 0.5 and is regarded as a major allele while the derived allele at the other site is regarded as a minor allele, so that *D* for the minor allele combination is positive. Taking the average over all possible situations, this results in a bias toward positive LD among minor alleles (Potapova and Kondrashov 2023). However, with stronger selection against – alleles and *h* < ½, the gene model yields slightly lower values of 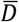 and 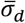, possibly due to the lower allele frequencies under the gene model compared with the sites model (see Johri and Charlesworth 2025).

Our forward-in-time simulations were conducted with small populations (1000 diploid individuals) that are a factor of 1000 smaller than the value of *N*_*e*_ commonly used for *Drosophila* populations, so that the mutation rate, recombination rate, and selection coefficients were a factor of 1000 larger than the natural population values (see the Methods section). To check if rescaling biased our results, we conducted simulations with a much larger population size (5000 individuals). LD statistics obtained using minor allele frequencies were entirely unaffected by rescaling (Table S3 in Supplementary File S3). rescaled population. The differences between the two *N* values in the means of the LD statistics for pairs of deleterious variants were within the limits of sampling error, so that there is no evidence that rescaling has a major systematic effect on these statistics.

In contrast to the simulations with *Drosophila*-like parameters, which we have discussed above, simulations of human-like populations yielded high variances between replicates and somewhat unclear patterns for the LD statistics (Table 3). Overall, however, LD between selected sites is higher with the gene model than the sites model when *h* < ½. In this case, we show 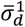 as well as 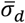, for comparison with the analyses of human data by Ragsdale (2022) (see *Discussion, Patterns of linkage disequilibrium in population genetic data*). The difference in LD between the two models is much clearer when using 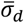 or 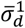 as opposed to 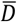, because 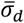 and 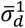 (partially) correct for differences in allele frequencies.

#### LD with epistasis

Similarly to the analytical results discussed earlier, there is a substantial decrease in 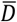 and 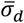 (using the selected/derived allele criterion) with increasing strength of synergistic epistasis for both models when *h* is close to 0.0, but only small changes when *h* is 0.2 or 0.5 (Figures 4 and Figure S2 in Supplementary File S3). 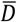 and 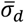 between selected alleles remain significantly higher with the gene model than the sites model (for which both statistics become negative) with large *ϵ*. There seems to be no significant difference between the gene and sites model with these parameters when *h* = 0.2 and *ϵ* is very small, consistent with their large SEs relative to the means when 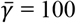 (see Figure 3 for the results with no epistasis, and Figure S3 in Supplementary File S3 for a comparison of the distribution of the statistics for the results in Figure 3 and Figure 4); 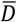 and 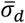 are both not significantly different from zero in this case. Importantly, however, with *h* = 0.2 and 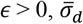 is negative with the sites model, but is close to zero with the gene model, so that the two models display differences in LD even with epistasis.

**Figure 4:**
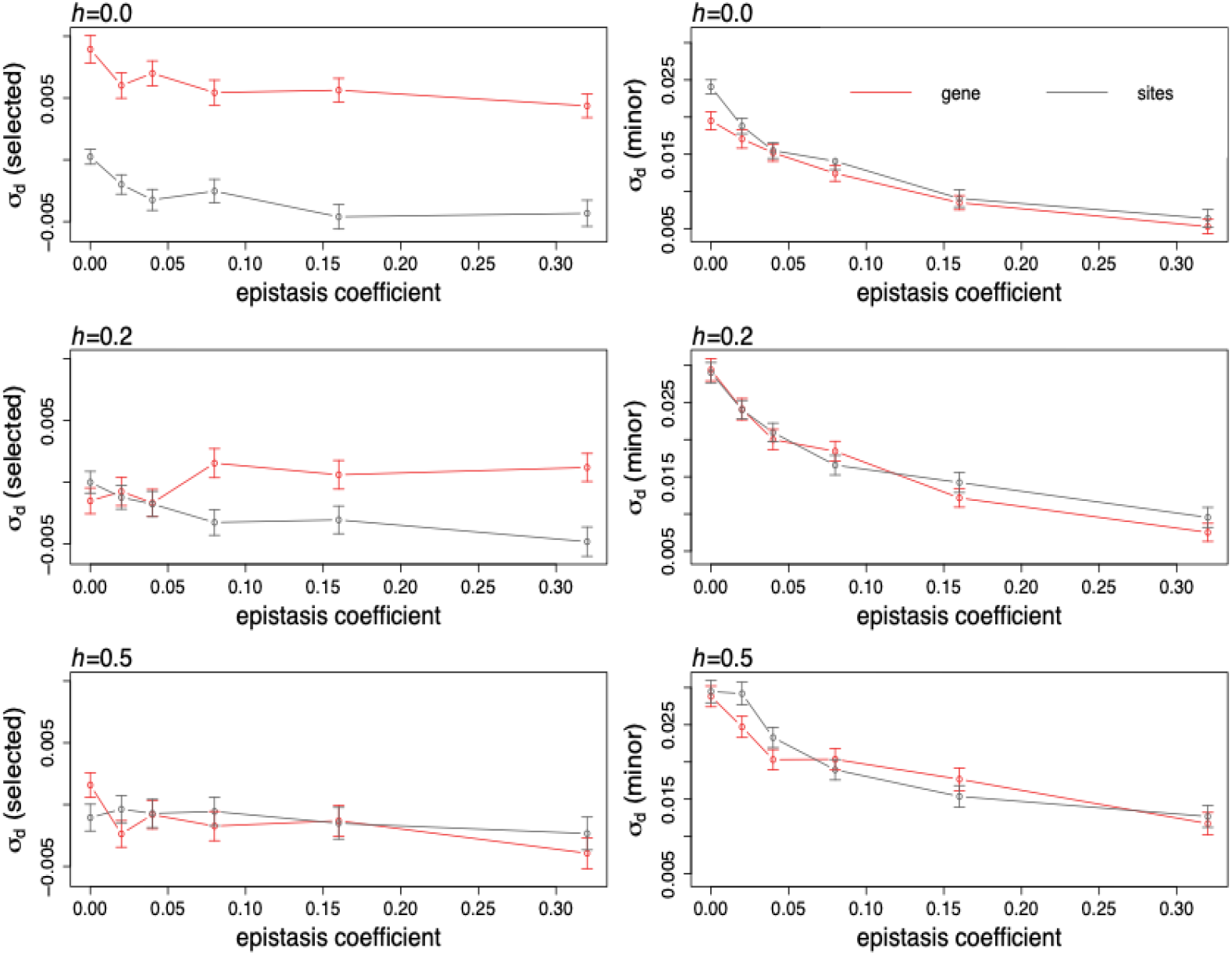
Statistical summaries of the 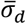 measure of linkage disequilibrium between all pairs of sites segregating for deleterious mutations for gene and sites model with epistasis, for closely linked pairs of alleles (1-100 basepairs apart). The simulation parameters were the same as in Figure 3. The sign of the LD statistic 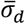 was assigned either by giving a positive sign to cases with excesses over random combinations of pairs of selectively deleterious alleles or to cases with excesses of combinations of minor alleles. The error bars represent the SEs across replicates. Similar results for 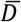 are shown in Figure S2 of Supplementary File S3.

When the same LD statistics are calculated with minor/major alleles as the sign criterion, there is a substantial decrease in LD with an increase in the epistasis coefficient; however, the values always remain positive and there is no significant difference in 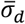 between the two fitness models (Figure 4), although 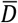 obtained using minor alleles is consistently and slightly smaller for the gene than the sites model, possibly due to lower mean allele frequencies at equilibrium (see Table 3 of Johri and Charlesworth 2025). In summary, if synergistic epistasis were prevalent amongst deleterious alleles, LD between selected/derived alleles will probably still be higher and positive with the gene model than the sites model when *h* < ½, whereas no difference is expected between LD statistics calculated using minor alleles as the sign criterion.

With much lower rates of crossing over (0.1 × the standard rate used here), when mutations are fully recessive, LD calculated using minor alleles is significantly higher with the sites model than the gene model (Figures S4 and S5 in Supplementary File S3). The difference between the two models is largest when there is no epistasis, but persists even when the epistasis coefficient is 0.32. The predicted differences in LD between the gene and the sites models using the derived and minor allele sign criteria can thus be entirely opposite, depending on the degree of dominance. HRI in the absence of synergistic epistasis seems to produce only a weak tendency towards negative LD when *h* = ½, and for the sites but not the gene model when *h* = 0 or 0.2.

### Effects on LD statistics of weighting towards low allele frequencies

We now consider the potential effects of preferentially using rare variants on estimates of LD, a procedure proposed by Garcia and Lohmueller (2021) and Good (2022). Use of such a procedure could produce substantially different patterns from those described above. Garcia and Lohmueller (2021) suggested that the effects of HRI on LD between deleterious mutations would be more easily detected in samples from populations if doubleton variants were used for estimating LD. This procedure might help to correct for the effect of allele frequency on LD statistics; in addition, because rare variants are more likely to be deleterious than common ones, larger negative *D* values might be expected for nonsynonymous than synonymous variants when conditioning on doubletons. In their multi-locus simulations, they indeed found negative LD between derived variants at sites subject to purifying selection, both with *h* = 0 and *h* = ½, with LD being measured by Lewontin’s *D’* statistic (Lewontin 1964). In contrast, Good (2022) proposed that multiplying allele frequencies by a function that gives greater weights to low frequency variants should lead to positive expected *D* values, for both neutral and deleterious variants. He validated this expectation by a combination of simulations and analytical formulae, using a model of a haploid population; this is equivalent to a diploid population with *h* = ½ in the absence of epistasis. However, while the simulations performed by Garcia and Lohmueller (2021) involved multiple selected sites, those of Good (2022) involved only two sites.

In order to reconcile these seemingly contradictory findings, we used the approach described in the section *Methods: Two-locus simulations* and section 6 of Supplementary File S1. This enables the means of *D, D*^2^ and the crossproduct *P* of the four allele frequencies at sites segregating for two mutations (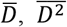 and 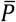) to be found, yielding estimates of the standardized LD statistics 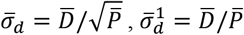 and 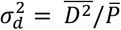. 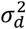 corresponds approximately to the mean squared correlation coefficient between the two loci (Ohta and Kimura 1969); the other two statistics are defined by Equations (6) and (7). These statistics were estimated either with or without weighting. The weighting factor *f* is 0 if no weighting is applied, and *f* > 1 if low frequency mutant alleles receive preferential weights (*f* is the inverse of the factor used by Good (2002)).

For neutral variants, the unweighted case gave values of 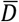 and 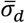 that were non-significantly negative and close to zero: 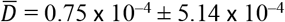 and 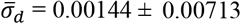. The value of 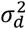 was close to the theoretical value of 0.45 for the case of no recombination under the infinite sites model (Ohta and Kimura 1971). In contrast, 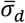 and 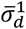 with weighting (*f* = 10)were 0.0107 and 0.915, respectively. The values when doubletons were sampled were 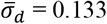 and 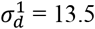.

Some representative results for 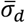 with selection are shown in Figure 5, for two different strengths of selection (*γ* = 2*Ns* = 10 and *γ* = 2*Ns* = 50) and varying amounts of synergistic epistasis. With no epistasis and no weighting, 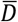 and 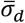 were slightly negative, but became increasingly negative as *ϵ* increased. The case of *γ* = 50 without epistasis gave no evidence for significantly non-zero LD 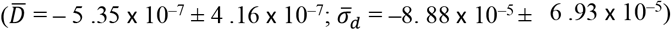, implying that HRI effects with two sites are not sufficiently strong to overcome any tendency for positive LD to be produced by purifying selection. Weighting causes the LD statistics to take larger values, which are positive with large weighting factors towards rare alleles (*f* = 100; see the details in section 7 of Supplementary File S1) even for the largest value of *ϵ* (0.64). They remain positive over the whole range of *ϵ* if doubletons are sampled. Further results are shown in Supplementary Files S4 and S5.

**Figure 5.**
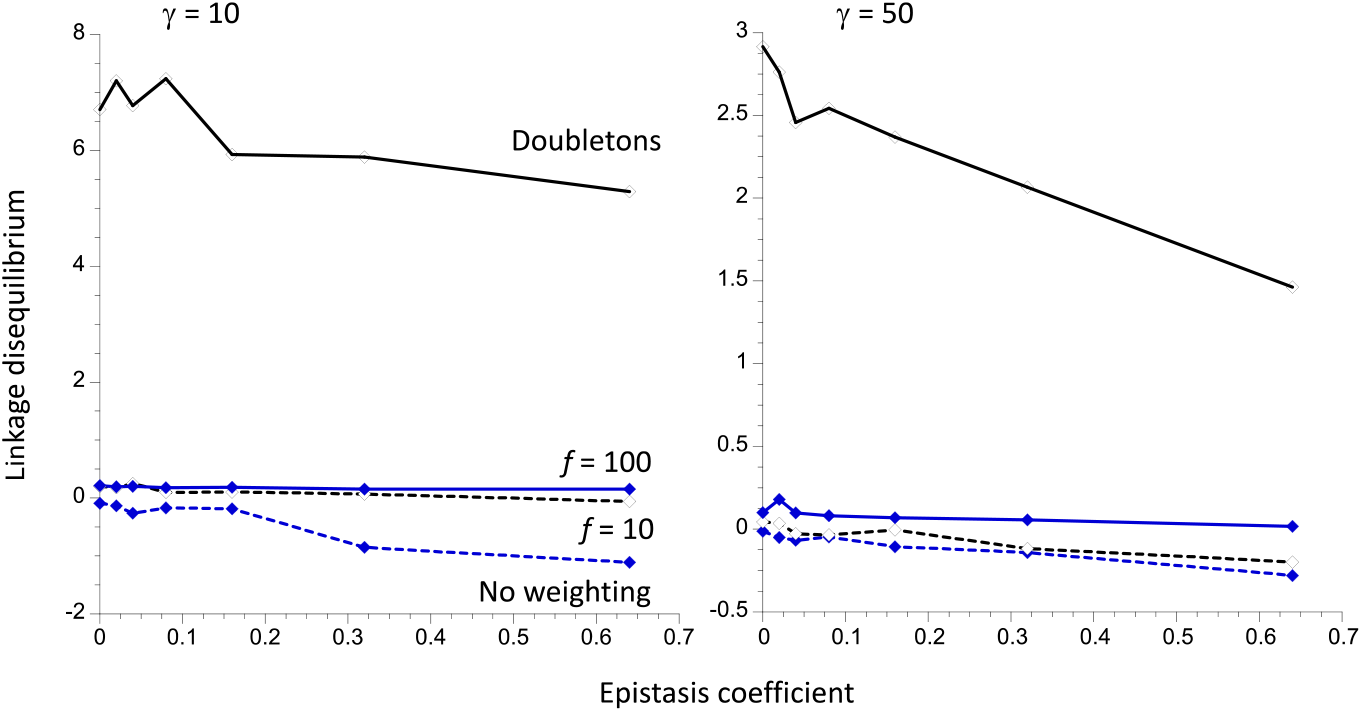
Mean linkage disequilibrium (LD) as a function of the epistasis coefficient (*ϵ*), from stochastic simulations of the two-site haploid model with *r* = 0. LD is measured by the 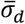 statistic described in the text (multiplied by 100). The results for 10^6^ replicate simulations with two different values of *γ* = 2*Ns* are shown, with *N* = 1000. The dashed curve with filled markers shows the population values of LD with no weighting towards low allele frequencies; the full curve with open markers is for doubleton variants in a sample of size 100. The dashed curve with open markers shows population values with a weight of 10 toward low frequency variants; the full curve with filled markers is for a weight of 100. See the main text for further details.

These results show that focussing on mutations at low frequencies results in positive LD between derived variants, which can persist even if there is strong synergistic epistasis, especially if doubletons are sampled. A heuristic argument that explains this effect is provided in section 7 of Supplementary File S1. These considerations suggest that the use of LD statistics estimated from low frequency variants is not helpful in diagnosing HRI effects caused by purifying selection. On the contrary, it is likely to conceal the effects of HRI or synergistic epistasis on LD between derived variants.

This raises the question of why Garcia and Lohmueller (2021) found consistently negative LD in their multi-locus simulations of deleterious mutations when sampling doubletons. It turns out that this was because they applied the *D*’ statistic (Lewontin 1964) to estimates of *D* for derived variants. *D*’ is defined as *D*/|*D*_*max*_|, where |*D*_*max*_| is the maximum value of the absolute value of *D* conditioned on the allele frequencies. The basis for this conclusion is described in section 5 of the Appendix and section 8 of Supplementary File S1, where it is shown that purifying selection can cause *D’* at selected sites to be lower than at neutral sites, reaching negative values even if their frequencies are similar and if doubletons are used.

To avoid these sources of bias, one should use unweighted values of *D* or 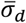 and avoid using *D*’. It is, however, currently unclear how nonequilibrium scenarios like population size changes might affect the signs of these statistics and whether unweighted values of *D* would be the best to use in all cases.

## Discussion

### The role of complementation

A major finding of this work is the effect on LD statistics of the distinction between the gene and sites models, which we discuss in detail below. In the accompanying paper (Johri and Charlesworth 2025), we discuss the implications for population genetics statistics relevant to the genetic load and inbreeding load caused by deleterious mutations. Our results raise the question of how widely the gene model might apply to real organisms. The distinction between the two models is based on the well-established finding of classical and molecular genetics that two recessive loss-of-function mutations affecting the amino-acid sequence of the same polypeptide usually fail to complement each other when present in the same cell in *trans* in a double heterozygote, due to the fact that both copies of the polypeptide are non-functional. In contrast, the same mutations present in *cis* in a double heterozygote result in the cell containing one functional and one non-functional polypeptide, with near wild-type performance (see Figure 1). This finding is of great historical importance, since it underlies the recognition of the gene as a coding sequence whose sites could be separated by recombination, and to Seymour Benzer’s concept of the cistron as the functional genetic unit (reviewed by Benzer 1957; Pontercorvo 1958; Hawley and Gilliland 2006).

If mutations within the same coding sequence fail to complement each other when in *trans*, the assumptions of the gene model are satisfied. The sites model must usually apply to mutations affecting different coding sequences, which necessarily complement each other unless there are unusual types of epistasis between them, e.g., due to interactions between mutations in different members of a multi-protein complex. Examples of such a failure of complementation between non-allelic loss-of-function mutations are reviewed by Hawley and Gilliland (2006) and Navarro-Quiles et al. (2024); these appear to be very rare compared with complementation.

It should be noted, however, that there are exceptions to a lack of allelic complementation (reviewed by Fincham et al. 1979, Chap. 4; Hawley and Gilliland 2006). In such cases, loss-of-function mutations affecting the same polypeptide complement each other, so that *trans* double heterozygotes show wild-type or near wild-type performance. In many cases, some mutations within the same gene complement each other, whereas others do not, allowing the construction of complementation maps (Fincham et al. 1979, Chap. 4). Some mutations within the same gene may thus obey the gene model and others the sites model. Allelic complementation has been best studied in fungi like *Neurospora*, and often involves cases where multimers of the same polypeptide form part of the same protein molecule (Fincham et al. 1979, pp. 348-353). In other cases, spatial interactions between mutations in distinct functional subunits of the same polypeptide are involved (Fincham et al. 1979, p. 352; Eng et al. 2015). It is often found that interallelic complementation is incomplete, when phenotypes such as enzyme activity are measured (Fincham et al. 1979, pp. 348-351), so that the distinction between the two models is somewhat blurred. Similarly, while Chandler et al. (2017) found multiple instances of hypomorphic alleles that complement each other phenotypically, the extent of complementation between two alleles was dependent on the genetic background. For simplicity, we have not attempted to include any of these complexities in our models.

It is, of course, less clear what happens to the heterozygous fitness effects of more minor effect mutations in the same coding sequence when configured in *cis* or *trans*, and there appears to be little relevant data. The simplest assumption is that the effects parallel those of loss-of-function mutations and are recessive or partially recessive, but are additive in their effects when in *trans*, as we have assumed here when setting *k* in Table 1 to ½. There are, of course, many other possibilities, including the extreme alternative of setting *k* = *h* to represent the fitness effect of a heterozygous effect of a mutation at one site on a background of homozygosity for a mutation at another site.

The “gene-based model” of Thornton et al. (2013) and Sanjak et al. (2015) is similar to ours, in that it assumes that a *trans* heterozygote with *n* mutations in a gene has the same fitness as a homozygote with *n* mutations. They did not, however, consider the position effect, *i*.*e*, the effects of the difference in fitness between *trans* and *cis* heterozygotes. Clark (1998) evaluated the effects of allelic complementation or non-complementation between pairs of deleterious alleles on allele frequencies at a multiallelic locus, but did not examine patterns of LD. Ragsdale (2022, Table S1) describes two versions of a two-site model. His “simple dominance” model is identical with our sites model (allowing for the difference in notation). His “gene-based” model is similar to our gene model, in that it assigns different fitnesses to *cis* and *trans* double heterozygotes. However, it differs in several other respects, e.g., by assigning the same fitness to single homozygotes, single heterozygotes, double homozygotes, and *cis* double heterozygotes.

The situation with respect to mutations affecting the same non-coding regulatory sequence such as a UTR, an enhancer or an insulator is somewhat unclear, although it seems likely that mutations within the same cis-regulatory sequence should show similar *cis*/*trans* behavior to mutations in the same polypeptide, fitting the gene model, while mutations in different cis-regulators of the same gene should complement each other, and hence fit the sites model. The phenomenon of transvection involves complementation between mutations in different cis-regulatory sequences of the same gene when the two homologous chromosomes or regions of homologs are paired in a cell nucleus, but failure of complementation when they are unpaired (Lewis 1954; Duncan 2002; Mellert and Truman 2012; Galouzis and Prud’homme 2021); the normal situation in this case is for them to fit the sites model. However, a recent study of hypomorphic regulatory mutations in two *D. melanogaster* genes showed a predominance of failure to complement (Chandler et al. 2017).

### The role of epistasis

The extent to which deleterious mutations with minor effects on fitness have epistatic effects rather than obeying an additive or multiplicative model for their combined effects, and the typical nature of such interactions, is still an unsettled issue with respect to both theoretical (Diaz-Colunga et al. 2023) and empirical (Bakerlee et al. 2022) questions. Attempts to characterize epistasis for fitness components among deleterious mutations in a variety of organisms have used analyses of the change in the means over generations of mutation accumulation lines or populations subject to inbreeding. These experiments have yielded somewhat conflicting results, although overall there seems to be a predominance of weak synergistic epistasis (reviewed by Charlesworth 1998; Fry 2004; de Visser et al. 2011; García Dominguez et al. 2019). A problem with interpreting these results is that data on fitness, or its components such as viability or fecundity, are inherently noisy and there may be statistical biases (due to regression towards the mean) that lead to an underestimation of synergistic epistasis among deleterious mutations (Berger and Postma 2014).

Microbial systems have allowed estimates of the fitness consequences of combining deletions of two or more genes, providing extensive information about epistasis among drastically deleterious mutations, using haploid genomes (e.g., Costanzo et al. 2016; Kuzmin et al. 2018). New technologies that allow the introduction of multiple mutations into the same coding sequence have opened a path towards characterizing epistasis between mutations at the intragenic level. This can be done either at the level of a specific biochemical function, such as resistance to an antibiotic (e.g., Bershtein et al. 2006), fluorescence intensity of the green fluorescent protein (e.g., Sarkisyan et al. 2016; Somermeyer et al. 2022), or by measurements of the fitness effects of mutant strains in microbial competition experiments (e.g., Bank et al. 2015, 2016; Puchta et al. 2016; Domingo et al. 2018; Bakerlee et al. 2022; Chen et al. 2023; Johnson et al. 2023; Papkou et al. 2023). Interestingly, although the stability of proteins is nearly linearly reduced by increasing numbers of mutations (De Pristo et al. 2005; Bershtein et al. 2006), the effect of mutations on function or fitness in the experiments just cited seems to show a predominance of synergistic epistasis among deleterious mutations, despite complexities such as the dependence of patterns of interaction between the same mutations in different environments (Bank et al. 2016) or between-species differences in fitness landscapes (Somermeyer et al. 2022).

There appear to be no reliable estimates of the parameters involved in quantifying epistasis (such as our *ϵ*) in organisms such as *Drosophila* or humans, making it difficult to evaluate the relevance of our model of epistasis to patterns of LD in natural populations. It is, however, possible that synergistic epistasis among deleterious mutations could reduce the positive LD expected under the gene model to near zero, unless mutations are highly recessive (see Figure 4). The extent to which the sign of LD between deleterious variants is affected by epistasis versus HRI also remains an open question, but our simulations suggest that synergistic epistasis is more likely to produce detectable negative LD than HRI in genomic regions where recombination rates are not unusually low.

### Linkage disequilibrium: theory

It has generally been accepted that multiplicative fitness models of multi-locus mutation and selection in randomly mating, discrete generation populations predict an absence of linkage disequilibrium when equilibrium has been reached in an infinite population (Haldane 1937; Kimura and Maruyama 1966). This is confirmed by our analytical results for the two-site version of the sites model with multiplicative fitnesses (section 2 of Supplementary File S1). Previous theoretical studies have found that synergistic epistasis on a multiplicative scale results in negative LD when a positive sign of *D* is assigned to haplotypes with pairs of mutant or wild-type alleles, whereas antagonistic epistasis causes positive LD (Kondrashov 1982, 1988; Charlesworth 1990; Shnol and Kondrashov 1993). Furthermore, in discrete generation finite populations, there is a general expectation that negative LD will be generated by Hill-Robertson interference in the absence of epistasis on the multiplicative fitness scale, provided that linkage is sufficiently tight and mutations are not highly recessive (Hill and Robertson 1966; Felsenstein 1974; Garcia and Lohmueller 2021; Roze 2021), although the opposite pattern can arise with sufficiently small values of *h* (Roze 2021; Ragsdale 2002).

Since an absence of epistasis on a linear scale produces weakly synergistic epistasis on a multiplicative scale, it might be expected that our deterministic two-site model would result in small negative values of the two measures of LD, *D* and *R*, when the epistatic parameter *ϵ* is equal to zero. The analytical results presented here confirm this expectation for the sites model (see section 2 of the Appendix and Supplementary File S1, sections 2 and 3, but the properties of the gene model are significantly different (section 3 of the Appendix and section 4 of File S1). Both the analytical results and results of exact iterations (Supplementary File S2) show that the additive sites model exhibits negative values of *R* that are very close to one-half the mutation rate when *ϵ* = 0, and much larger negative values when *ϵ* > 0.

In contrast, the correlation coefficient *R* is often positive under the additive gene model when *h* < ½, for sufficiently small values of *ϵ*. This reflects the fact that the higher fitness of *cis* compared with *trans* double heterozygotes (see Table 1) produces an excess of + + and – – haplotypes compared with + – and – + haplotypes, as was mentioned in the Introduction. However, the fitness of the – –/+ – and – –/– + genotypes, 1– (1 + *k*)*s* in the absence of epistasis, also influences the magnitude of LD, through the contribution of these genotypes to the marginal fitness of the – – haplotype; a reduction in this marginal fitness causes a reduction in the frequency of the – – haplotype, diminishing the value of *D* and *R*. For this reason, our numerical results that assume *h* ≤ *k* = ½ are conservative with respect to the magnitude of positive LD among pairs of deleterious mutations. In the absence of a *cis*/*trans* effect, a modification of the sites model, with the fitnesses of – –/+ – and – –/– + becoming 1 – (1 + *k*)*s* rather than 1 – (1 + *h*)*s*, results in negative LD with *h* < *k* ≤ ½ (see section 5 of Supplementary File S1). This model generates positive LD only under the biologically implausible condition *k* < *h*, which would imply that a homozygote for – has a *higher* fitness than a heterozygote. This finding emphasizes the fact that positive LD under the deterministic two-site gene model arises from the *cis*/*trans* effect, unless epistasis is antagonistic rather than synergistic.

Similarly, the finding of positive LD in simulations of the gene model without epistasis, at least with *h* < ½ and sufficiently strong selection (see *Simulation results for multiple sites*; *linkage disequilibrium with no epistasis*), makes sense given the intragenic position effect on fitness. The results in Table 2 show that the approximations for LD derived in section 4 of the Appendix for the multi-site gene model (Equations A19 and A21) agree well with the simulation results with *k* = ½ when the selection regime is the same for all sites and selection is strong relative to drift, so that this approach provides useful insights into when to expect positive LD between variants within the same gene.

In the simulations of multiple mutations within the same gene, diploid genotypes with *cis* and *trans* configurations of mutations are likely to be relatively abundant in a randomly mating population, unless selection is very intense. For example, with a scaled selection coefficient of *γ* = 20 and *h* = 0.2, the mean frequency of a deleterious mutation at a given site under the gene model with *k* = ½ is approximately 0.001 (see Table 3 of Johri and Charlesworth 2025). The mean number of mutations per haplotype with 1000 selected sites in a coding sequence is thus approximately 1. Since mean *D* values are small in this case (Table 2), a Poisson distribution should give a good approximation to the distribution of the number of mutations per haplotype. This implies a frequency of approximately 0.13 of individuals with two mutations in *cis* over a wild-type haplotype, 0.14 with two mutations in *trans*, and 0.06 with three or more mutations in *cis* over a wild-type haplotype; the frequencies of wild-type homozygotes and heterozygotes for single mutations are only 0.14 and 0.28, respectively. The frequency of individuals that are homozygous at one or more sites is approximately 0.001 in this example, so they will have a negligible effect on the haplotype dynamics compared with the heterozygous carriers of mutations, implying that the value of *k* can play only a small role when the number of selected sites is large. Positive LD under the multi-site gene model can thus arise from the *cis/trans* effect.

As mentioned above, positive LD has also been found for the sites model in recent theoretical studies that included finite population effects. Roze (2021) studied a diploid finite population model with two loci under a multiplicative version of the sites model, assuming *γ* = 2*N*_*e*_*s* >> 1 but *s* << 1. He found small positive mean *D* with *h* < 0.25 in both his approximate analytical treatment and in multi-locus simulations (see his Figure 1). Similarly, the analysis of the additive sites model for a two-locus system in a finite population by Ragsdale (2022) found positive mean values of *D* and 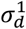 for sufficiently small values of *h*, provided that *γ* was sufficiently large. We similarly found small positive mean values of *D* and *σ*_*d*_ for the sites model when *γ* =100 and *h*=0.0 (Table 2). Ragsdale’s model of an intragenic position effect also generated positive mean LD. No such effect is expected in an infinite population (see Figure 2).

However, for very small values of the recombination rate and with *h* << ½, negative LD was found for the sites model by both Roze (2021) and Ragsdale (2022), reflecting the build-up of associative overdominance caused by the interaction between drift and selection, such that a predominance of + – and – + haplotypes is generated when linkage is close (Zhao and Charlesworth 2016; Becher et al. 2020; Gilbert et al. 2020; Abu-Awad and Waller 2023). The interactions between selection and drift when there is dominance are thus rather complex.

Our multi-locus simulations of the sites model with multiplicative rather than linear fitnesses failed to generate significantly positive LD (Table S2 in Supplementary File S3), when deleterious versus beneficial alleles were used for the sign criterion. The absence of positive LD in our simulations may be due to the stronger effects of HRI in multi-site compared with two-site models. The existence of positive LD among derived or selected mutations discussed here should not be confused with the much more frequently observed positive *D* when an excess frequency of haplotypes with pairs of minor alleles is used as the sign criterion for *D* (e.g., Figure 3). As mentioned in the section *Simulation results for multiple sites: linkage disequilibrium with no epistasis*, this effect is expected even for neutral variants (Potapova and Kondrashov 2023).

### Patterns of linkage disequilibrium in population genetic data

Previous theoretical and simulation studies have suggested that, while HRI and synergistic epistasis can create negative LD between selected variants, processes such as antagonistic epistasis, admixture and population bottlenecks could also result in positive LD (Sohail et al. 2017; Sandler et al. 2021; Garcia and Lohmueller 2021; Ragsdale 2022). Several recent studies have evaluated the sign of LD among variants in coding sequences in natural populations of *D. melanogaster*, humans, *Capsella grandiflora* and *Schizophyllum commune* (Pool et al. 2012; Sohail et al. 2017; Sandler et al. 2021; Garcia and Lohmueller 2021; Ragsdale 2022; Stolyarova et al. 2022). Garcia and Lohmueller (2021) found a larger magnitude of negative LD among doubleton nonsynonymous variants compared with synonymous variants in humans when using Lewontin’s *D’* statistic (Lewontin 1964). As described in the subsection *Effects of weighting towards low frequencies on LD statistics, D’* is biased towards negative values when low frequency variants are used, and differences in *D’* between neutral and selected sites do not necessarily reflect the effects of epistasis or HRI alone.

Sohail et al. (2017) and Sandler et al. (2021) used estimates of LD based on the excess or deficiency over the Poisson expectation of the variance in the numbers of minor alleles over a set of defined genomic regions, which is proportional to the sum of pairwise *D* values over the regions in question, whereas Stolyarova et al. (2022) used estimates of the correlation coefficients between pairs of variants, polarizing with respect to minor versus major alleles. Both of these methods are likely to give higher magnitudes of the LD statistic than when combinations of derived alleles are used for determining the sign of LD (Potapova and Kondrashov 2023). Ragsdale (2022) polarized LD using the ancestral/derived criterion; he employed the 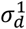 measure, whose denominator involves the cross products of the four variant frequencies at a pair of diallelic sites (see Equation 7). This will tend to give *higher* magnitudes for sites with lower frequencies, for a given correlation coefficient.

These methodological differences across studies make it hard to compare the results, but nevertheless there are some patterns in common. Notably, in most cases the magnitude of signed LD between synonymous sites was found to be significantly higher than that between nonsynonymous variants, with the exception of the analysis of the hypervariable mushroom species *S. commune* by Stolyarova et al. (2022), where the opposite pattern was observed – see their Appendix 2, Figure 1.

There seems to be a preponderance of slightly negative values of LD between loss-of-function (LOF) mutations (presumed to be the most harmful), but positive values between pairs of synonymous and nonsynonymous variants. Because these patterns applied to loosely linked variants in humans and *Drosophila*, and the extent of under-dispersion of loosely linked loss-of-function alleles was found to be much higher than explained by simulations that included the demographic history of humans, Sohail *et al*. (2017) proposed that synergistic epistasis was responsible.

A possible explanation for a predominance of negative signed LD between loosely linked LOF mutations is that these are kept at very low frequencies in the population. This means that a finite sample from the population has a high probability of not containing the – – haplotype at a pair of diallelic sites. This automatically generates *D* < 0, even if *D* is zero in the population as a whole, as would be expected for most pairs of sites that are not tightly linked. With two sites with *D* = 0 in the population and a minor allele frequency *q* at each site, a sample of size *k* has a probability of approximately [1 – exp(–*kq*)]^2^ of segregating at both sites, and a probability of approximately *kq*^2^ of containing the – – haplotype, giving a net probability of 1 – *kq*^2^/[1 – exp(–*kq*)]^2^ that a segregating sample contains no – – haplotypes. With *k* = 1000 and *q* = 0.001, this probability is equal to 0.9975 for 10000 replicate pairs. But with *q* = 0.01, it is only 0.9. Despite the fact that the expected value of *D* is zero in both cases, with *q* = 0.001 a finite set of pairs of segregating sites is likely to have a high probability of containing none or only a few – – haplotypes, resulting in a large preponderance of cases with *D* < 0. This is illustrated in section of Supplementary File S1. It is thus possible that observations of negative LD between loosely linked or independent LOF mutations, such as those shown in Figure S6 of Sohail et al. (2017) are the result of this sampling effect. It follows that it cannot safely be inferred that synergistic epistasis is the cause of this pattern unless a very large number of pairs of variants is analysed; however, our analytical and simulation results confirm that it does indeed generate negative LD for the sites model but not necessarily for the gene model (see Figures 2 - 4). As noted by Ragsdale (2022, p.10) only a few hundred pairs of within-gene LOF mutations were present in his human population dataset.

Our modeling work adds a new perspective on findings of positive LD among closely linked pairs of putatively selected variants. If fitness is governed by the gene model, then LD between variants at non-synonymous sites within the same gene should be relatively larger than under the sites model, and the sign should often be positive, provided the effects of sampling just discussed do not overwhelm the effects of selection. However, as we have discussed in the section *The role of complementation*, the gene model is unlikely to apply between different genes but is likely to be applicable within a gene. Our models thus make the following predictions about signed LD patterns in population genomics data. (*a*) LD between derived or deleterious alleles at nonsynonymous sites within genes should be positive or nearly zero (assuming that the population is panmictic and at equilibrium). (*b*) LD between derived or deleterious nonsynonymous variants in the same gene should be higher than between variants in different genes. (*c*) There should be higher LD between derived nonsynonymous vs derived synonymous variants within the same gene.

There are, however, several caveats that need to be borne in mind when using these findings to interpret data. First, the sign of LD statistics when calculated using minor/major alleles (as opposed to using derived/ancestral) is almost always positive, unless the minor alleles are very rare (Potapova and Kondrashov 2023). This applies to neutral sites (Potapova and Kondrashov 2023) and to deleterious variants that obey the sites model as well as the gene model – see Figure 3. It is thus not surprising that the studies which polarize their variants by the minor/major allele criterion (Pool *et al*. 2012; Sohail et al. 2017; Sandler et al. 2021; Stolyarova et al. 2022) often found positive LD. Second, weighting towards low allele frequencies (Good 2022) also tends to promote positive LD, as described in the subsection *Effects of weighting towards low frequencies on LD statistics*, and should thus be avoided if the sign of LD is used for inferences about selection. It follows that patterns of signed LD using this criterion do not necessarily imply effects of admixture, non-equilibrium demography, or antagonistic/complementary epistasis, which have all been proposed as possible explanations in these previous studies.

This leaves the study of humans by Ragsdale (2022), who found that LD between derived missense variants was found to be significantly positive within the same domain of a gene, as opposed to near-zero values between SNPs in different domains, even after matching the distance between SNPs found within compared between domains. This observation could be explained by a gene model operating at the level of domains within genes, perhaps because there is higher probability of complete failure of complementation within domains versus across the entire gene. Ragsdale’s estimates of 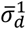 from the Yoruba population (which is closest to having a stable size) between missense variants within genes is ∼0.02 (his Figure 8). As the average length of a gene in humans is quite large (∼ 60 kb), it is difficult to compare this estimate to that obtained in our simulations. Other estimates of 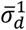 within 100 base pairs provided by Ragsdale (2002) range from 0.25-0.3 (within the same gene; his Figure S17), ∼0.5 (within the same domain; his Figure S18), and ∼0.2 (outside domains; his Figure S19). While estimates of 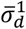 within 100 bases in our simulations of human-like populations are within this range (Table S4 in Supplementary File S3), the large SEs make it difficult to make any reliable conclusion. In general, however, both demographic processes (like admixture and bottlenecks) as well as selective processes such as positive epistasis can result in positive LD between derived nonsynonymous variants, and joint modeling of these processes in a specific population would be required to distinguish between these scenarios, taking into account important features such as recombination hotspots, which strongly influence the rates of recombination within genes in taxa such as mammals and flowering plant (e.g., Myers et al. 2010; Choi et al. 2013; Baudat et al. 2014; Sandler et al. 2021; Joseph et al. 2024).

There are multiple reasons why within-gene LD should be larger in magnitude, and more likely to be positive, than between-gene LD: (a) On average, there is a much higher frequency of recombination between pairs of sites within different genes than within the same gene (b) Fitness follows the gene model within genes but not between genes (c) There might be antagonistic epistasis within genes but not between genes. While it might be possible to differentiate between (a) and (b) by comparing the patterns of LD between species with compact genes with little spacing and/or low recombination rates between genes with patterns in species with high recombination rates between genes, it will be harder to distinguish between (b) and (c).

In summary, it will probably be quite difficult to distinguish among the different possible explanations for patterns of signed LD from population genomic data, but it is essential to use the ancestral/derived or beneficial/deleterious criterion rather than the minor/major allele criterion for the sign of LD. The problem is further complicated by the fact that, if average gene lengths and rates of recombination are sufficiently large, even patterns of LD within genes may largely reflect genome-wide patterns generated by demographic processes rather than the effects of selection. Analyses of large-scale resequencing data in the light of population genetic models of the type investigated here, using suitable choices of species with respect to their genomic properties and demographic histories, offer the best way to approach this problem.

### Evolution of the recombination rate within genes

Previous models of recombination in the presence of selection against recurrent deleterious mutations have shown that higher rates of recombination are likely to evolve if there is synergistic epistasis among mutations at different sites, and reduced rates to evolve if there is antagonistic epistasis (Feldman et al. 1980; Kondrashov 1982, 1988; Charlesworth 1990; Otto and Feldman 1997; Roze 2021). As we have shown, under the gene model, pairs of deleterious mutations result in a smaller fitness cost in *cis* than when they are in *trans*, promoting positive linkage disequilibrium. This suggests that recombination modifiers that increase the recombination rate within genes are likely to be disfavored, whereas modifiers that reduce them will be favored. This is consistent with the fact that recombination hotspots are often found in promoter regions or transcription start sites upstream of genes, and that genes are often relatively “cold” with respect to recombination rates (Myers et al. 2010; Joseph et al. 2024). Further modeling of this possible consequence of the *cis-trans* effect for the evolution of recombination rates would seem to be desirable.

## Acknowledgments

We thank Kirk Lohmueller, Denis Roze and three anonymous reviewers for their helpful comments on an earlier version of the manuscript. This research was conducted using computational resources provided by ITS Research Computing at the University of North Carolina at Chapel Hill. PJ was funded by the National Institute of General Medical Sciences of the National Institutes of Health under the award number R35GM154969.

## Appendix

### 1. Deterministic treatment of the sites and gene models: recursion equations for the haplotype frequencies

The full recursion relations for a general fitness matrix and mutation model are given below; these differ slightly in form from those usually used for models of mutation and selection, where the changes in haplotype frequencies due to selection and mutation are added to each other (e.g., Haldane [1937]). This is because the small equilibrium values of *D* resulting from the interaction between mutation and selection are very sensitive to approximations, and incorrect signs of *D* can result if approximations are made prematurely.

Assume that there are two diallelic loci, A and B, with alleles A_1_, A_2_ and B_1_, B_2_, with subscript 1 corresponding to + in Table 1 and subscript 2 to –. Let the frequencies of the four haplotypes A_1_B_1_, A_1_B_2_, A_2_B_1_ and A_2_B_2_ be *x*_1_, *x*_2_, *x*_3_ and *x*_4_, respectively. The corresponding allele frequencies are *p*_1_, *q*_1_ (A_1_ and A_2_) and *p*_2_, *q*_2_ (B_1_ and B_2_). The coefficient of linkage disequilibrium is *D* = *x*_1_*x*_4_ – *x*_2_ *x*_3_. Assume that A_2_ and B_2_ are both deleterious and are kept at such low frequencies that mutations can be treated as unidirectional, with mutation rates *u*_1_ and *u*_2_ for A_1_ to A_2_ and B_1_ to B_2_, respectively. Let *w*_*ij*_ be the fitness of individuals formed by a pair of haplotypes (*i* and *j*) with frequencies *x*_*i*_ and *x*_*j*_. Under the gene model, the fitnesses of the two double heterozygotes (*w*_14_ and *w*_23_) are unequal, in contrast to the usual assumptions of two-locus models (e.g., Karlin 1975). The marginal fitness of haplotype is *w*_*i*_ = ∑_*j*_ *x*_*j*_*w*_*ij*_ and the population mean fitness is 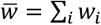.

Recombination occurs between the two loci with frequency *r* per generation. Random mating is assumed.

Assume that the order of events within a generation is selection, mutation and recombination. The post-selection frequency of haplotype *i* is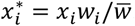. Mutation changes the haplotype frequencies to new values 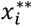 as follows:

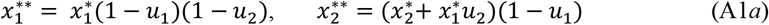

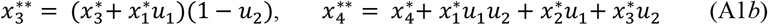

In the absence of recombination, these are the haplotype frequencies in the new generation. If recombination occurs post-mutation, the haplotype frequencies in the new generation are given by the following expressions, where the triple asterisks indicate haplotype frequencies after mutation without any selection within the generation in question (*i*.*e*, the frequencies given by Equations (A1) without the asterisks on the right-hand side); the effects of selection on the double heterozygotes are captured by the terms in *w*_14_ and *w*_23_:

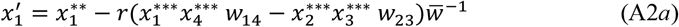

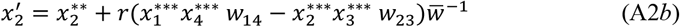

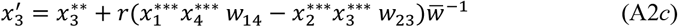

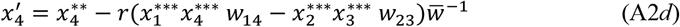

These recursion equations can be used to obtain the equilibrium values for the system, as well as the approach to equilibrium. Given the symmetry of the fitness matrix in Table 1, for the system studied here we can write *x*_2_ = *x*_3_ = *x* = *q*(1 – *q*) – *D; x*_4_ = *y* = *q*^2^ + *D*; *x*_1_ = *z* = 1 – 2*x* – *y*; *q* = *x* + *y*. It follows that only the recursions for *x* and *y* need to be considered. Let the mutation rate per locus from + to – be *u*. If *u << hs*, the frequency *q* of the – allele at each locus is << 1 and reverse mutations can be ignored. The coefficient of linkage disequilibrium is denoted by *D* = *yz* – *x*^2^. Together with the fitness values for each genotype, these variables can be used to determine the recursion relations for the frequencies of the four haplotypes, *x*_1_ (+ +), *x*_2_ (– +), *x*_3_ (+ –) and *x*_4_ (– –).

Denote the marginal fitnesses of the – + and + – haplotypes by *w*_*x*_ and that of the – – haplotype by *w*_*y*_; the marginal fitness of the + + haplotype is *w*_*z*_. From Table 1, we obtain the following expressions for the marginal fitnesses under the sites model:

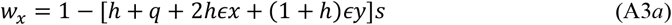

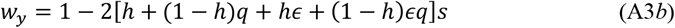

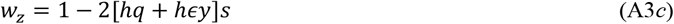

The equivalent expressions for the gene model are as follows:

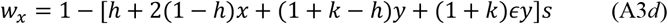

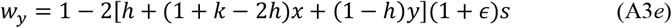

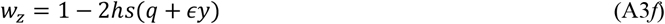

For both models, the mean fitness of the population is given by:

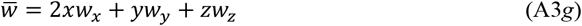

If these expressions are substituted into Equations (A1), using specified values of the mutation rate, recombination rates and selection parameters, a numerical solution to the equilibrium version of the recursions could be found by an iterative method. It is, however, simpler to run the recursions to close to equilibrium, thereby obtaining estimates of the equilibrium values of *x* and *y* and all derived quantities of interest. The initial starting point of the recursions was arbitrarily chosen to have *D* = 0, with *q* given by the solution of the linear equation that approximates the cubic equation describing mutation-selection equilibrium at a single locus when *q* is sufficiently small but non-negligible (Bürger 2000, p.100):

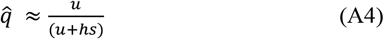

### 2. Properties of equilibria with respect to LD under the additive and multiplicative sites models

As shown in part 1 of Supplementary File S1, when *r* = 0, a formula for the equilibrium value of 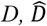, can be obtained for a general symmetric fitness model (Equation S1.3). Neglecting the small terms in the mutation rate in this expression, we have:

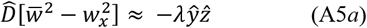

where

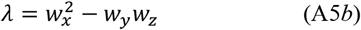

The analysis in section 2 of Supplementary File S1 shows that 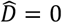 under the multiplicative sites model with no epistasis (Equation S1.6b).

Equations (S1.7) and (S1.8) in section 3 of Supplementary File S1 show that, for the additive sites model with no epistasis, we have:

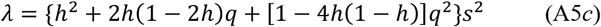

We can approximate *q* for an equilibrium population by equating it to the equilibrium solution 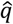 given by Equation (A4). We can also approximate the haplotype frequencies by neglecting the small terms involving 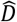, so that 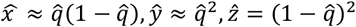. Ignoring terms in 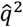, Equation (S10b) then gives the following approximation:

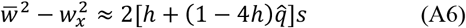

Equations (A5*c*) and (A.6) yield the following approximation for 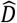:

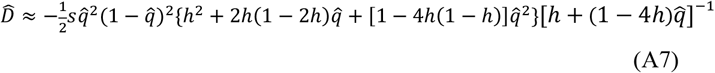

This expression can be extended to include the effect of recombination when *r, D* and *s* are sufficiently small that their second-order terms can be neglected. In this case, Equations (A2) imply that the change in *D* caused by recombination is approximately – *rD*. If this term is added to the right-hand side of the recursion equation for *D* (Equation S1.3), and 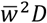 is approximated by *D*, Equation (A5*a*) becomes:

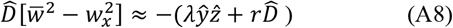

This yields the following approximate expression for 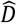:

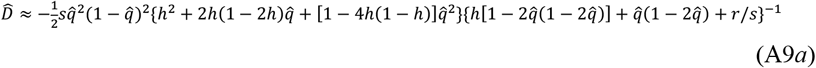

This equation can be converted into an expression for the equilibrium value of the correlation coefficient, 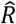, by dividing by 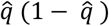:

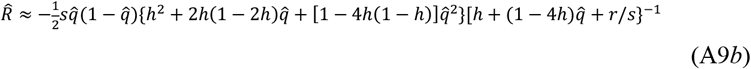

If second-order terms in 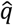 are neglected, this expression simplifies to:

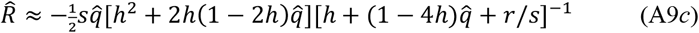

The magnitudes of 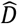 and 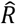 under the additive sites model are, however, very small in the absence of epistasis, of the order of the mutation rate. This can be seen as follows. Provided *h* is bounded away from 0, and *u* << *hs*, we have 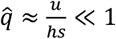 (Haldane 1937). The leading term in braces in the numerator of Equation (A9*b*) is *h*^2^, and the leading term in braces in the denominator is *h*, so that the magnitude of the ratio of numerator to denominator in Equation (A9*b*) is approximately 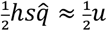. If 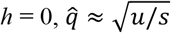, so that the term in braces in the numerator is approximately *u*/*s* and the term in braces in the denominator is approximately 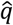. This cancels the 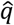 term in the numerator, so that (assuming 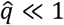), the magnitude of the ratio of numerator to denominator is approximately 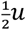, as before. It follows that, provided that 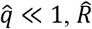 should be approximately equal to the negative of one-half of the mutation rate, and independent of the dominance coefficient and strength of selection. Provided that the epistasis coefficient *ϵ* is of the same order as *s* or greater, much larger negative values of 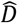 and 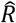 are expected with synergistic epistasis (*ϵ* > 0), and positive values are expected with antagonistic epistasis (*ϵ* < 0) – see Equation (A9*c*).

Equations (A9) must be modified if epistasis is present, which introduces extra terms of the order of *ϵs*. Use of Equations (A3) leads to the following expression for the additive sites model when *ϵ* > *s*:

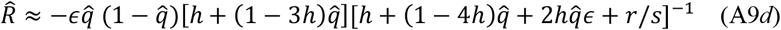

### 3. Properties of equilibria with respect to LD under the additive and multiplicative gene models

The same type of approach can be applied to the additive and multiplicative gene models, as described in section 4 of Supplementary File S1. Equation (S1.16c) yields the following approximation for 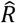:

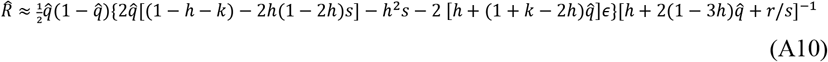

Similarly, for the multiplicative gene model without synergistic or antagonistic epistasis, Equation (S1.16d) yields:

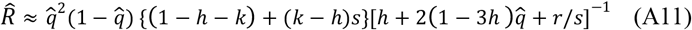

For *h* ≤ ½, and 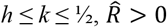.

### 4. Extension to a large number of selected sites

It is possible to derive an approximation for the LD measures considered above in the case of a large number of sites subject to the same selection regime. It is assumed that Equation (A8) applies to a given pair of sites, provided that expressions for *w*_*x*_, *w*_y_ for such a pair of sites, as well as for 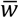, can be obtained. The equilibrium allele frequency at each site, 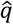, is treated initially as a given. For the purpose of comparisons with the simulation results, we consider only the additive gene model with *k* = ½. The effects of LD on the distribution of the number of deleterious mutations per haplotype are ignored, so that the probability that a haplotype carries *i* mutations (*P*_*i*_) is assumed to follow a Poisson distribution with mean 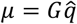, where 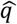 is the equilibrium frequency of mutant alleles at a given site (*G* >> 1):

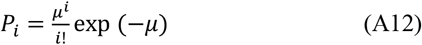

Consider first the net marginal fitness of a haplotype that lacks mutations at a pair of randomly chosen sites, *w*_z_. Such a haplotype may carry 0, 1, 2, … *G* – 2 mutations located at other sites. Ignoring LD, the probability of *i* mutations at other sites is *P*_*i*_; the marginal fitness of such a haplotype is denoted by *w*_*zi*_. Taking into account the other haplotypes that are encountered to form diploid individuals, and using the fitness function for the gene model with *k* = ½, we have:

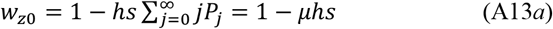

For *i* ≥ 1, we have:

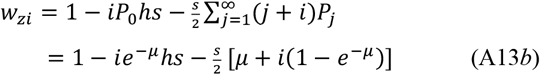

This yields:

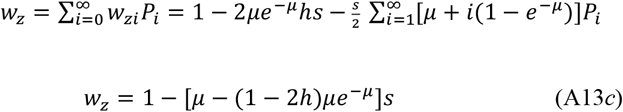

Next, consider the marginal fitness of a haplotype with a mutation at one of the two chosen sites. As before, the probability that such a haplotype carries *i* mutations at other sites is *P*_*i*_ and the marginal fitness of such a haplotype is:

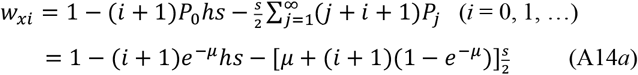

so that:

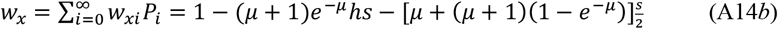

The same procedure can be carried out for the marginal fitness of a haplotype with mutations at both chosen sites, *w*_*y*_. The fitness of such a haplotype that carries *i* mutations at other sites is:

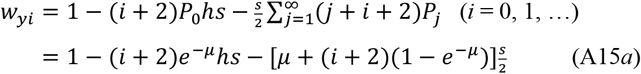

and:

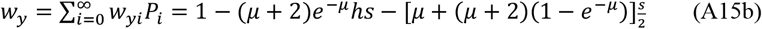

If second-order terms in *s* are neglected, we have:

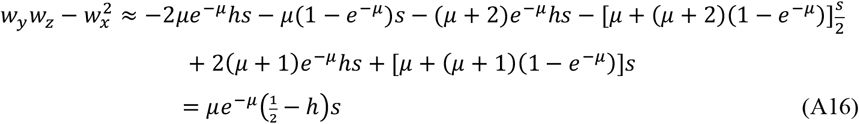

Furthermore, if 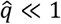 and *s* << 1, we have:

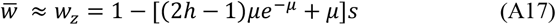

Using this expression together with Equation (A14*b*), we obtain:

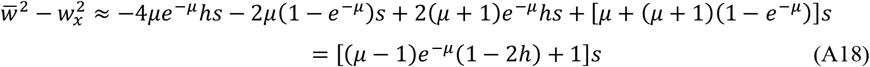

Substituting from Equations (A16) and (A18) into Equation (A8) yields the expressions:

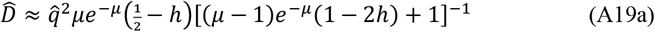

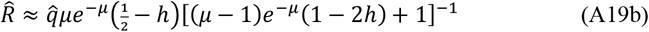

Equations (A13*c*) and (A14*b*) can be used to obtain an equation for the value of *μ* and hence 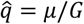. Provided that *q* << 1, the change in *q* due to selection at a given site is given by:

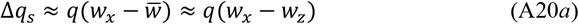

Substituting from Equations (A13*c*) and (A14*b*) and simplifying, the following expression is obtained:

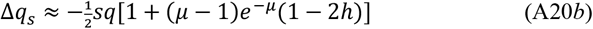

At mutation-selection equilibrium, this must balance the change in *q* due to mutation, which is approximately equal to the mutation rate *u*. The equilibrium value of *μ* is then determined by the following transcendental equation, which can be solved numerically by Newton-Raphson iteration:

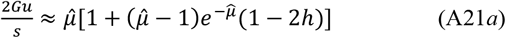

With semi-dominance (*h* = ½), this equation is equivalent to the standard equation

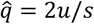 (Haldane 1927), as expected from the equivalence of the sites and gene models in this case. For smaller values of 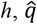 is reduced below this value, as seen in the two-site model described above. Importantly, for 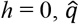 does not obey the standard single locus result for a completely recessive mutation, 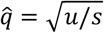 (Haldane 1927), but instead takes much larger values.

But for sufficiently small values of *G*, 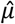 is << 1, and Equation (A21*a*) can be approximated by:

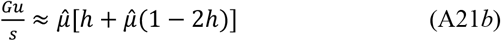

When *h* = 0, this yields the standard equation for complete recessivity. When 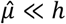, this reduces to the standard Haldane equation 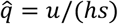.

### 5. The effect of using Lewontin’s *D*’statistic on the sign of LD

Let the allele frequencies at the two sites in question, A and B, be *p*_*A*_, *q*_*A*_, *p*_*B*_ and *q*_*B*_, where *p* and *q* refer to ancestral and derived variants, respectively. If *D* is defined as positive for associations between derived variants, we have:

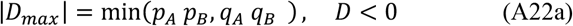

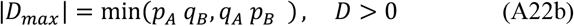

First, consider the situation if unweighted haplotype frequencies from the whole population are used for the LD statistics. As described in the section *Methods: two-locus simulations*, the LD statistics were calculated after a second mutation at site B, was introduced at a site segregating for a mutation at site A, either in *cis* with the ancestral A variant (case 1) or in *cis* with the derived variant (case 2). For case 1 in the neutral model described above, *D* < 0 for all generations, and *D*^′^ = − *q*_*A*_*q*_*B*_/*q*_*A*_ *q*_*B*_ = – 1 if *q*_*A*_ *q*_*B*_ < *p*_*A*_ *p*_*B*_ or *D*^′^ = − *q*_*A*_*q*_*B*_/*p*_*A*_ *p*_*B*_ > – 1 if *q*_*A*_ *q*_*B*_ > *p*_*A*_ *p*_*B*_. Given that *q*_*A*_ and *q*_*B*_ are the frequencies of derived variants, whose probability distribution is skewed towards low frequencies, the large majority of cases will involve the first situation, and the expectation of *D*’ for case 1 will be – 1 or close to – 1.

For case 2, *D*^′^ = *p*_*A*_*q*_*B*_/*p*_*A*_*q*_*B*_ = 1 if *p*_*A*_ *q*_*B*_ < *q*_*A*_ *p*_*B*_ and *D*^′^ = *p*_*A*_*q*_*B*_/ *q*_*A*_*p*_*B*_ > 1 if *p*_*A*_ *q*_*B*_ > *q*_*A*_ *p*_*B*_. Again, the majority of cases will satisfy the first condition, since *q*_*B*_ is likely to be much smaller than *q*_*A*_, due to B_2_ arriving in the population as the second mutation. The expectation of *D*’ for case 2 will thus be 1 or close to 1. Case 1 has a higher probability of occurring than case 2 (their probabilities are proportional to the integrals between 1/*N*_*H*_ and 1 – 1/ *N*_*H*_ of *pq*^−1^ and 1, respectively), so that the overall expectation of *D’* is negative. If purifying selection is acting, the probability of case 1 is increased over case 2, due to the increased skew of the probability distribution of allele frequencies towards low values, so the magnitude of *D*’ will increase with the strength of selection.

The properties of *D*’ for doubleton variants are described in section 8 of Supplementary File S1.

