## Supplementary File S1 for "A gene-based model of fitness and its implications for genetic variation: Linkage disequilibrium"

### Supplementary Appendix

#### 1. A recursion relation for *D*

A recursion relation for *D* can be derived from Equations (A1), by a similar method to that used by Felsenstein (1965) for the recursion of the quantity $Z = \ln(x_1x_4/x_2x_3)$, with the addition of a term involving the mutation rates. To obtain insights into the properties of the system with low frequencies of recombination, $r = 0$ is assumed. Without much loss of generality, second-order terms in the mutation rates can be neglected, so the following results are accurate to terms of order $u_i^2$.

Equations (A1) imply that the new value of *D* (*D′*) is given by:

$$\bar{w}^2D' = x_1w_1(1-u_1-u_2)[x_4w_4 + x_2w_2u_1 + x_3w_3u_2] - [x_2w_2(1-u_1) + x_1w_1u_2][x_3w_3(1-u_2) + x_1w_1u_1] \quad \text{(S1.1a)}$$

Expanding the expressions in brackets and cancelling terms, we obtain the following expression:

$$\bar{w}^2D' = (x_1x_4w_1w_4 - x_2x_3w_2w_3)(1-u_1-u_2) \quad \text{(S1.1b)}$$

Using the notation for the marginal fitnesses under a symmetric fitness model of the type used here ($x = x_2 = x_3$, $y = x_4$, $z = 1 - 2x - y$), the recursion for *D* can be written as:

$$\bar{w}^2D' = (Dw_x^2 - \lambda yz)(1-2u) \quad \text{(S1.2a)}$$

where

$$\lambda = w_x^2 - w_yw_z \quad \text{(S1.2b)}$$

The equilibrium value of *D*, $\hat{D}$, must therefore satisfy:

$$\hat{D}[\bar{w}^2 - w_x^2(1-2u)] = -\lambda\hat{y}\hat{z}(1-2u) \quad \text{(S1.3)}$$

**2. The multiplicative sites model**

Since the equivalent of the sites model with multiplicative fitnesses is widely use in models of multi-locus systems with mutation and selection (e.g, Roze 2021), we will first examine the properties of the multiplicative version of the sites model with no epistasis, as displayed in the following fitness matrix.

**Fitness matrix for the sites model with multiplicative fitnesses**

| | ++ | + - | - + | - - |
|---|---|---|---|---|
| + + | 1 | $1 - hs$ | $1 - hs$ | $(1 - hs)^2$ |
| + - | $1 - hs$ | $1 - s$ | $(1 - hs)^2$ | $(1 - s)(1 - hs)$ |
| - + | $1 - hs$ | $(1 - hs)^2$ | $1 - s$ | $(1 - s)(1 - hs)$ |
| - - | $(1 - hs)^2$ | $(1 - s)(1 - hs)$ | $(1 - s)(1 - hs)$ | $(1 - s)^2$ |

Equations (A3*a*)-A(3*c*) show that the marginal fitnesses for the additive version of the sites model are independent of *D*. Each marginal fitness under multiplicativity for a given deleterious allele frequency *q* is thus equal to its value with $D = 0$, plus the terms in *D* in the expressions for the marginal fitnesses that contribute to a deviation from additivity under the multiplicative model, e.g., $D(hs)^2$ is contributed by – – haplotypes to the marginal fitness of the + + haplotype. Using subscript 0 for the marginal fitness values when $D = 0$, we obtain:

$$w_x = w_{x0} + Dh(1-h)s^2, \; w_y = w_{y0} + D(1-h)^2 s^2, \; w_z = w_{z0} + Dh^2 s^2 \qquad \text{(S1.4a)}$$

and

$$w_{x0} = (1 - qhs)\{1 - [h + q(1-h)s]\} \qquad \text{(S1.4b)}$$

$$w_{y0} = \{1 - [h + q(1-h)s]\}^2 \qquad \text{(S1.4c)}$$

$$w_{z0} = [1 - qhs]^2 \qquad \text{(S1.4d)}$$

These expressions imply that:

$$w_{x0} w_{y0} - w_{x0}^2 = 0 \qquad \text{(S1.5a)}$$

and so:

$$w_y w_z - w_x^2 = w_{x0} w_{y0} - w_{x0}^2 + D\,O(s^2) + D^2 O(s^4)$$

$$w_y w_z - w_x^2 = D\,O(s^2) + D^2 O(s^4) \qquad \text{(S1.6a)}$$

After some algebra, it can be seen that the $O(s^4)$ term is zero, and that:

$$\bar{w}^2 D' = D[(1 - 2h + h^2 s]^2 (1 - 2u) s^2 \quad \text{(S1.6b)}$$

The right hand side lies between 0 and 1, so that the only possible equilibria must have $D = 0$.

**3. The additive sites model**

The properties of the additive sites model in the absence of recombination and epistasis can be examined by the following procedure. From Equations (A3$a$)-(A3$b$) of the Appendix to the main text, we have:

$$w_x^2 = 1 - 2(h + q)s + (h + q)^2 s^2 \quad \text{(S1.7a)}$$

$$w_y w_z = 1 - 2(h + q)s \ + 4[h + (1 - h)q]hs^2 q \quad \text{(S1.7b)}$$

After some simplification, these expressions yield the following result:

$$\lambda = w_x^2 - w_y w_z = \{h^2 + 2h(1 - 2h)q + [1 - 4h(1 - h)]q^2\}s^2 \quad \text{(S1.8)}$$

This expression shows that a sufficient condition for $\lambda > 0$ is $h \leq \frac{1}{2}$, so that $\lambda > 0$ for all cases of interest. From Equation (S1.8), $\lambda = O(s^2)$, as would be expected from the fact that the additive and multiplicative models differ by $O(s^2)$ terms.

Since $2u < 1$ by hypothesis, a sufficient condition for $\hat{D} < 0$ is $\bar{w} \geq w_x$, which is intuitively likely, since $w_x$ refers to haplotypes carrying a single mutant allele, which are less frequent than wild-type haplotypes but much more frequent than double mutant haplotypes.

It can be shown as follows that $\bar{w} \geq w_x$ holds under light conditions. We have:

$$\bar{w} - w_x = (1 - 2x - y)w_z + (2x - 1)w_x + y w_y$$

or

$$\bar{w} - w_x = (w_z - w_x)(1 - 2x) + y(w_y - w_z) \quad \text{(S1.9a)}$$

Furthermore, we have:

$$w_z - w_x = [h + q(1 - 2h)]s \quad \text{(S1.9b)}$$

$$w_y - w_z = -2[h + q(1 - 2h)]s \quad \text{(S1.9c)}$$

These relations follow from the fact that $[h + q(1 - 2h)]s$ is the additive effect on fitness of each locus.

We thus obtain:

$$\bar{w} - w_x = [h + q(1 - 2h)][1 - 2q(1 + q)]s \quad \text{(S1.10a)}$$

If $q$ < ½ or $h$ < ½, both very light conditions, $\bar{w} - w_x < 0$ and $\hat{D} < 0$, as expected.

Using these relations and Equations (A3), after neglecting $O(q^2)$ terms we have:

$$\bar{w}^2 - w_x^2 \approx 2[h + (1 - 4h)q]s + O(s^2) \quad \text{(S1.10b)}$$

**4. The additive and multiplicative gene models**

A similar approach can be used for the additive gene model. First, assume a lack of narrow sense epistasis. Using Equations (A3*d*)-(A3*f*) of the Appendix to the main text, we have:

$$w_x^2 = 1 - 2[h + 2(1 - h) + (1 + k - h)y]s + O(s^2) \quad \text{(S1.11a)}$$

$$w_y w_z \approx 1 - 2\{qh + h + (1 + k - 2h)x + (1 - h)y\}s + O(s^2) \quad \text{(S1.11b)}$$

After some simplification, we obtain the expression:

$$w_x^2 - w_y w_z \approx -2sq(1 - h - k] + O(s^2) \quad \text{(S1.12)}$$

Furthermore, applying the approach used to obtain Equations (S1.9) and (S1.10), approximating $x$ by $q$ (*i.e*, neglecting $D$) and neglecting terms in $y$, we have:

$$\bar{w} - w_x \approx [h + 2(1 - 3h)q]s \quad \text{(S1.13)}$$

If $h \leq$ ½ and $q$ < ½, $w_x < \bar{w}$. By the argument that yielded Equation (A7), $\hat{D}$ is given by the expression:

$$\hat{D}[(\bar{w}^2 - w_x^2(1 - 2u)] = 2s\hat{q}\hat{y}\hat{z}(1 - h - k)(1 - 2u) + O(s^2) \quad \text{(S1.14)}$$

The equilibrium value of $D$ is thus positive when $h + k < 1$ and $\hat{q} < \frac{1}{2}$, provided that the term in $O(s^2)$ is sufficiently small.

When $h = k = ½$, however, the additive sites and gene models are equivalent in the absence of narrow sense epistasis and $\widehat{D} < 0$, indicating that the $O(s^2)$ term in the expression for the gene model must then be sufficiently large to affect the sign of $D$. The $O(s^2)$ terms in $w_x^2$ and $w_y w_z$ are as follows:

$$w_x^2: \quad [h + 2(1-h)x + (1+k-h)y]^2 s^2 \qquad \text{(S1.15}a\text{)}$$

$$w_y w_z: \; 4[h + (1+k-2h)x + (1-h)y]qhs^2 \qquad \text{(S1.15}b\text{)}$$

The leading terms in these quantities when $q << 1$ and $D << q$ are :

$$w_x^2: \quad [[h^2 + 4h(1-h)q]s^2, \;\; w_y w_z: \; 4qh^2 s^2$$

Their contribution to $w_y w_z - w_x^2$ is thus approximately $(1-2h)q - 4h(1-2h)qs^2 - h^2 s^2$. The full approximate expression for $\widehat{D}$ with $\hat{q} << 1$ is thus:

$$\widehat{D}[\overline{w}^2 - w_x^2] \approx s\hat{q}^2(1-\hat{q})^2[2\hat{q}[(1-h-k) - 2h(1-2h)s] - h^2 s] \qquad \text{(S1.16a)}$$

Neglecting terms of order $q^2$, $Dq$ and $s^2$, we also have:

$$\overline{w}^2 - w_x^2 \approx 2(w_z - w_x) \approx 2[h + 2(1-3h)q]s \qquad \text{(S1.16}b\text{)}$$

If the contributions from terms in $\epsilon s$ are included, the final approximation for $\widehat{D}$ for the additive gene model is as follows:

$$\widehat{D} \approx \frac{1}{2}s\hat{q}^2(1-\hat{q})^2\{2\hat{q}[(1-h-k) - 2h(1-2h)s] - h^2 s - [h + (1+k-2h)\hat{q}]\epsilon\}[h + 2(1-3h)\hat{q} + r/s]^{-1} \qquad \text{(S1.16}c\text{)}$$

The multiplicative version of the gene model (without narrow sense epistasis) has the fitness matrix shown below. It does not reduce to the multiplicative sites model when $h = k = ½$, since the fitnesses of the + –/+ - and + –/– + genotypes are both $1 - s$ in this case, compared with $(1 - hs)^2$ and $1 - s$, respectively, for the sites model.

| | + + | + - | - + | - - |
|---|---|---|---|---|
| + + | 1 | $1 - hs$ | $1 - hs$ | $1 - 2hs + (hs)^2$ |
| + - | $1 - hs$ | $1 - s$ | $1 - s$ | $1 - (1+k)s + ks^2$ |
| - + | $1 - hs$ | $1 - s$ | $1 - s$ | $1 - (1+k)s + ks^2$ |
| - - | $1 - 2hs + (hs)^2$ | $1 - (1+k)s + ks^2$ | $1 - (1+k)s + ks^2$ | $1 - 2s + s^2$ |

If $x$ and $y$ (and hence $q$) are << 1, the leading contribution from terms in $s^2$ in the expressions for $w_x$, $w_y$ and $w_z$ comes from the $(hs)^2$ contribution to the fitness of – –/+ +, which is added to $w_y$. There is an additional contribution to $w_y$ of approximately $2q(k - h^2)s^2$. After including these terms in the numerator of the expression for $\widehat{D}$, the first of these terms contributes ½ $h^2s$ after the factor of $2s$ in the denominator is cancelled. It follows that the $-h^2s$ term in the numerator of Equation (S1.16a) disappears. After adding the additional terms in $s$ to the $2qs$ term in numerator of the additive model, which sum to $-4qh(1 - 2h)s$ and simplifying, the following expression for the gene model with multiplicative fitnesses and no epistasis is obtained:

$$\widehat{D} \approx \hat{q}^3(1-\hat{q})^2\{(1-h-k)+(k-h)s\}[h+2(1-3h)\hat{q}+r/s]^{-1} \qquad \text{(S1.16d)}$$

Figure S1.1 shows the plot of the corresponding formula for the correlation coefficient $R$ (Equation A11) against the dominance coefficient, for the case of $u = 0.0001$, $k = 0.5$ and $s = 0.01$ (as in Figure 2 of the main text for the additive model), compared with the exact results from the recursion equations. The agreement is extremely close, and the numerical values differ only slightly from those for the additive gene model.

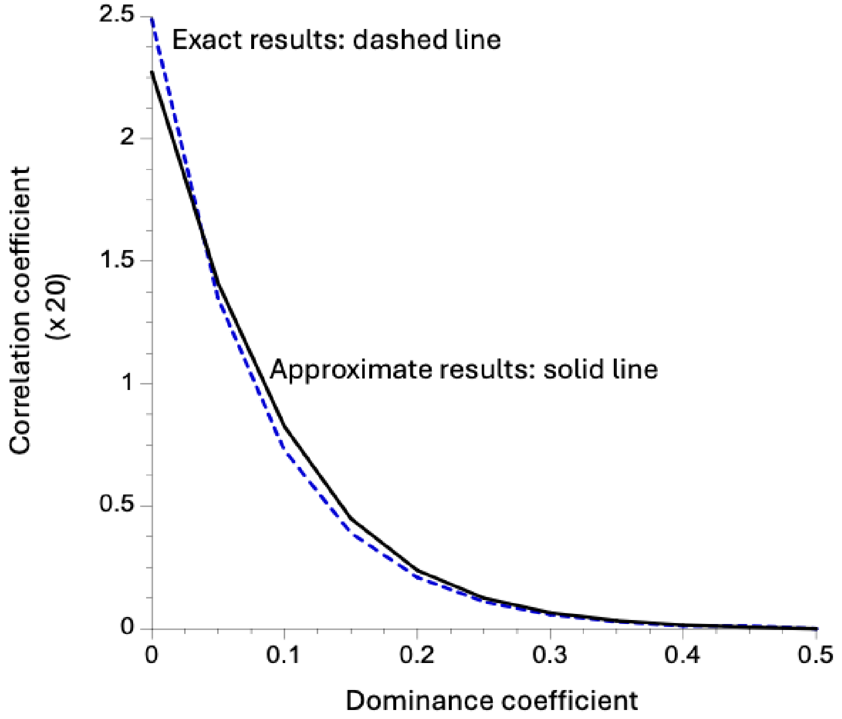


**5. The sources of LD under the two-site gene model**

There are two potential sources of LD in the two-site deterministic version of the gene model. The first is the higher fitness of *cis* versus *trans* double heterozygotes, which favors positive LD between the deleterious alleles. The second is the effect of the parameter $k$ on the fitness of – – /+ – and – – /– + genotypes; if $k > h$, this reduces the marginal fitness of – – ($w_y$) haplotypes compared with the sites model, due to the fitnesses of these genotypes being 1 – (1 + $k$)$s$ instead of 1 – (1 + $h$)$s$. This causes a reduction in the frequency of – – haplotypes, making negative LD more likely (the opposite would happen with $k < h$).

The two effects can be disentangled by considering a model in which there is no *cis*-*trans* difference, but the fitnesses of the – – /+ – and – – /– + genotypes are both 1 – (1 + $k$)$s$. If $q << 1$, the arguments used above lead to the result that, ignoring terms in $s^2$ and $q^2$, $w_x$ and $w_z$ are the same as for the sites model, whereas $w_y$ is the same as for the gene model, and is thus approximately equal to $1 - 2[h + (1 + k - 2h)q]s$. This leads to the result that the quantity that determines the sign of LD is:

$$w_y w_z - w_x^2 \approx 2q(h - k)s \qquad \text{(S1.17a)}$$

whereas the corresponding expression for the gene model is:

$$w_y w_z - w_x^2 \approx 2q(1 - h - k)s \qquad \text{(S1.17b)}$$

The difference between Equations (S1.17b) and (S1.17.a) is $2q(1 - 2h)s$; this reflects the product of $2q$ and the difference in fitness between the *cis* and *trans* double heterozygotes. With $k = ½$ and $h \leqslant ½$, as assumed in the simulations, positive LD results from the *cis-trans* difference. It is larger, the smaller $k$. Equation (S1.17a) shows that positive LD can also occur when $k < h$ in the absence of a *cis-trans* difference, but this is implausible biologically, since it implies that homozygosity for – in – – /+ – or – – /– + has a smaller effect on fitness than the heterozygous +/– site.

**6. Effects of using low allele frequencies on measures of linkage disequilibrium**

Here, we describe how we can examine the effects of using measures of LD on low frequency variants, using a two-locus haploid model that is similar to the one used by Good (2022) in order to avoid complexities due to dominance. In the absence of epistasis this should be equivalent to the diploid case with $h = ½$, which was used in many of the multi-locus simulations of Garcia and Lohmueller (2021). We investigated both the consequences of analyzing finite samples in which only doubletons with respect to derived variants were used (Garcia and Lohmueller 2021), and the procedure of weighting population haplotype frequencies by an exponential function of the allele frequencies at the pairs of sites involved (Good 2022). Recombination was assumed to be absent.

To save computer time, the simulation method of Good (2022) was used, in which a deleterious mutation $A_2$ at the first site under consideration (A) segregates at a frequency $q$ in the absence of a mutation at the second site, with wild-type allele ($A_1$) at frequency $p = 1 – q$. The probability density function for $q$ is assumed to be equivalent to that for a locus subject to irreversible mutation from wild-type to mutant, with a scaled selection coefficient of $\gamma = 2Ns$, where $N$ is the size of the haploid population and $s$ is the selection coefficient against the deleterious mutation. From standard diffusion equation results (Fisher 1930), the p.d.f. of $q$ is proportional to:

$$\phi(q) = \frac{\exp(\gamma p) - 1}{pq[\exp(\gamma) - 1]} \qquad \text{(S1.18a)}$$

More accurately, $\phi(q)$ is proportional to the density function for the sojourn time at frequency $q$ between fixation and loss of $A_2$, on a background of $B_1$ (Ewens 2004, p.167).

In the neutral limit, when $\gamma$ tends to 0, this expression reduces to:

$$\phi(q) = q^{-1} \tag{S1.18b}$$

For $q_i = i/N$, $\phi(q_i)$ can be regarded as being proportional to the probability of $i$ copies of $A_2$ being present in the population. The cumulative probability distribution of $q_i$ is thus given by:

$$\Phi(q_i) = \frac{\sum_{j=1}^{i} \phi(q_j)}{\sum_{j=1}^{N-1} \phi(q_j)} \tag{S1.19}$$

In order to initiate a replicate simulation, a uniform random number $z$ was generated and compared with the successive terms $\Phi(q_1), \Phi(q_2), \ldots \Phi(q_i)$; at the first $i$ for which $z \leq \Phi(q_i)$, the initial frequency of $A_2$ was set to $q_i$. Given $q_i$, there are probabilities $p_i$ and $q_i$ of the mutation at site B arising in a wild-type or mutant background, respectively. Comparison of a second random number with $q_i$ decided which of these possibilities was realized. The first type of event (case 1) means that the following three haplotypes were present in the population: $A_1B_1$, $A_1B_2$ and $A_2B_1$. The second type of event (case 2) involves haplotypes $A_1B_1$, $A_2B_1$ and $A_2B_2$. The fates of these haplotypes over a single generation were followed by determining their frequencies after selection, followed by trinomial sampling of $N$ post-selection haplotypes to form the next generation; this procedure was repeated until one of the two sites lost variability.

The fitness model for determining the post-selection frequencies of the haplotypes in a given generation assumed that the relative fitnesses of $A_1B_1$, $A_1B_2$, $A_2B_1$ and $A_2B_2$ are 1, $1+s$, $1+s$ and $1 + 2s(1 + \epsilon)$, respectively, where $\epsilon$ is a measure of epistasis of the same form as that used in the two-site diploid model. Simulations were run with or without weighting by allele frequencies using the method of Good (2022). In the absence of weights, the sums over all generations and replicates of $D$, $D^2$ and the allele frequency cross product $P = p_A q_A p_B q_B$ (where subscripts $A$ and $B$ denote sites A and B) were obtained. The means of $D$, $D^2$ and $P$, denoted by $\overline{D}$ , $\overline{D^2}$and $\overline{P}$, were found by dividing the respective sums of these statistics by the sum of the durations of the replicate simulations. This enabled the standardized LD statistics $\bar{\sigma}_d = \frac{\overline{D}}{\sqrt{\overline{P}}}$, $\sigma_d^1 = \frac{\overline{D}}{\overline{P}}$ (Good 2022, Ragsdale 2022) and $\sigma_d^2 = \frac{\overline{D^2}}{\overline{P}}$ to be estimated.

In addition, the effect of calculating LD statistics from variants present only at a pre-set frequency in a sample of $k$ haplotypes (Garcia and Lohmueller 2021) was determined by trinomial sampling of $k$ haplotypes each generation, recording statistics only for samples where the numbers of alleles at each site in the sample matched the pre-set allele frequency (usually 2 out of $k$). The sample LD statistics were then determined in the same way as for the whole population.

It is important to verify that focussing on doubletons biases samples towards low frequency variants. If a sample of size $k$ is used, and $D$ is calculated from variants present at frequency $j$, the mean frequency of $A_2$ is $j/k$. This can be compared with the mean frequency of $A_2$ at segregating sites without conditioning on a specific frequency, which is equal to:

$$\sum_{i=1}^{k-1} \frac{i}{ika_k} = \frac{k-1}{ka_k} \qquad \text{(S1.20)}$$

where $a_k$ is Watterson's correction factor. If $j = 2$ (doubletons), the mean conditional frequency of $A_2$ is less than the overall mean frequency at segregating sites when $2 < k - 1/a_k$. This condition is satisfied for $k \geq 6$.

When applying Good's weighting method, the step of sampling $k$ haplotypes each generation was omitted. In this case, the population frequencies of haplotypes carrying $A_2$ were weighted by $\exp(-fq_A)$ and the frequencies of haplotypes carrying $B_2$ by $\exp(-fq_B)$, where the factor $f$ is 0 if no weighting is applied, and $f > 1$ if low frequency mutant alleles receive preferential weights. The LD statistics were then calculated in the same way as for the unweighted case.

**7. Causes of positive LD with weightings towards low frequencies**

The reason that weightings towards low allele frequencies cause mean $D$ values to be positive can be understood heuristically as follows. Consider the case of complete neutrality in the absence of recombination, with the probability distribution of the frequency $q$ of derived mutations at a given site being proportional to $1/q$ (Fisher 1930). A mutation introduced at a second site will be present with an initial frequency of $1/N_H$, where $N_H$ is the number of haploid genomes in the breeding population ($N_H = 2N$ for a diploid population and $N$ for a haploid population.) There is a probability $p = 1 - q$ that the second mutation will be in *cis* with the wild-type allele at the first site (case 1), in which case $D$ takes the value $- q/N_H$. There is a probability $q$ that it will be in *cis* with the mutant allele (case 2), giving $D = p/N_H$. The respective net expected values of $D$ are thus proportional to $-pq/N_H$ and $pq/N_H$, respectively, giving an overall expectation of 0 (this is true for any probability distribution of $q$). Under neutrality and in the absence of recombination, the expectation of $D$ is known to change by a factor of $1 - 1/(2N_e)$ per generation (Hill and Robertson 1968), so that $\overline{D}$ will remain at 0, as would be expected intuitively.

In an arbitrary generation following the initial one, $\overline{D}$ for case 1 is easily seen to be equal to – $q_A q_B$ (the only haplotypes present are $A_1B_1$, $A_2B_1$ and $A_1B_2$). The magnitude of $D$ in this case increases with $q_A$. In case 2, $\overline{D} = p_A q_B$, since the only haplotypes are $A_1B_1$, $A_2, B_1$ and $A_2B_2$), and is a decreasing function of $q_A$. Both cases have the same magnitude of dependence on $q_B$. It follows that, since a lack of weighting gives an expected $D$ of zero, weightings towards lower mutation frequencies, as used by Good (2022) result in a positive expected $D$.

**8. The effect of using Lewontin's *D*′ statistic**

Using the notation employed in the previous two sections, when defining $D$ as positive for associations between derived variants we have:

$$|D_{max}| = \min(p_A\, p_B, q_A\, q_B), \quad D < 0 \qquad \text{(S1.21a)}$$

$$|D_{max}| = \min(p_A\, q_B, q_A\, p_B), \quad D > 0 \qquad \text{(S1.21b)}$$

Section 5 of the Appendix shows that unweighted estimates of mean $D'$ that take into account the sign of $D$ are biased towards negative values. It seems likely that this will also apply to estimates based on doubleton frequencies, although the expectation of $D$ itself in this case is biased towards positivity unless there is strong synergistic epistasis (see Figure 5). This expectation was tested by using the simulations described above to estimate mean $D'$ from doubletons. For the cases of neutrality, $\gamma = 10$ and $\gamma = 50$ (with no epistasis), the mean $D'$ values for cases 1 and 2 were – 1 and 1, respectively; the overall mean $D'$ values were – 0.705, – 0.857 and – 0.922, respectively.

Consistent with the findings of Garcia and Lohmueller (2021), neutral variants have a smaller magnitude of negative LD than selected variants as measured by $D'$. However, this effect is entirely due to the fact that the probability of case 2 increases as selection becomes stronger.

**9. Effects of sample size and allele frequency on the distribution of *D***

As described in the main text *Discussion: patterns of LD in population genetic data,* finite samples are likely to have a preponderance of negative values of the linkage disequilibrium coefficient $D$ for pairs of segregating sites with low variant frequencies, even if these sites are in linkage equilibrium (*D* = 0). This possibility was investigated by a Monte Carlo simulation, in which repeated samples were drawn from a population segregating for two loci, each with a frequency *q* of the rarer allele, and with a specified value of *D*. This was done by multinomial sampling of the four haplotypes into a sample of size *k*, retaining only samples in which both loci were segregating, and recording the value of *D* for each replicate.

Figure S1.2 shows histograms of the distribution of *D* values for 10000 replicate samples of size 1000 from a population with *D* = 0, and allele frequencies of 0.001 (left-hand panel) and 0.01 (right-hand panel), respectively. It can be seen that the former case yields only negative *D* values, reflecting the fact that none of the samples contains – – haplotypes; the latter case gives a predominance of negative *D* values, with a small proportion of positive ones; the magnitude of *D* is, however, very much larger for the positive values, leading to a mean *D* very close to zero in this case. If a much larger number of replicate simulations is used for the case of *q* = 0.001, the proportion of samples with – – present (where *D* is almost always positive) approaches the expected value of 0.001/0.4 = 0.0025, derived in the main text, and mean *D* is close to zero.

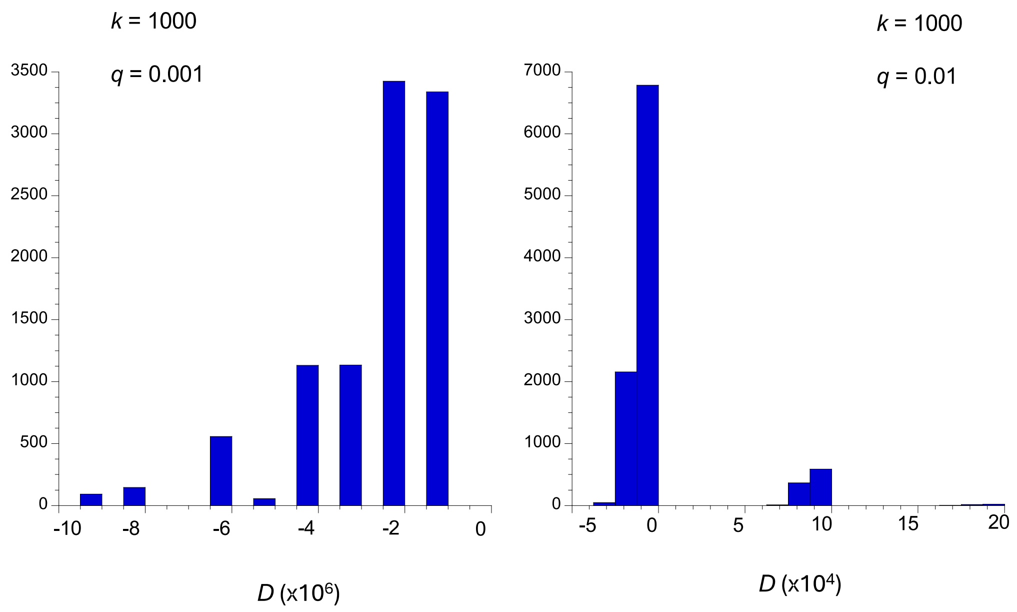
