## Supplementary File S2 for "A gene-based model of fitness and its implications for genetic variation: Linkage disequilibrium"

### Exact recursions for two-locus mutation model plus approximate LD results
### Additive fitnesses plus epistasis
### No recombination

### Results used in the relevant figure and table

#### Mutation rate= 1.0000000000000000E-004
#### Selection coefficient= 1.0000000000000000E-002
#### Maximum number of generations= 5000

##### Dominance coefficient=0

Initial state (approximate equilibrium equation; no LD)
x = 9.0000000000000011E-002 y= 1.0000000000000002E-002 q=
0.10000000000000001

**Sites Model**

Epistasis parameter= 0.0000000000000000

Iteration of haplotype frequencies

Population genetic statistics for sites model
Gen= 5000 x= 9.0002497580254484E-002 y= 9.9949999364381103E-003
q= 9.9997497516692596E-002

wx= 0.99900002502487173 wy= 0.99800005004974357 wz=
1.0000000000000000
wy*wz-wx**2= -9.9994995073071635E-007
wbar= 0.99980001000982344

D= -4.4995731628277857E-006 Correlation= -4.9996369502277469E-005
Approx. D= -5.0625000000000011E-006 Approx. Corr.=
-5.6250000000000012E-005

q relative to approx. equilib. value= 9.9997497516692596E-002

Load statistics for sites model

Relative L= 0.49997497548004716 Relative B= 0.94444997820969534

Relative V x h = 0.49497097762226605

Epistasis parameter=    8.0000000000000002E-002

Iteration of haplotype frequencies

Population genetic statistics for sites model
Gen=          5000  x=    9.0039276412929789E-002 y=    9.2546006895169036E-003
q=    9.9293877102446693E-002

wx=   0.99899965755236164       wy=   0.99785525226312310       wz=
1.0000000000000000
wy*wz-wx**2=   -1.4506352661269872E-004
wbar=   0.99980001099660254

D=   -6.0467334051889088E-004  Correlation=   -6.7610669667680636E-003
Approx. D=   -5.5862068965517261E-004  Approx. Corr.=
-6.2068965517241385E-003

q relative to approx. equilib. value=    9.9293877102446693E-002

Load statistics for sites model

Relative L=   0.49997251243407659       Relative B=   0.94485900596208205

Relative V x h =   0.50253809772371594

Epistasis parameter=   0.32000000000000001

Iteration of haplotype frequencies

Population genetic statistics for sites model
Gen=          5000  x=    9.0123061606239008E-002 y=    7.5718688645382308E-003
q=    9.7694930470777236E-002

wx=   0.99899882072461965       wy=   0.99742085386106261       wz=
1.0000000000000000
wy*wz-wx**2=   -5.7778994811819029E-004
wbar=   0.99980001236256588

D=   -1.9724305751517707E-003  Correlation=   -2.2375682988090836E-002
Approx. D=   -1.5804878048780488E-003  Approx. Corr.=
-1.7560975609756096E-002

q relative to approx. equilib. value=    9.7694930470777236E-002

Load statistics for sites model

Relative L= 0.49996910328702870 Relative B= 0.94579014434548525

Relative V x h = 0.52500066065532691

**Gene model**
Epistasis parameter= 0.0000000000000000

Iteration of haplotype frequencies

Population genetic statistics for gene model
Gen= 5000 x= 6.5569821689364233E-002 y= 6.8799756568001464E-003
q= 7.2449797346164385E-002
wx= 0.99858540393105988 wy= 0.99789530583644848 wz=
1.0000000000000000
wy*wz-wx**2= 7.2249689229042513E-004
wbar= 0.99980001013134634

D= 1.6310025212998569E-003 Correlation= 2.4270573173134285E-002
Approx. D= 2.0250000000000003E-003 Approx. Corr.=
2.2500000000000003E-002

q relative to approx. equilib. value= 0.72449797346164380

Load statistics for gene model

Relative L= 0.49997467144594293 Relative B= 0.65567017876494937

Relative V x h = 0.52617629609261840

Epistasis parameter= 8.0000000000000002E-002

Iteration of haplotype frequencies

Population genetic statistics for gene model
Gen= 5000 x= 6.5593889423364318E-002 y= 6.3557662112840281E-003
q= 7.1949655634648349E-002
wx= 0.99858515879822640 wy= 0.99773747343215213 wz=
1.0000000000000000
wy*wz-wx**2= 5.6515406007306979E-004
wbar= 0.99980001003531438

D= 1.1790132653395485E-003 Correlation= 1.7657061740091240E-002
Approx. D= 1.5390000000000002E-003 Approx. Corr.=
1.7100000000000001E-002

q relative to approx. equilib. value= 0.71949655634648346

Load statistics for gene model

Relative L= 0.49997491125393212 Relative B= 0.65585005502643379

Relative V x h = 0.53437968379769674

Epistasis parameter= 0.32000000000000001

Iteration of haplotype frequencies

Population genetic statistics for gene model
Gen= 5000 x= 6.5648325049297129E-002 y= 5.1728948914038283E-003
q= 7.0821219940700952E-002
wx= 0.99858461017804545 wy= 0.99726376189906618 wz=
1.0000000000000000
wy*wz-wx**2= 9.2538214627180260E-005
wbar= 0.99980000978540151

D= 1.5724969751468310E-004 Correlation= 2.3896105781266142E-003
Approx. D= 8.0999999999999800E-005 Approx. Corr.=
8.9999999999999770E-004

q relative to approx. equilib. value= 0.70821219940700952

Load statistics for gene model

Relative L= 0.49997553500540504 Relative B= 0.65625472197239498

Relative V x h = 0.55876730001052854

**Dominance coefficient= 1.0000000000000000E-002**

Initial state (approximate equilibrium equation; no LD)
x = 8.3329790490440522E-002 y= 8.4175799900921121E-003 q=
9.1747370480532636E-002

**Sites model**

Epistasis parameter= 0.0000000000000000

Iteration of haplotype frequencies

Population genetic statistics for sites model
Gen= 5000 x= 8.6818984413030972E-002 y= 9.2189435706268759E-003
q= 9.6037927983657845E-002

wx= 0.99893962073018283 wy= 0.99789844904576197 wz=

0.99998079241460369
wy*wz-wx**2= -1.0840384764287947E-006
wbar= 0.99978080847419593

D= -4.3400407673810371E-006 Correlation= -4.9992035315733395E-005
Approx. D= -4.1549977713789732E-006 Approx. Corr.=
-4.9862093099293565E-005

q relative to approx. equilib. value= 1.0467649097805545

Load statistics for sites model

Relative L= 0.54797882494197614 Relative B= 8.2111101734188660E-002

Relative V x h = 0.45594667533483452

Epistasis parameter= 2.0000000000000000E-002

Iteration of haplotype frequencies

Population genetic statistics for sites model
Gen= 5000 x= 8.6813619410393228E-002 y= 9.0349866504388938E-003
q= 9.5848606060832120E-002

wx= 0.99893934162859432 wy= 0.99786024157410247 wz=
0.99998079413906082
wy*wz-wx**2= -3.8731444303441442E-005
wbar= 0.99978081045347589

D= -1.5196863336569555E-004 Correlation= -1.7535857210555206E-003
Approx. D= -1.3762480092471653E-004 Approx. Corr.=
-1.6515678260406120E-003

q relative to approx. equilib. value= 1.0447013964413259

Load statistics for sites model

Relative L= 0.54797387774171558 Relative B= 8.2104066374778309E-002

Relative V x h = 0.45750037015922895

Epistasis parameter= 4.0000000000000001E-002

Iteration of haplotype frequencies

Population genetic statistics for sites model

Gen= 5000 x= 8.6807889737666760E-002 y= 8.8582099386164230E-003
q= 9.5666099676283187E-002

wx= 0.99893906583516268 wy= 0.99782204369974448 wz=
0.99998079591462219
wy*wz-wx**2= -7.6375811602136245E-005
wbar= 0.99978081245701678

D= -2.9379268865611929E-004 Correlation= -3.3958938332291021E-003
Approx. D= -2.6566732815880338E-004 Approx. Corr.=
-3.1881434790032304E-003

q relative to approx. equilib. value= 1.0427121690270353

Load statistics for sites model

Relative L= 0.54796886979634463 Relative B= 8.2096715826347366E-002

Relative V x h = 0.45905051394555696

Epistasis parameter= 8.0000000000000002E-002

Iteration of haplotype frequencies

Population genetic statistics for sites model
Gen= 5000 x= 8.6795416813629497E-002 y= 8.5245820798809310E-003
q= 9.5319998893510421E-002

wx= 0.99893852343544443 wy= 0.99774567717199547 wz=
0.99998079960717501
wy*wz-wx**2= -1.5165354033164213E-004
wbar= 0.99978081653577788

D= -5.6132010917790687E-004 Correlation= -6.5092594533889322E-003
Approx. D= -4.9674804346998486E-004 Approx. Corr.=
-5.9612299580540894E-003

q relative to approx. equilib. value= 1.0389398452976464

Load statistics for sites model

Relative L= 0.54795867445864654 Relative B= 8.2081140456726806E-002

Relative V x h = 0.46214084273168810

Epistasis parameter= 0.16000000000000000

Iteration of haplotype frequencies

Population genetic statistics for sites model
Gen= 5000 x= 8.6766906212620273E-002 y= 7.9272956360103841E-003
q= 9.4694201848630657E-002

wx= 0.99893747094637864 wy= 0.99759306360733169 wz=
0.99998080748648122
wy*wz-wx**2= -3.0215357177487689E-004
wbar= 0.99978082497064613

D= -1.0396962277388205E-003 Correlation= -1.2127960631428198E-002
Approx. D= -8.7905407692190731E-004 Approx. Corr.=
-1.0549097408600243E-002

q relative to approx. equilib. value= 1.0321189735756329

Load statistics for sites model

Relative L= 0.54793758960732220 Relative B= 8.2046924376897812E-002

Relative V x h = 0.46828586448047854

Epistasis parameter= 0.32000000000000001

Iteration of haplotype frequencies

Population genetic statistics for sites model
Gen= 5000 x= 8.6698686916528364E-002 y= 6.9525833444194027E-003
q= 9.3651270260947772E-002

wx= 0.99893546785012599 wy= 0.99728833044676590 wz=
0.99998082478097294
wy*wz-wx**2= -6.0286170435364106E-004
wbar= 0.99978084284693225

D= -1.8179770770696685E-003 Correlation= -2.1418023959839001E-002
Approx. D= -1.4289119736920286E-003 Approx. Corr.=
-1.7147672702428687E-002

q relative to approx. equilib. value= 1.0207515460164509

Load statistics for sites model

Relative L= 0.54789290141554892 Relative B= 8.1968912138300196E-002

Relative V x h = 0.48045942674156777

Epistasis parameter= 0.64000000000000001

Iteration of haplotype frequencies

Population genetic statistics for sites model
Gen= 5000 x= 8.6533175266205381E-002 y= 5.5795183956651130E-003
q= 9.2112693661870498E-002

wx= 0.99893173082947373 wy= 0.99668091667274239 wz=
0.99998086328329694
wy*wz-wx**2= -1.2027593855714613E-003
wbar= 0.99978088164949153

D= -2.9052299379804802E-003 Correlation= -3.4739941613750838E-002
Approx. D= -2.0791899503976607E-003 Approx. Corr.=
-2.4951340188910983E-002

q relative to approx. equilib. value= 1.0039818381652188

Load statistics for sites model

Relative L= 0.54779589637738169 Relative B= 8.1788226962350150E-002

Relative V x h = 0.50447374317768101

**Gene Model**

Epistasis parameter= 0.0000000000000000

Iteration of haplotype frequencies

Population genetic statistics for gene model
Gen= 5000 x= 6.4100613756416194E-002 y= 6.3747553975845054E-003
q= 7.0475369154000697E-002
wx= 0.99853582399120899 wy= 0.99777640167487303 wz=
0.99998590492615091
wy*wz-wx**2= 6.8854614900382138E-004
wbar= 0.99978591772737180

D= 1.4079777401918292E-003 Correlation= 2.1493024167138948E-002
Approx. D= 1.6438183055586984E-003 Approx. Corr.=
1.9726658328119460E-002

q relative to approx. equilib. value= 0.76814592924986846

Load statistics for gene model

Relative L= 0.53520568082283249 Relative B= 5.7738649732116787E-002

Relative V x h = 0.49604551501696220

Epistasis parameter= 2.0000000000000000E-002

Iteration of haplotype frequencies

Population genetic statistics for gene model
Gen= 5000 x= 6.4106686677979835E-002 y= 6.2447386617030540E-003
q= 7.0351425339682888E-002
wx= 0.99853576757509954 wy= 0.99773437217268801 wz=
0.99998590473595983
wy*wz-wx**2= 6.4662971647699852E-004
wbar= 0.99978591752613821

D= 1.2954156143780843E-003 Correlation= 1.9806941504530971E-002
Approx. D= 1.5371602500444464E-003 Approx. Corr.=
1.8446707245961300E-002

q relative to approx. equilib. value= 0.76679500427328717

Load statistics for gene model

Relative L= 0.53520618389011554 Relative B= 5.7742223077857743E-002

Relative V x h = 0.49789036280204035

Epistasis parameter= 4.0000000000000001E-002

Iteration of haplotype frequencies

Population genetic statistics for gene model
Gen= 5000 x= 6.4112520290512787E-002 y= 6.1199098627233127E-003
q= 7.0232430153236103E-002
wx= 0.99853571349431480 wy= 0.99769233899027754 wz=
0.99998590455467329
wy*wz-wx**2= 6.0470494886011306E-004
wbar= 0.99978591733243194

D= 1.1873156174941335E-003 Correlation= 1.8182520690661452E-002
Approx. D= 1.4305021945301940E-003 Approx. Corr.=
1.7166756163803136E-002

q relative to approx. equilib. value= 0.76549801684112928

Load statistics for gene model

Relative L= 0.53520666812875883 Relative B= 5.7745651995043695E-002

Relative V x h = 0.49973234281631868

Epistasis parameter= 8.0000000000000002E-002

Iteration of haplotype frequencies

Population genetic statistics for gene model
Gen= 5000 x= 6.4123523766667884E-002 y= 5.8846236376929959E-003
q= 7.0008147404360885E-002
wx= 0.99853561178767802 wy= 0.99760826239889866 wz=
0.99998590421652400
wy*wz-wx**2= 5.2083232064559226E-004
wbar= 0.99978591696704877

D= 9.8348293470228165E-004 Correlation= 1.5105638924751618E-002
Approx. D= 1.2171860835016896E-003 Approx. Corr.=
1.4606853999486814E-002

q relative to approx. equilib. value= 0.76305344815539455

Load statistics for gene model

Relative L= 0.53520758150865821 Relative B= 5.7752109999300867E-002

Relative V x h = 0.50340834887455743

Epistasis parameter= 0.16000000000000000

Iteration of haplotype frequencies

Population genetic statistics for gene model
Gen= 5000 x= 6.4143202344941072E-002 y= 5.4643755905604257E-003
q= 6.9607577935501500E-002
wx= 0.99853543089439001 wy= 0.99744007322355832 wz=
0.99998590362437612
wy*wz-wx**2= 3.5300618217870561E-004
wbar= 0.99978591631903058

D= 6.1916068451350892E-004 Correlation= 9.5605017412954546E-003
Approx. D= 7.9055386144468028E-004 Approx. Corr.=
9.4870496708541648E-003

q relative to approx. equilib. value=   0.75868744325780046

Load statistics for gene model

Relative L=   0.53520920134851646       Relative B=    5.7763627610961252E-002

Relative V x h =   0.51073245199116746

Epistasis parameter=   0.32000000000000001

Iteration of haplotype frequencies

Population genetic statistics for gene model
Gen=         5000  x=    6.4175258760671788E-002 y=    4.7812604065305374E-003
q=    6.8956519167202329E-002
wx=   0.99853513904450564       wy=   0.99710358126396281       wz=
0.99998590269547882
wy*wz-wx**2=    1.7100884508347569E-005
wbar=   0.99978591530043937

D=    2.6258870873790974E-005  Correlation=    4.0900701171102242E-004
Approx. D=   -6.2710582669338140E-005  Approx. Corr.=
-7.5255898641113481E-004

q relative to approx. equilib. value=   0.75159123150928697

Load statistics for gene model

Relative L=   0.53521174741881439       Relative B=    5.7782300590301768E-002

Relative V x h =   0.52529307075644527

Epistasis parameter=   0.64000000000000001

Iteration of haplotype frequencies

Population genetic statistics for gene model
Gen=         5000  x=    6.4220302045492825E-002 y=    3.8246089257052305E-003
q=    6.8044910971198053E-002
wx=   0.99853473509814994       wy=   0.99643029695006602       wz=
0.99998590146783584
wy*wz-wx**2=   -6.5536845205738103E-004
wbar=   0.99978591398551075

D=   -8.0550098337303644E-004  Correlation=   -1.2702097458036410E-002

Approx. D= -1.7692394708973751E-003 Approx. Corr.=
-2.1231776300941734E-002

q relative to approx. equilib. value= 0.74165516259276465

Load statistics for gene model

Relative L= 0.53521503426729278 Relative B= 5.7808346506753629E-002

Relative V x h = 0.55418519611037287

#### Dominance coefficient= 5.0000000000000003E-002

Initial state (approximate equilibrium equation; no LD)
x = 7.2564392256023791E-002 y= 6.2045455655368922E-003 q=
7.8768937821560678E-002

**Sites Model**

Epistasis parameter= 0.0000000000000000

Epistasis parameter= 0.0000000000000000

Iteration of haplotype frequencies

Population genetic statistics for sites model
Gen= 5000 x= 7.4629644542442919E-002 y= 6.5934616996254879E-003
q= 8.1223106242068410E-002

wx= 0.99868776894299172 wy= 0.99745676099168412 wz=
0.99991877689429920
wy*wz-wx**2= -1.5153805762446737E-006
wbar= 0.99971880431640392

D= -3.7312879848372393E-006 Correlation= -4.9999897115019031E-005
Approx. D= -3.5068039328963098E-006 Approx. Corr.=
-4.8326787062771820E-005

q relative to approx. equilib. value= 1.0311565508991283

Load statistics for sites model

Relative L= 0.70298921565280703 Relative B= 0.35481439488444183

Relative V x h = 0.33723513405099204

Epistasis parameter= 8.0000000000000002E-002

Iteration of haplotype frequencies

Population genetic statistics for sites model
Gen= 5000 x= 7.4438138402031506E-002 y= 6.0694081163267053E-003
q= 8.0507546518358217E-002

wx= 0.99868387118669288 wy= 0.99726798515708037 wz=
0.99991900690140467
wy*wz-wx**2= -1.8226123560638463E-004
wbar= 0.99971903436337184

D= -4.1205693007889677E-004 Correlation= -5.5663750111508885E-003
Approx. D= -4.1860584885762515E-004 Approx. Corr.=
-5.7687501520124066E-003

q relative to approx. equilib. value= 1.0220722628091812

Load statistics for sites model

Relative L= 0.70241409863415516 Relative B= 0.35370742673424904

Relative V x h = 0.34016526712556799

Epistasis parameter= 0.32000000000000001

Iteration of haplotype frequencies

Population genetic statistics for sites model
Gen= 5000 x= 7.3854774268140549E-002 y= 4.8958165802818919E-003
q= 7.8750590848422439E-002

wx= 0.99867241062522838 wy= 0.99670493519387704 wz=
0.99991968274834953
wy*wz-wx**2= -7.2170115122893730E-004
wbar= 0.99971970993301640

D= -1.3058389786937408E-003 Correlation= -1.7999423058158815E-002
Approx. D= -1.2870435846130772E-003 Approx. Corr.=
-1.7736572230524481E-002

q relative to approx. equilib. value= 0.99976707857633174

Load statistics for sites model

Relative L= 0.70072517372701359 Relative B= 0.35039727631436846

Relative V x h =   0.34892656713648867

**Gene Model**

Epistasis parameter=    0.0000000000000000
Iteration of haplotype frequencies

Population genetic statistics for gene model
Gen=         5000  x=    5.8168048486129560E-002 y=    4.7329589872517515E-003
q=    6.2901007473381315E-002
wx=   0.99832617917178135       wy=   0.99728136841929804       wz=
0.99993709899242644
wy*wz-wx**2=    5.6347839666248678E-004
wbar=   0.99973712140378046

D=    7.7642224608538007E-004  Correlation=    1.3172096321174392E-002
Approx. D=    9.7056710462041745E-004  Approx. Corr.=
1.3375252991798437E-002

q relative to approx. equilib. value=   0.79855091629995312

Load statistics for gene model

Relative L=   0.65719648908360506       Relative B=   0.26328109063861238

Relative V x h =   0.39908147027233676

Epistasis parameter=    8.0000000000000002E-002

Iteration of haplotype frequencies

Population genetic statistics for gene model
Gen=         5000  x=    5.8184519385820407E-002 y=    4.3607518544416671E-003
q=    6.2545271240262074E-002
wx=   0.99832603032576517       wy=   0.99707101750301130       wz=
0.99993710586850926
wy*wz-wx**2=    3.5344476133025715E-004
wbar=   0.99973712825574701

D=    4.4884089992371090E-004  Correlation=    7.6550430972457889E-003
Approx. D=    6.1852514897315085E-004  Approx. Corr.=
8.5238107802357473E-003

q relative to approx. equilib. value=   0.79403471685690485

Load statistics for gene model

Relative L= 0.65717935908340275 Relative B= 0.26330964623451947

Relative V x h = 0.40398298232959323

Epistasis parameter= 0.32000000000000001

Iteration of haplotype frequencies

Population genetic statistics for gene model
Gen= 5000 x= 5.8221474201282222E-002 y= 3.5280173200197208E-003 q= 6.1749491521301943E-002
wx= 0.99832570125378728 wy= 0.99643965163516601 wz= 0.99993712154282177
wy*wz-wx**2= -2.7720873666614576E-004
wbar= 0.99973714387847967

D= -2.8498238311962243E-004 Correlation= -4.9188753784757617E-003
Approx. D= -4.3760071796864868E-004 Approx. Corr.= -6.0305158544523216E-003

q relative to approx. equilib. value= 0.78393200707093746

Load statistics for gene model

Relative L= 0.65714030199716911 Relative B= 0.26337309838200368

Relative V x h = 0.41855851211835904

#### Dominance coefficient= 0.10000000000000001

Initial state (approximate equilibrium equation; no LD)
x = 6.0308544485159571E-002 y= 4.1556297612746311E-003 q= 6.4464174246434205E-002

**Sites model**

Epistasis parameter= 0.0000000000000000

Iteration of haplotype frequencies

Population genetic statistics for sites model
Gen= 5000 x= 6.1280562132483174E-002 y= 4.2974058856415530E-003 q= 6.5577968018124724E-002

wx= 0.99834422032129755 wy= 0.99681959657833552 wz= 0.99986884406425947
wy*wz-wx**2= -2.3244775576580778E-006
wbar= 0.99966888061059922

D= -3.0640037446405533E-006 Correlation= -5.0002102536943590E-005
Approx. D= -2.8828799798139634E-006 Approx. Corr.=
-4.7802181339716600E-005

q relative to approx. equilib. value= 1.0172777171926952

Load statistics for sites model

Relative L= 0.82779847575669485 Relative B= 0.58591619450212573

Relative V x h = 0.25883662106968530

Epistasis parameter= 8.0000000000000002E-002

Iteration of haplotype frequencies

Population genetic statistics for sites model
Gen= 5000 x= 6.1015456697885613E-002 y= 3.9425769837240802E-003
q= 6.4958033681609698E-002

wx= 0.99833718772353541 wy= 0.99657721582744929 wz=
0.99986945312054976
wy*wz-wx**2= -2.3002460994581142E-004
wbar= 0.99966948947596346

D= -2.7696915605705114E-004 Correlation= -4.5600272128329131E-003
Approx. D= -3.0859463547351114E-004 Approx. Corr.=
-5.1169305793716934E-003

q relative to approx. equilib. value= 1.0076609906346983

Load statistics for sites model

Relative L= 0.82627631185528039 Relative B= 0.58273652539988752

Relative V x h = 0.26063225855030359

Epistasis parameter= 0.32000000000000001

Iteration of haplotype frequencies

Population genetic statistics for sites model
Gen= 5000 x= 6.0234894722479448E-002 y= 3.1528431334423425E-003
q= 6.3387737855921791E-002

wx= 0.99831647428118697 wy= 0.99585390734897794 wz= 0.99987120670472018
wy*wz-wx**2= -9.1013477858670200E-004
wbar= 0.99967124250238315

D= -8.6516217704871862E-004 Correlation= -1.4572445250392222E-002
Approx. D= -1.0221073853373011E-003 Approx. Corr.= -1.6947969712464477E-002

q relative to approx. equilib. value= 0.98330178889133979

Load statistics for sites model

Relative L= 0.82189374433227280 Relative B= 0.57347370379882423

Relative V x h = 0.26640941091105164

**Gene Model**

Epistasis parameter= 0.0000000000000000
Iteration of haplotype frequencies

Population genetic statistics for gene model
Gen= 5000 x= 5.0970036673662268E-002 y= 3.3101555948458075E-003 q= 5.4280192268508075E-002
wx= 0.99803619716036096 wy= 0.99661519624409645 wz= 0.99989143961532512
wy*wz-wx**2= 4.3075247270452088E-004
wbar= 0.99969147118781387

D= 3.6381632213960263E-004 Correlation= 7.0872592043569353E-003
Approx. D= 4.5803636630052679E-004 Approx. Corr.= 7.5948834482854758E-003

q relative to approx. equilib. value= 0.84202105903048230

Load statistics for gene model

Relative L= 0.77132202921650073 Relative B= 0.46498343397762909

Relative V x h = 0.32362157876727704

Epistasis parameter= 8.0000000000000002E-002

Iteration of haplotype frequencies

Population genetic statistics for gene model

Gen= 5000 x= 5.0979248282607546E-002 y= 3.0473041073775242E-003 q= 5.4026552389985071E-002
wx= 0.99803605450727328 wy= 0.99634926311453542 wz= 0.99989145932642398
wy*wz-wx**2= 1.6515259795502057E-004
wbar= 0.99969149089122489

D= 1.2843574422972265E-004 Correlation= 2.5130417602621060E-003
Approx. D= 1.9476713247735581E-004 Approx. Corr.= 3.2295114090389486E-003

q relative to approx. equilib. value= 0.83808647239398271

Load statistics for gene model

Relative L= 0.77127277067269118 Relative B= 0.46499172398259481

Relative V x h = 0.32675417447033239

Epistasis parameter= 0.32000000000000001

Iteration of haplotype frequencies

Population genetic statistics for gene model
Gen= 5000 x= 5.0999834789263739E-002 y= 2.4608944441975823E-003 q= 5.3460729233461318E-002
wx= 0.99803573815698321 wy= 0.99555121481570119 wz= 0.99989150356894596
wy*wz-wx**2= -6.3213357659219760E-004
wbar= 0.99969153511736919

D= -3.9715512597588570E-004 Correlation= -7.8484998940485159E-003
Approx. D= -5.9504056899215691E-004 Approx. Corr.= -9.8666047087006322E-003

q relative to approx. equilib. value= 0.82930914509338438

Load statistics for gene model

Relative L= 0.77116220526620238 Relative B= 0.46500977310765457

Relative V x h = 0.33608124246679538

**Dominance coefficient= 0.14999999999999999**

Initial state (approximate equilibrium equation; no LD)
x = 5.0088780211284531E-002 y= 2.7968946002739068E-003 q=

5.2885674811558439E-002

**Sites Model**

Epistasis parameter=    0.0000000000000000

Iteration of haplotype frequencies

Population genetic statistics for sites model
Gen=         5000  x=    5.0518392278034764E-002 y=    2.8451407389593433E-003
q=    5.3363533016994105E-002

wx=   0.99796636467007316        wy=   0.99609281993912446        wz=
0.99983990940102196
wy*wz-wx**2=   -3.5101698587780561E-006
wbar=   0.99963995146889539

D=   -2.5259170964783695E-006  Correlation=   -5.0002450288203927E-005
Approx. D=   -2.3962289015962961E-006  Approx. Corr.=
-4.7839633776037700E-005

q relative to approx. equilib. value=    1.0090356832381995

Load statistics for sites model

Relative L=   0.90012132821686519        Relative B=   0.72725434463951000

Relative V x h =   0.23409423817010389

Epistasis parameter=    8.0000000000000002E-002

Iteration of haplotype frequencies

Population genetic statistics for sites model
Gen=         5000  x=    5.0265584416498865E-002 y=    2.6071491411800299E-003
q=    5.2872733557678894E-002

wx=   0.99795681034706252        wy=   0.99578925661207562        wz=
0.99984075608356515
wy*wz-wx**2=   -2.8711208717391301E-004
wbar=   0.99964079795332916

D=   -1.8837681268127349E-004  Correlation=   -3.7617275741811279E-003
Approx. D=   -2.1784894488730567E-004  Approx. Corr.=
-4.3492563398105340E-003

q relative to approx. equilib. value=   0.99975529755598924

Load statistics for sites model

Relative L= 0.89800511687807594 Relative B= 0.72242877032516917

Relative V x h = 0.23591418320511715

Epistasis parameter= 0.32000000000000001

Iteration of haplotype frequencies

Population genetic statistics for sites model
Gen= 5000 x= 4.9530509067140807E-002 y= 2.0781161467316382E-003
q= 5.1608625213872446E-002

wx= 0.99792871699152652 wy= 0.99488190244977659 wz=
0.99984317913279974
wy*wz-wx**2= -1.1358399892816839E-003
wbar= 0.99964322045464860

D= -5.8533404973431374E-004 Correlation= -1.1958973474426232E-002
Approx. D= -7.6349153101859177E-004 Approx. Corr.=
-1.5242765501536096E-002

q relative to approx. equilib. value= 0.97585263680124423

Load statistics for sites model

Relative L= 0.89194886301015663 Relative B= 0.70849475279529772

Relative V x h = 0.24204256688858605

**Gene Model**

Epistasis parameter= 0.0000000000000000

Iteration of haplotype frequencies

Population genetic statistics for gene model
Gen= 5000 x= 4.4404852964548693E-002 y= 2.3555205550352056E-003
q= 4.6760373519583898E-002
wx= 0.99771331797165863 wy= 0.99589423967878565 wz=
0.99985971887935954
wy*wz-wx**2= 3.2266966078808856E-004
wbar= 0.99965975687805730

D= 1.6898802334420343E-004 Correlation= 3.7911927623671426E-003
Approx. D= 2.0351865140821277E-004 Approx. Corr.=

4.0631584668209177E-003

q relative to approx. equilib. value=   0.88417844125464717

Load statistics for gene model

Relative L=   0.85060780427760085       Relative B=   0.61222407931877754

Relative V x h =   0.28794609162238849

Epistasis parameter=    8.0000000000000002E-002

Iteration of haplotype frequencies

Population genetic statistics for gene model
Gen=         5000  x=    4.4409544834406470E-002 y=    2.1685054249069710E-003
q=    4.6578050259313442E-002
wx=   0.99771316070761695       wy=   0.99556909083761258       wz=
0.99985974540783862
wy*wz-wx**2=   -2.0933483744212111E-006
wbar=   0.99965978340104034

D=   -1.0093410521549995E-006  Correlation=   -2.2728537226949223E-005
Approx. D=    1.2283526833019575E-005  Approx. Corr.=
2.4523509618731365E-004

q relative to approx. equilib. value=   0.88073094321439138

Load statistics for gene model

Relative L=   0.85054149682084912       Relative B=   0.61220802226723803

Relative V x h =   0.29010908331214463

Epistasis parameter=   0.32000000000000001

Iteration of haplotype frequencies

Population genetic statistics for gene model
Gen=         5000  x=    4.4420021136616805E-002 y=    1.7513095352214318E-003
q=    4.6171330671838234E-002
wx=   0.99771281067572770       wy=   0.99459347434359224       wz=
0.99985980475074909
wy*wz-wx**2=   -9.7681552290718177E-004
wbar=   0.99965984273159003

D= -3.8048224078680165E-004 Correlation= -8.6395600482281291E-003
Approx. D= -5.6142184689255998E-004 Approx. Corr.=
-1.1208535015713499E-002

q relative to approx. equilib. value= 0.87304039962343927

Load statistics for gene model

Relative L= 0.85039317044445695 Relative B= 0.61217189784313242

Relative V x h = 0.29655428293419822

#### Dominance coefficient= 0.20000000000000001

Initial state (approximate equilibrium equation; no LD)
x = 4.2010431875004424E-002 y= 1.9308349282771775E-003 q=
4.3941266803281601E-002

**Sites model**

Epistasis parameter= 0.0000000000000000

Iteration of haplotype frequencies

Population genetic statistics for sites model
Gen= 5000 x= 4.2196402364830465E-002 y= 1.9464846544900182E-003
q= 4.4142887019320481E-002

wx= 0.99755857112983404 wy= 0.99529371380773446 wz=
0.99982342845193362
wy*wz-wx**2= -5.1295786894778317E-006
wbar= 0.99962347377017735

D= -2.1098199104729876E-006 Correlation= -5.0002495200669045E-005
Approx. D= -2.0181866531900346E-006 Approx. Corr.=
-4.8040131060657465E-005

q relative to approx. equilib. value= 1.0045884024450524

Load statistics for sites model

Relative L= 0.94131557461842441 Relative B= 0.81144443999023286

Relative V x h = 0.24136086506169621

Epistasis parameter= 2.0000000000000000E-002

Iteration of haplotype frequencies

Population genetic statistics for sites model
Gen= 5000 x= 4.2143060279759482E-002 y= 1.9032365920859445E-003
q= 4.4046296871845428E-002

wx= 0.99755570880969646 wy= 0.99520116443508289 wz=
0.99982366255359301
wy*wz-wx**2= -9.1718975730792351E-005
wbar= 0.99962370782335430

D= -3.6839676036799773E-005 Correlation= -8.7492241071721153E-004
Approx. D= -3.9422851990821118E-005 Approx. Corr.=
-9.3840625366855906E-004

q relative to approx. equilib. value= 1.0023902376104457

Load statistics for sites model

Relative L= 0.94073044165812081 Relative B= 0.80997007028536339

Relative V x h = 0.24185966258036484

Epistasis parameter= 4.0000000000000001E-002

Iteration of haplotype frequencies

Population genetic statistics for sites model
Gen= 5000 x= 4.2089951307655478E-002 y= 1.8617883803633776E-003
q= 4.3951739688018854E-002

wx= 0.99755285455249998 wy= 0.99510864305161084 wz=
0.99982389515511172
wy*wz-wx**2= -1.7829802746194279E-004
wbar= 0.99962394037677715

D= -6.9967041240001637E-005 Correlation= -1.6650896582063425E-003
Approx. D= -7.8192944282019628E-005 Approx. Corr.=
-1.8612744690335655E-003

q relative to approx. equilib. value= 1.0002383382523798

Load statistics for sites model

Relative L= 0.94014905808408045 Relative B= 0.80850333679209430

Relative V x h = 0.24236752474624687

Epistasis parameter=    8.0000000000000002E-002

Iteration of haplotype frequencies

Population genetic statistics for sites model
Gen=         5000  x=    4.1984430242360954E-002 y=    1.7838619972563282E-003
q=    4.3768292239617285E-002

wx=   0.99754716955240663       wy=   0.99492368391009567       wz=
0.99982435599520159
wy*wz-wx**2=   -3.5142395223297651E-004
wbar=   0.99962440112179418

D=   -1.3180140831621501E-004  Correlation=   -3.1491787251605578E-003
Approx. D=   -1.5383864346063840E-004  Approx. Corr.=
-3.6619153051880449E-003

q relative to approx. equilib. value=   0.99606350530496313

Load statistics for sites model

Relative L=   0.93899719550962280       Relative B=   0.80559239368956392

Relative V x h =   0.24341026988284503

Epistasis parameter=   0.16000000000000000

Iteration of haplotype frequencies

Population genetic statistics for sites model
Gen=         5000  x=    4.1776153105439022E-002 y=    1.6453255146875612E-003
q=    4.3421478620126581E-002

wx=   0.99753588945079585       wy=   0.99455409735676337       wz=
0.99982526107717984
wy*wz-wx**2=   -6.9754069728555912E-004
wbar=   0.99962530601784860

D=   -2.4009929087055043E-004  Correlation=   -5.7805031730370432E-003
Approx. D=   -2.9797047958649719E-004  Approx. Corr.=
-7.0927735395118650E-003

q relative to approx. equilib. value=   0.98817084210425632

Load statistics for sites model

Relative L=   0.93673495531964313       Relative B=   0.79985809325216473

Relative V x h =   0.24560256213005840

Epistasis parameter=   0.32000000000000001

Iteration of haplotype frequencies

Population genetic statistics for sites model
Gen=         5000  x=    4.1370424510637785E-002 y=    1.4219431744176628E-003
q=    4.2792367685055448E-002

wx=   0.99751366191792046       wy=   0.99381622519437018       wz=
0.99982701044197309
wy*wz-wx**2=   -1.3892003480854287E-003
wbar=   0.99962705502592819

D=   -4.0924355767531229E-004  Correlation=   -9.9910097354191200E-003
Approx. D=   -5.6057046096921281E-004  Approx. Corr.=
-1.3343601480629953E-002

q relative to approx. equilib. value=   0.97385375520988959

Load statistics for sites model

Relative L=   0.93236243504449101       Relative B=   0.78872194349545399

Relative V x h =   0.25040347891570924

Epistasis parameter=   0.64000000000000001

Iteration of haplotype frequencies

Population genetic statistics for sites model
Gen=         5000  x=    4.0599984061472964E-002 y=    1.1130749760996557E-003
q=    4.1713059037572622E-002

wx=   0.99747038503450292       wy=   0.99234544933063273       wz=
0.99983029829187631
wy*wz-wx**2=   -2.7701224080469711E-003
wbar=   0.99963034221248537

D=   -6.2690431817236103E-004  Correlation=   -1.5683162823535726E-002
Approx. D=   -1.0021777112690243E-003  Approx. Corr.=
-2.3855448909710086E-002

q relative to approx. equilib. value=   0.94929122604306504

Load statistics for sites model

Relative L=   0.92414446858314103       Relative B=   0.76766053900706199

Relative V x h =   0.26158087779600692

**Gene Model**

Epistasis parameter=    0.0000000000000000

Iteration of haplotype frequencies

Population genetic statistics for gene model
Gen=         5000  x=    3.8691893966057288E-002 y=    1.7106720664878633E-003
q=    4.0402566032545148E-002
wx=   0.99735869095956986       wy=   0.99512140757953449       wz=
0.99983838973584216
wy*wz-wx**2=    2.3622731339967284E-004
wbar=   0.99963843204195635

D=    7.8304724473690634E-005  Correlation=    2.0197142939279121E-003
Approx. D=    8.8526257284252984E-005  Approx. Corr.=
2.1072446374188498E-003

q relative to approx. equilib. value=   0.91946748402637823

Load statistics for gene model

Relative L=   0.90391989493796410       Relative B=   0.71562775268025802

Relative V x h =   0.28103640325658430

Epistasis parameter=    2.0000000000000000E-002

Iteration of haplotype frequencies

Population genetic statistics for gene model
Gen=         5000  x=    3.8692502818259950E-002 y=    1.6747328616350666E-003
q=    4.0367235679895017E-002
wx=   0.99735864600773938       wy=   0.99502440859630525       wz=
0.99983839707862388
wy*wz-wx**2=    1.3934097864432005E-004
wbar=   0.99963843938328534

D=    4.5219145198879818E-005  Correlation=    1.1673155494671291E-003

Approx. D= 5.3842436291477905E-005 Approx. Corr.=
1.2816444365932201E-003

q relative to approx. equilib. value= 0.91866344820264334

Load statistics for gene model

Relative L= 0.90390154161615677 Relative B= 0.71561954084103319

Relative V x h = 0.28144269212947937

Epistasis parameter= 4.0000000000000001E-002

Iteration of haplotype frequencies

Population genetic statistics for gene model
Gen= 5000 x= 3.8693086661572014E-002 y= 1.6402722271581182E-003
q= 4.0333358888730131E-002
wx= 0.99735860291101686 wy= 0.99492740804716961 wz=
0.99983840412086111
wy*wz-wx**2= 4.2449077371498056E-005
wbar= 0.99963844642412769

D= 1.3492387911012704E-005 Correlation= 3.4858125523735511E-004
Approx. D= 1.9158615298702822E-005 Approx. Corr.=
4.5604423576759063E-004

q relative to approx. equilib. value= 0.91789249202342005

Load statistics for gene model

Relative L= 0.90388393951024448 Relative B= 0.71561166364481188

Relative V x h = 0.28184853178470970

Epistasis parameter= 8.0000000000000002E-002

Iteration of haplotype frequencies

Population genetic statistics for gene model
Gen= 5000 x= 3.8694185240309813E-002 y= 1.5754362087496187E-003
q= 4.0269621449059434E-002
wx= 0.99735852184188223 wy= 0.99473340262084398 wz=
0.99983841737458945
wy*wz-wx**2= -1.5135010455935571E-004
wbar= 0.99963845967522780

D=   -4.6206202900925253E-005  Correlation=   -1.1955658239257599E-003
Approx. D=   -5.0209026686847332E-005  Approx. Corr.=
-1.1951561658836683E-003

q relative to approx. equilib. value=   0.91644197763665836

Load statistics for gene model

Relative L=   0.90385081176047266       Relative B=   0.71559683458900747

Relative V x h =   0.28265896966181236

Epistasis parameter=   0.16000000000000000

Iteration of haplotype frequencies

Population genetic statistics for gene model
Gen=         5000  x=    3.8696141340087239E-002 y=    1.4600111342613770E-003
q=    4.0156152474348616E-002
wx=   0.99735837756698265       wy=   0.99434537666617906       wz=
0.99983844098294927
wy*wz-wx**2=   -5.3900209852797420E-004
wbar=   0.99963848327889837

D=   -1.5250544728175747E-004  Correlation=   -3.9566959304486371E-003
Approx. D=   -1.8894431065794763E-004  Approx. Corr.=
-4.4975569691861859E-003

q relative to approx. equilib. value=   0.91385969034806469

Load statistics for gene model

Relative L=   0.90379180258449276       Relative B=   0.71557040753654444

Relative V x h =   0.28427551460065231

Epistasis parameter=   0.32000000000000001

Iteration of haplotype frequencies

Population genetic statistics for gene model
Gen=         5000  x=    3.8699304612151035E-002 y=    1.2734083338568245E-003
q=    3.9972712946007863E-002
wx=   0.99735814445775484       wy=   0.99356927780985982       wz=
0.99983847918552138

wy*wz-wx**2= -1.3144726253486727E-003
wbar= 0.99963852147386933

D= -3.2440944640711733E-004 Correlation= -8.4536894499898877E-003
Approx. D= -4.6641487860014827E-004 Approx. Corr.=
-1.1102358575791222E-002

q relative to approx. equilib. value= 0.90968503764262532

Load statistics for gene model

Relative L= 0.90369631515790849 Relative B= 0.71552760963790063

Relative V x h = 0.28749514654354491

Epistasis parameter= 0.64000000000000001

Iteration of haplotype frequencies

Population genetic statistics for gene model
Gen= 5000 x= 3.8703701353254273E-002 y= 1.0141553688086534E-003
q= 3.9717856722062926E-002
wx= 0.99735782086690505 wy= 0.99201695901805964 wz=
0.99983853233534048
wy*wz-wx**2= -2.8658424879971500E-003
wbar= 0.99963857461309868

D= -5.6335277378567056E-004 Correlation= -1.4770519797738031E-002
Approx. D= -1.0213560144845494E-003 Approx. Corr.=
-2.4311961789001295E-002

q relative to approx. equilib. value= 0.90388510872646799

Load statistics for gene model

Relative L= 0.90356346708507285 Relative B= 0.71546799645781956

Relative V x h = 0.29389972166495243

#### Dominance coefficient= 0.25000000000000000

Initial state (approximate equilibrium equation; no LD)
x = 3.5756483418224190E-002 y= 1.3790476691406398E-003 q=
3.7135531087364830E-002

**Sites Model**

Epistasis parameter=    0.0000000000000000

Iteration of haplotype frequencies

Population genetic statistics for sites model
Gen=         5000  x=    3.5836382449501811E-002 y=    1.3835300687584811E-003
q=    3.7219912518260290E-002

wx=   0.99712780087481967       wy=   0.99444170131222953       wz=
0.99981390043740981
wy*wz-wx**2=   -7.2151308601053898E-006
wbar=   0.99961394765594114

D=   -1.7918191084695390E-006  Correlation=   -5.0002499734147821E-005
Approx. D=   -1.7257157411175024E-006  Approx. Corr.=
-4.8263016274076544E-005

q relative to approx. equilib. value=    1.0022722559345374

Load statistics for sites model

Relative L=   0.96513086015318073       Relative B=   0.86127651404087169

Relative V x h =   0.26657446556995229

Epistasis parameter=    8.0000000000000002E-002

Iteration of haplotype frequencies

Population genetic statistics for sites model
Gen=         5000  x=    3.5666377571390191E-002 y=    1.2687864595634583E-003
q=    3.6935164030953649E-002

wx=   0.99711511302220002       wy=   0.99400165034269494       wz=
0.99981481666526029
wy*wz-wx**2=   -4.2097081492697974E-004
wbar=   0.99961486369961339

D=   -9.5419882429996494E-005  Correlation=   -2.6825221649742375E-003
Approx. D=   -1.1103797683422219E-004  Approx. Corr.=
-3.1053942171961153E-003

q relative to approx. equilib. value=   0.99460443810700327

Load statistics for sites model

Relative L=   0.96284075096063992       Relative B=   0.85473536201708300

Relative V x h = 0.26881306902180035

Epistasis parameter= 0.32000000000000001

Iteration of haplotype frequencies

Population genetic statistics for sites model
Gen= 5000 x= 3.5173830772021061E-002 y= 1.0129080574336007E-003
q= 3.6186738829454661E-002

wx= 0.99707780285022873 wy= 0.99268350257115689 wz=
0.99981744565295572
wy*wz-wx**2= -1.6618610541162537E-003
wbar= 0.99961749215988349

D= -2.9657200967755926E-004 Correlation= -8.5033061714633370E-003
Approx. D= -4.1523301037748331E-004 Approx. Corr.=
-1.1612803348716595E-002

q relative to approx. equilib. value= 0.97445055368460876

Load statistics for sites model

Relative L= 0.95626960026330199 Relative B= 0.83584790423306010

Relative V x h = 0.27649869799675686

**Gene Model**

Epistasis parameter= 0.0000000000000000

Iteration of haplotype frequencies

Population genetic statistics for gene model
Gen= 5000 x= 3.3869790269793226E-002 y= 1.2710080758988232E-003
q= 3.5140798345692052E-002
wx= 0.99697606554498597 wy= 0.99430353907344138 wz=
0.99982429600826539
wy*wz-wx**2= 1.6756070307022419E-004
wbar= 0.99962434114474652

D= 3.6132367526232889E-005 Correlation= 1.0656653680458420E-003
Approx. D= 3.8141956065066863E-005 Approx. Corr.=
1.0667144086553785E-003

q relative to approx. equilib. value= 0.94628506222302355

Load statistics for gene model

Relative L= 0.93914713809865824 Relative B= 0.78611759485957100

Relative V x h = 0.29375709117783705

Epistasis parameter= 8.0000000000000002E-002

Iteration of haplotype frequencies

Population genetic statistics for gene model
Gen= 5000 x= 3.3870898012497437E-002 y= 1.1710929540311396E-003
q= 3.5041990966528574E-002
wx= 0.99697589255632402 wy= 0.99384941689704875 wz=
0.99982432160797963
wy*wz-wx**2= -2.8611130890099368E-004
wbar= 0.99962436673935573

D= -5.6848176867137223E-005 Correlation= -1.6811998900893429E-003
Approx. D= -6.4140132449952447E-005 Approx. Corr.=
-1.7938042648025566E-003

q relative to approx. equilib. value= 0.94362433875225848

Load statistics for gene model

Relative L= 0.93908315157577427 Relative B= 0.78608127637822411

Relative V x h = 0.29507225917871172

Epistasis parameter= 0.32000000000000001

Iteration of haplotype frequencies

Population genetic statistics for gene model
Gen= 5000 x= 3.3873376143475957E-002 y= 9.4760770567965604E-004
q= 3.4820983849155614E-002
wx= 0.99697550574452143 wy= 0.99248698023720838 wz=
0.99982437890841902
wy*wz-wx**2= -1.6474804641850582E-003
wbar= 0.99962442402836005

D= -2.6489321054350146E-004 Correlation= -7.8817366160832125E-003
Approx. D= -3.7098639799501050E-004 Approx. Corr.=
-1.0375360285176364E-002

q relative to approx. equilib. value= 0.93767297328361821

Load statistics for gene model

Relative L= 0.93893992906505619 Relative B= 0.78599994996463829

Relative V x h = 0.29899263170114443

#### Dominance coefficient= 0.29999999999999999

Initial state (approximate equilibrium equation; no LD)
x = 3.0909728018113612E-002 y= 1.0194739375238855E-003 q=
3.1929201955637497E-002

**Sites Model**

Epistasis parameter= 0.0000000000000000

Iteration of haplotype frequencies

Population genetic statistics for sites model
Gen= 5000 x= 3.0942997049961146E-002 y= 1.0200919057382408E-003
q= 3.1963088955699388E-002

wx= 0.99668036911044311 wy= 0.99355251675462042 wz=
0.99980822146626591
wy*wz-wx**2= -9.7834603597357628E-006
wbar= 0.99960826982008710

D= -1.5471498517119486E-006 Correlation= -5.0002500099598648E-005
Approx. D= -1.4980342225597902E-006 Approx. Corr.=
-4.8464814109070043E-005

q relative to approx. equilib. value= 1.0010613168506051

Load statistics for sites model

Relative L= 0.97932544978278047 Relative B= 0.88999005053809899

Relative V x h = 0.30175500143930223

Epistasis parameter= 8.0000000000000002E-002

Iteration of haplotype frequencies

Population genetic statistics for sites model
Gen= 5000 x= 3.0807835228095402E-002 y= 9.3626179588876047E-004

q=   3.1744097023984161E-002

wx=  0.99666679755658250      wy=  0.99304002925299673      wz=
0.99980908601219387
wy*wz-wx**2=  -4.9426133073293688E-004
wbar=  0.99960913419299657

D=  -7.1425899979361829E-005  Correlation=  -2.3238205613700358E-003
Approx. D=  -8.2478395855911112E-005  Approx. Corr.=
-2.6683636882077191E-003

q relative to approx. equilib. value=  0.99420264459128915

Load statistics for sites model

Relative L=  0.97716451750749378      Relative B=  0.88275946857940935

Relative V x h =  0.30411761714838620

Epistasis parameter=  0.32000000000000001

Iteration of haplotype frequencies

Population genetic statistics for sites model
Gen=        5000  x=   3.0415515395561456E-002 y=   7.4892812805121074E-004
q=   3.1164443523612667E-002

wx=  0.99662684223419007      wy=  0.99150408108368115      wz=
0.99981157539685162
wy*wz-wx**2=  -1.9478053410102447E-003
wbar=  0.99961162307960660

D=  -2.2229441208523187E-004  Correlation=  -7.3623947642713055E-003
Approx. D=  -3.1381334555871828E-004  Approx. Corr.=
-1.0152575440806806E-002

q relative to approx. equilib. value=  0.97604830734299630

Load statistics for sites model

Relative L=  0.97094230097904777      Relative B=  0.86182214553707248

Relative V x h =  0.31226953659192380

**Gene Model**

Epistasis parameter=    0.0000000000000000

Iteration of haplotype frequencies

Population genetic statistics for gene model
Gen=         5000  x=    2.9863687301112475E-002 y=    9.6683474719411102E-004
q=    3.0830522048306587E-002
wx=   0.99657030636081578       wy=   0.99344891794211632       wz=
0.99981501686770913
wy*wz-wx**2=    1.1277112941410206E-004
wbar=   0.99961506386214172

D=    1.6313657422991484E-005  Correlation=    5.4597245313445534E-004
Approx. D=    1.5875060548338475E-005  Approx. Corr.=
5.1359431370717431E-004

q relative to approx. equilib. value=   0.96559012314628412

Load statistics for gene model

Relative L=   0.96234034464023355       Relative B=   0.83252231348384698

Relative V x h =   0.31936357368306495

Epistasis parameter=    8.0000000000000002E-002

Iteration of haplotype frequencies

Population genetic statistics for gene model
Gen=         5000  x=    2.9864227501481086E-002 y=    8.9126060264365845E-004
q=    3.0755488104124745E-002
wx=   0.99657013617502210       wy=   0.99292596355705609       wz=
0.99981503926628501
wy*wz-wx**2=   -4.0972507359016141E-004
wbar=   0.99961508625624029

D=   -5.4639445879300919E-005  Correlation=   -1.8329487068853719E-003
Approx. D=   -6.1307494120234124E-005  Approx. Corr.=
-1.9834368676523756E-003

q relative to approx. equilib. value=   0.96324011313707392

Load statistics for gene model

Relative L=   0.96228435939418200       Relative B=   0.83248655776872504

Relative V x h =   0.32049926086306757

Epistasis parameter=   0.32000000000000001

Iteration of haplotype frequencies

Population genetic statistics for gene model
Gen=         5000  x=    2.9865437686506579E-002 y=    7.2195853955942457E-004
q=    3.0587396226066002E-002
wx=   0.99656975496892197       wy=   0.99135705540675367       wz=
0.99981508946224662
wy*wz-wx**2=   -1.9775334782843323E-003
wbar=   0.99961513644216682

D=   -2.1363026833093324E-004  Correlation=   -7.2046288888273453E-003
Approx. D=   -2.9285515812595191E-004  Approx. Corr.=
-9.4745304117310244E-003

q relative to approx. equilib. value=   0.95797559452203651

Load statistics for gene model

Relative L=   0.96215889457758219       Relative B=   0.83240641217360056

Relative V x h =   0.32388521455873731

## Dominance coefficient=   0.34999999999999998

Initial state (approximate equilibrium equation; no LD)
x =   2.7108163369772110E-002  y=    7.7761673187756069E-004  q=
2.7885780101649670E-002

**Sites Model**

Epistasis parameter=    0.0000000000000000

Iteration of haplotype frequencies

Population genetic statistics for sites model
Gen=         5000  x=    2.7120433953631420E-002 y=    7.7690552848267683E-004
q=    2.7897339482114095E-002

wx=   0.99622102660517886       wy=   0.99263733458673253       wz=
0.99980471862362519
wy*wz-wx**2=   -1.2842848483129998E-005
wbar=   0.99960476767794926

D=   -1.3560216976481188E-006  Correlation=   -5.0002500123772657E-005

Approx. D=   -1.3184463516387736E-006  Approx. Corr.=
-4.8636506046328193E-005

q relative to approx. equilib. value=    1.0004145259850106

Load statistics for sites model

Relative L=   0.98808080512669794       Relative B=   0.90384823845619300

Relative V x h =   0.34258480998041874

Epistasis parameter=    8.0000000000000002E-002

Iteration of haplotype frequencies

Population genetic statistics for sites model
Gen=         5000  x=    2.7012296637610868E-002 y=    7.1363759513632091E-004
q=    2.7725934232747190E-002

wx=   0.99620684304295271       wy=   0.99205072788337212       wz=
0.99980551882331747
wy*wz-wx**2=   -5.7028143512161122E-004
wbar=   0.99960556771759668

D=   -5.5089833942298383E-005  Correlation=   -2.0436031946307314E-003
Approx. D=   -6.3038112353434243E-005  Approx. Corr.=
-2.3254291149701072E-003

q relative to approx. equilib. value=   0.99426783585326262

Load statistics for sites model

Relative L=   0.98608070600818065       Relative B=   0.89553879552910121

Relative V x h =   0.34502401111701775

Epistasis parameter=   0.32000000000000001

Iteration of haplotype frequencies

Population genetic statistics for sites model
Gen=         5000  x=    2.6697677707076602E-002 y=    5.7198540844888334E-004
q=    2.7269663115525485E-002

wx=   0.99616502959381625       wy=   0.99029205258093733       wz=
0.99980783111087623

wy*wz-wx**2= -2.2430169283640389E-003
wbar= 0.99960787954270025

D= -1.7164911798536742E-004 Correlation= -6.4709693503066535E-003
Approx. D= -2.4269022719780470E-004 Approx. Corr.=
-8.9526621146316671E-003

q relative to approx. equilib. value= 0.97790569301348906

Load statistics for sites model

Relative L= 0.98030114324848816 Relative B= 0.87140260802969516

Relative V x h = 0.35347859312146446

**Gene Model**

Epistasis parameter= 0.0000000000000000

Iteration of haplotype frequencies

Population genetic statistics for gene model
Gen= 5000 x= 2.6551751001246359E-002 y= 7.5237076045033512E-004
q= 2.7304121761696692E-002
wx= 0.99614617497323843 wy= 0.99256539116409392 wz=
0.99980887114766803
wy*wz-wx**2= 6.8481366202632898E-005
wbar= 0.99960891937149099

D= 6.8556952727734377E-006 Correlation= 2.5813459836626033E-004
Approx. D= 5.8095995411145859E-006 Approx. Corr.=
2.1431180939365224E-004

q relative to approx. equilib. value= 0.97914139974450387

Load statistics for gene model

Relative L= 0.97770157127191182 Relative B= 0.86028894411961554

Relative V x h = 0.35327693311024261

Epistasis parameter= 8.0000000000000002E-002

Iteration of haplotype frequencies

Population genetic statistics for gene model
Gen= 5000 x= 2.6552018884989339E-002 y= 6.9386557059144247E-004

q= 2.7245884455580783E-002
wx= 0.99614601166174843 wy= 0.99197143924105602 wz= 0.99980889024409125
wy*wz-wx**2= -5.2501272817406797E-004
wbar= 0.99960893846409538

D= -4.8472649175412136E-005 Correlation= -1.8289118272750845E-003
Approx. D= -5.3872637720463366E-005 Approx. Corr.= -1.9873215675148210E-003

q relative to approx. equilib. value= 0.97705297668789148

Load statistics for gene model

Relative L= 0.97765383976094200 Relative B= 0.86025680733500021

Relative V x h = 0.35430458675493781

Epistasis parameter= 0.32000000000000001

Iteration of haplotype frequencies

Population genetic statistics for gene model
Gen= 5000 x= 2.6552619823406523E-002 y= 5.6261562074018390E-004
q= 2.7115235444146707E-002
wx= 0.99614564530767746 wy= 0.99018955418527743 wz= 0.99980893309290042
wy*wz-wx**2= -2.3057849357321913E-003
wbar= 0.99960898130433562

D= -1.7262037245132505E-004 Correlation= -6.5436078864313475E-003
Approx. D= -2.3291934950519721E-004 Approx. Corr.= -8.5922216982402405E-003

q relative to approx. equilib. value= 0.97236782852428150

Load statistics for gene model

Relative L= 0.97754673916058843 Relative B= 0.86018469006483211

Relative V x h = 0.35736934730181286

#### Dominance coefficient= 0.40000000000000002

Initial state (approximate equilibrium equation; no LD)
x = 2.4078073746484154E-002 y= 6.0947506638794240E-004 q=

2.4687548812872096E-002

**Sites Model**

Epistasis parameter=    0.0000000000000000

Iteration of haplotype frequencies

Population genetic statistics for sites model
Gen=          5000  x=    2.4080693567726699E-002 y=    6.0834456946112087E-004
q=    2.4689038137187820E-002

wx=   0.99575310961862817       wy=   0.99170373154235369       wz=
0.99980248769490254
wy*wz-wx**2=   -1.6397462804684793E-005
wbar=   0.99960253719538850

D=   -1.2040346783932787E-006  Correlation=   -5.0002500125294633E-005
Approx. D=   -1.1745270833398974E-006  Approx. Corr.=
-4.8779943765700953E-005

q relative to approx. equilib. value=    1.0000603269416097

Load statistics for sites model

Relative L=   0.99365701152890784       Relative B=   0.90234512437582026

Relative V x h =   0.38672362220519274

Epistasis parameter=    8.0000000000000002E-002

Iteration of haplotype frequencies

Population genetic statistics for sites model
Gen=          5000  x=    2.3993131587196369E-002 y=    5.5921343065698677E-004
q=    2.4552345017853354E-002

wx=   0.99573849462656328       wy=   0.99104180160856858       wz=
0.99980322334326155
wy*wz-wx**2=   -6.4836196501438792E-004
wbar=   0.99960327269662386

D=   -4.3604215218720488E-005  Correlation=   -1.8206712199982692E-003
Approx. D=   -4.9440433803047806E-005  Approx. Corr.=
-2.0533384158384774E-003

q relative to approx. equilib. value=   0.99452340141001572

Load statistics for sites model

Relative L= 0.99181825844032612 Relative B= 0.89174833418892363

Relative V x h = 0.38920962894755967

Epistasis parameter= 0.32000000000000001

Iteration of haplotype frequencies

Population genetic statistics for sites model
Gen= 5000 x= 2.3737766752156837E-002 y= 4.4903303468940916E-004
q= 2.4186799786846246E-002

wx= 0.99569535165125056 wy= 0.98905688109137635 wz=
0.99980535607713639
wy*wz-wx**2= -2.5448661198019806E-003
wbar= 0.99960540500397121

D= -1.3596824923957709E-004 Correlation= -5.7609274658781181E-003
Approx. D= -1.9191746030475341E-004 Approx. Corr.=
-7.9706318007592668E-003

q relative to approx. equilib. value= 0.97971653525340030

Load statistics for sites model

Relative L= 0.98648749007183389 Relative B= 0.86087575072303357

Relative V x h = 0.39786171114456070

**Gene Model**

Epistasis parameter= 0.0000000000000000

Iteration of haplotype frequencies

Population genetic statistics for gene model
Gen= 5000 x= 2.3807620646302173E-002 y= 5.9791455498141180E-004
q= 2.4405535201283586E-002
wx= 0.99570773149213954 wy= 0.99165951833629196 wz=
0.99980475571838967
wy*wz-wx**2= 3.2015932809770220E-005
wbar= 0.99960480476529057

D= 2.2844065203191100E-006 Correlation= 9.5943537584510869E-005

Approx. D= 1.1729514214564947E-006 Approx. Corr.=
4.8714504067326705E-005

q relative to approx. equilib. value= 0.98857668642091079

Load statistics for gene model

Relative L= 0.98798808677337591 Relative B= 0.86936325709685547

Relative V x h = 0.39253594679974801

Epistasis parameter= 8.0000000000000002E-002

Iteration of haplotype frequencies

Population genetic statistics for gene model
Gen= 5000 x= 2.3807755753373017E-002 y= 5.5163264018477296E-004
q= 2.4359388393557790E-002
wx= 0.99570757701274926 wy= 0.99099287757399224 wz=
0.99980477184796179
wy*wz-wx**2= -6.3417105477958646E-004
wbar= 0.99960482089163694

D= -4.1747162723423242E-005 Correlation= -1.7565912493967576E-003
Approx. D= -4.6045411236910902E-005 Approx. Corr.=
-1.9123378274241887E-003

q relative to approx. equilib. value= 0.98670745233551893

Load statistics for gene model

Relative L= 0.98794777090750752 Relative B= 0.86933641161571129

Relative V x h = 0.39349642965914672

Epistasis parameter= 0.32000000000000001

Iteration of haplotype frequencies

Population genetic statistics for gene model
Gen= 5000 x= 2.3808059205534280E-002 y= 4.4767471813662917E-004
q= 2.4255733923670909E-002
wx= 0.99570723002898698 wy= 0.98899293589834647 wz=
0.99980480808133221
wy*wz-wx**2= -2.6329954623584007E-003
wbar= 0.99960485711776115

D= -1.4066591003928861E-004 Correlation= -5.9434475222103940E-003
Approx. D= -1.8770049921201313E-004 Approx. Corr.=
-7.7954948218987366E-003

q relative to approx. equilib. value= 0.98250879856585682

Load statistics for gene model

Relative L= 0.98785720559726953 Relative B= 0.86927610094164764

Relative V x h = 0.39636193457748520

#### Dominance coefficient= 0.45000000000000001

Initial state (approximate equilibrium equation; no LD)
x = 2.1622219167201234E-002 y= 4.8890166443104276E-004 q=
2.2111120831632275E-002

**Sites Model**

Epistasis parameter= 0.0000000000000000

Iteration of haplotype frequencies

Population genetic statistics for sites model
Gen= 5000 x= 2.1620388880926131E-002 y= 4.8768595343776761E-004
q= 2.2108074834363899E-002

wx= 0.99527891925165635 wy= 0.99075681117682202 wz=
0.99980102732649068
wy*wz-wx**2= -2.0449461440419370E-005
wbar= 0.99960107711903556

D= -1.0810194440678822E-006 Correlation= -5.0002500126004277E-005
Approx. D= -1.0573181343792020E-006 Approx. Corr.=
-4.8899612301731223E-005

q relative to approx. equilib. value= 0.99986224139013258

Load statistics for sites model

Relative L= 0.99730720241078485 Relative B= 0.86313951424319257

Relative V x h = 0.43287099948729374

Epistasis parameter= 8.0000000000000002E-002

Iteration of haplotype frequencies

Population genetic statistics for sites model
Gen= 5000 x= 2.1548514326842155E-002 y= 4.4858808255995907E-004
q= 2.1997102409402114E-002

wx= 0.99526399368341489 wy= 0.99001867442337632 wz=
0.99980170309489591
wy*wz-wx**2= -7.2806033841765760E-004
wbar= 0.99960175275229379

D= -3.5284431849764292E-005 Correlation= -1.6401271227979304E-003
Approx. D= -3.9659568429132080E-005 Approx. Corr.=
-1.8342043488899476E-003

q relative to approx. equilib. value= 0.99484339020629631

Load statistics for sites model

Relative L= 0.99561811926535093 Relative B= 0.84527527775288080

Relative V x h = 0.43538574952580922

Epistasis parameter= 0.32000000000000001

Iteration of haplotype frequencies

Population genetic statistics for sites model
Gen= 5000 x= 2.1338432706759815E-002 y= 3.6078735762798045E-004
q= 2.1699220064387795E-002

wx= 0.99521987905982123 wy= 0.98780492732466507 wz=
0.99980366795183051
wy*wz-wx**2= -2.8516181157536291E-003
wbar= 0.99960371721627650

D= -1.1006879377474868E-004 Correlation= -5.1849871344734804E-003
Approx. D= -1.5486711113966937E-004 Approx. Corr.=
-7.1624059464991214E-003

q relative to approx. equilib. value= 0.98137133027398538

Load statistics for sites model

Relative L= 0.99070695930865893 Relative B= 0.79308592947949652

Relative V x h = 0.44416925735870655

**Gene Model**

Epistasis parameter= 0.0000000000000000

Iteration of haplotype frequencies

Population genetic statistics for gene model
Gen= 5000 x= 2.1519649212389847E-002 y= 4.8424195263895774E-004
q= 2.2003891165028804E-002
wx= 0.99525819931816095 wy= 0.99073643754797225 wz=
0.99980196497951479
wy*wz-wx**2= 1.3537272388619570E-006
wbar= 0.99960201458453835

D= 7.0726236523627195E-008 Correlation= 3.2865779259830710E-006
Approx. D= -9.6641935176001870E-007 Approx. Corr.=
-4.4695659788056416E-005

q relative to approx. equilib. value= 0.99515041922026548

Load statistics for gene model

Relative L= 0.99496353865420339 Relative B= 0.83812471673668920

Relative V x h = 0.43526517220286803

Epistasis parameter= 8.0000000000000002E-002

Iteration of haplotype frequencies

Population genetic statistics for gene model
Gen= 5000 x= 2.1519718099589340E-002 y= 4.4690752270794656E-004
q= 2.1966625622297287E-002
wx= 0.99525805428288883 wy= 0.98999579519205960 wz=
0.99980197859598297
wy*wz-wx**2= -7.3883978023692176E-004
wbar= 0.99960202819828337

D= -3.5625118522220750E-005 Correlation= -1.6582091016649121E-003
Approx. D= -3.9095998545530080E-005 Approx. Corr.=
-1.8081399621013388E-003

q relative to approx. equilib. value= 0.99346504365675248

Load statistics for gene model

Relative L=   0.99492950429151139       Relative B=   0.83810611391540513

Relative V x h =   0.43618251020390658

Epistasis parameter=   0.32000000000000001

Iteration of haplotype frequencies

Population genetic statistics for gene model
Gen=         5000  x=    2.1519872986265454E-002 y=    3.6295689146030450E-004
q=    2.1882829877725760E-002
wx=   0.99525772815671176       wy=   0.98777385507783355       wz=
0.99980200921525308
wy*wz-wx**2=   -2.9596604985450137E-003
wbar=   0.99960205881143005

D=   -1.1590135199717952E-004  Correlation=   -5.4149460659730399E-003
Approx. D=   -1.5348473612684024E-004  Approx. Corr.=
-7.0984728690411851E-003

q relative to approx. equilib. value=   0.98967528803063110

Load statistics for gene model

Relative L=   0.99485297142515827       Relative B=   0.83806427746352707

Relative V x h =   0.43892039257901672

**Dominance coefficient=   0.50000000000000000**

Initial state (approximate equilibrium equation; no LD)
x =    1.9599999999999999E-002  y=    4.0000000000000002E-004  q=
2.0000000000000000E-002

**Sites Model**

Epistasis parameter=    0.0000000000000000

Iteration of haplotype frequencies

Population genetic statistics for sites model
Gen=         5000  x=    1.9596180935781617E-002 y=    3.9882026392964955E-004
q=    1.9995001199711267E-002

wx=   0.99480004998800287       wy=   0.98980004998800286       wz=
0.99980004998800287
wy*wz-wx**2=   -2.5000000000052758E-005

wbar= 0.99960009997600563

D= -9.7980904680688985E-007 Correlation= -5.0002500125915142E-005
Approx. D= -9.6039999999999994E-007 Approx. Corr.=
-4.8999999999999998E-005

q relative to approx. equilib. value= 0.99975005998556332

Load statistics for sites model

Relative L= 0.99975005998556354 Relative B= 1.0003988202639296

Relative V x h = 0.48027977048667292

Epistasis parameter= 2.0000000000000000E-002

Iteration of haplotype frequencies

Population genetic statistics for sites model
Gen= 5000 x= 1.9581163090181198E-002 y= 3.9039077171027658E-004
q= 1.9971553861891474E-002

wx= 0.99479625111153158 wy= 0.98959629015060868 wz=
0.99980020638322675
wy*wz-wx**2= -2.2100609690323125E-004
wbar= 0.99960025633995209

D= -8.4721919481542396E-006 Correlation= -4.3285780132148717E-004
Approx. D= -8.1498169717138101E-006 Approx. Corr.=
-4.1580698835274541E-004

q relative to approx. equilib. value= 0.99857769309457367

Load statistics for sites model

Relative L= 0.99935915011961796 Relative B= 1.0004138412372778

Relative V x h = 0.48089258572574217

Epistasis parameter= 4.0000000000000001E-002

Iteration of haplotype frequencies

Population genetic statistics for sites model
Gen= 5000 x= 1.9566191017037760E-002 y= 3.8229918133696970E-004
q= 1.9948490198374729E-002

wx= 0.99479245924210069 wy= 0.98939253570193686 wz=
0.99980036217834367
wy*wz-wx**2= -4.1702143360033972E-004
wbar= 0.99960041210391160

D= -1.5643079857685649E-005 Correlation= -8.0013511170889311E-004
Approx. D= -1.6272558139534881E-005 Approx. Corr.=
-8.3023255813953478E-004

q relative to approx. equilib. value= 0.99742450991873643

Load statistics for sites model

Relative L= 0.99896974022085616 Relative B= 1.0004288164261683

Relative V x h = 0.48151897845754932

Epistasis parameter= 8.0000000000000002E-002

Iteration of haplotype frequencies

Population genetic statistics for sites model
Gen= 5000 x= 1.9536383176144569E-002 y= 3.6705174295987431E-004
q= 1.9903434919104443E-002

wx= 0.99478489608217646 wy= 0.98898504290287370 wz=
0.99980067200941458
wy*wz-wx**2= -8.0907897167381559E-004
wbar= 0.99960072187301929

D= -2.9094978619151335E-005 Correlation= -1.4914927346406947E-003
Approx. D= -3.2437350993377485E-005 Approx. Corr.=
-1.6549668874172184E-003

q relative to approx. equilib. value= 0.99517174595522218

Load statistics for sites model

Relative L= 0.99819531745146572 Relative B= 1.0004586304633729

Relative V x h = 0.48281229354955313

Epistasis parameter= 0.16000000000000000

Iteration of haplotype frequencies

Population genetic statistics for sites model
Gen= 5000 x= 1.9477305719999002E-002 y= 3.3983202794141246E-004
q= 1.9817137747940413E-002

wx= 0.99476984933650159 wy= 0.98817012120212389 wz=
0.99980128489127584
wy*wz-wx**2= -1.5932962799148909E-003
wbar= 0.99960133463231038

D= -5.2886920579434116E-005 Correlation= -2.7227028579621883E-003
Approx. D= -6.4447894736842113E-005 Approx. Corr.=
-3.2881578947368422E-003

q relative to approx. equilib. value= 0.99085688739702060

Load statistics for sites model

Relative L= 0.99666341922384771 Relative B= 1.0005177201765438

Relative V x h = 0.48555960478414456

Epistasis parameter= 0.32000000000000001

Iteration of haplotype frequencies

Population genetic statistics for sites model
Gen= 5000 x= 1.9361249129468742E-002 y= 2.9562764544160996E-004
q= 1.9656876774910351E-002

wx= 0.99474005622233852 wy= 0.98654052922657121 wz=
0.99980248522378545
wy*wz-wx**2= -3.1621065585067942E-003
wbar= 0.99960253472476557

D= -9.0765159102400989E-005 Correlation= -4.7100612128879241E-003
Approx. D= -1.2722181818181817E-004 Approx. Corr.=
-6.4909090909090906E-003

q relative to approx. equilib. value= 0.98284383874551751

Load statistics for sites model

Relative L= 0.99366318808612430 Relative B= 1.0006338007725246

Relative V x h = 0.49168533808832421

Epistasis parameter=   0.64000000000000001

Iteration of haplotype frequencies

Population genetic statistics for sites model
Gen=         5000  x=    1.9137115382469785E-002 y=    2.3382901001668490E-004
q=    1.9370944392486468E-002

wx=   0.99468156825913112       wy=   0.98328231651196318       wz=
0.99980479405041101
wy*wz-wx**2=   -6.3010482807901447E-003
wbar=   0.99960484308964881

D=   -1.4140447664011749E-004  Correlation=   -7.4440213025521104E-003
Approx. D=   -2.4800202531645568E-004  Approx. Corr.=
-1.2653164556962023E-002

q relative to approx. equilib. value=   0.96854721962432333

Load statistics for sites model

Relative L=   0.98789227587786288       Relative B=    1.0008579806937477

Relative V x h =   0.50636916585414682

**Gene Model**

Epistasis parameter=    0.0000000000000000

Iteration of haplotype frequencies

Population genetic statistics for gene model
Gen=         5000  x=    1.9596180935781617E-002 y=    3.9882026392964955E-004
q=    1.9995001199711267E-002
wx=   0.99480004998800287       wy=   0.98980004998800286       wz=
0.99980004998800287
wy*wz-wx**2=   -2.5000000000052758E-005
wbar=   0.99960009997600563

D=   -9.7980904680688985E-007  Correlation=   -5.0002500125915142E-005
Approx. D=   -1.9207999999999999E-006  Approx. Corr.=
-9.7999999999999997E-005

q relative to approx. equilib. value=   0.99975005998556332

Load statistics for gene model

Relative L=   0.99975005998556332       Relative B=    1.0003988202639296

Relative V x h =   0.48027977048667292

Epistasis parameter=    2.0000000000000000E-002

Iteration of haplotype frequencies

Population genetic statistics for gene model
Gen=         5000  x=    1.9596190242236463E-002 y=    3.9069095738635120E-004
q=    1.9986881199622813E-002
wx=   0.99480001398071660       wy=   0.98959613381176381       wz=
0.99980005304981234
wy*wz-wx**2=   -2.2880073314324179E-004
wbar=   0.99960010303720293

D=   -8.7844627014860288E-006  Correlation=   -4.4847504566756260E-004
Approx. D=   -9.7576640000000013E-006  Approx. Corr.=
-4.9784000000000004E-004

q relative to approx. equilib. value=   0.99934405998114062

Load statistics for gene model

Relative L=   0.99974240699285377       Relative B=    1.0003988110187072

Relative V x h =   0.48050221315182173

Epistasis parameter=    4.0000000000000001E-002

Iteration of haplotype frequencies

Population genetic statistics for gene model
Gen=         5000  x=    1.9596199176745710E-002 y=    3.8288643603360222E-004
q=    1.9979085612779311E-002
wx=   0.99479997941201059       wy=   0.98939221750962714       wz=
0.99980005598929778
wy*wz-wx**2=   -4.3260457663596430E-004
wbar=   0.99960010597610049

D=   -1.6277425889162783E-005  Correlation=   -8.3133253014408537E-004
Approx. D=   -1.7594527999999999E-005  Approx. Corr.=
-8.9767999999999998E-004

q relative to approx. equilib. value=   0.99895428063896552

Load statistics for gene model

Relative L=   0.99973505974875809       Relative B=    1.0003988021429839

Relative V x h =   0.48072447689293529

Epistasis parameter=    8.0000000000000002E-002

Iteration of haplotype frequencies

Population genetic statistics for gene model
Gen=         5000  x=    1.9596216015694161E-002 y=    3.6817688429077012E-004
q=    1.9964392899984931E-002
wx=   0.99479991425873904       wy=   0.98898438455668014       wz=
0.99980006152949275
wy*wz-wx**2=   -8.4022087771817855E-004
wbar=   0.99960011151518746

D=   -3.0400099574199452E-005  Correlation=   -1.5537353363911238E-003
Approx. D=   -3.3268256000000008E-005  Approx. Corr.=
-1.6973600000000002E-003

q relative to approx. equilib. value=   0.99821964499924654

Load statistics for gene model

Relative L=   0.99972121203143571       Relative B=    1.0003987854148351

Relative V x h =   0.48116850884381046

Epistasis parameter=   0.16000000000000000

Iteration of haplotype frequencies

Population genetic statistics for gene model
Gen=         5000  x=    1.9596246087947918E-002 y=    3.4190647737139003E-004
q=    1.9938152565319306E-002
wx=   0.99479979789880113       wy=   0.98816871743024226       wz=
0.99980007142398297
wy*wz-wx**2=   -1.6554836337937529E-003
wbar=   0.99960012140769883

D=   -5.5623450346559827E-005  Correlation=   -2.8465546558135517E-003
Approx. D=   -6.4615712000000005E-005  Approx. Corr.=
-3.2967200000000004E-003

q relative to approx. equilib. value=   0.99690762826596524

Load statistics for gene model

Relative L= 0.99969648075269346 Relative B= 1.0003987555404610

Relative V x h = 0.48205483810992067

Epistasis parameter= 0.32000000000000001

Iteration of haplotype frequencies

Population genetic statistics for gene model
Gen= 5000 x= 1.9596294962757929E-002 y= 2.9920790365267191E-004
q= 1.9895502866410602E-002
wx= 0.99479960877339835 wy= 0.98653737936216335 wz=
0.99980008750604421
wy*wz-wx**2= -3.2861034014319834E-003
wbar= 0.99960013748654375

D= -9.6623130654678591E-005 Correlation= -4.9551157315172491E-003
Approx. D= -1.2731062400000000E-004 Approx. Corr.=
-6.4954400000000008E-003

q relative to approx. equilib. value= 0.99477514332053008

Load statistics for gene model

Relative L= 0.99965628364033554 Relative B= 1.0003987069872760

Relative V x h = 0.48382207442840508

Epistasis parameter= 0.64000000000000001

Iteration of haplotype frequencies

Population genetic statistics for gene model
Gen= 5000 x= 1.9596363402881432E-002 y= 2.3941083714168636E-004
q= 1.9835774240023116E-002
wx= 0.99479934391356317 wy= 0.98327469330246364 wz=
0.99980011002824209
wy*wz-wx**2= -6.5475880990665258E-003
wbar= 0.99960016000423746

D= -1.5404710255947918E-004 Correlation= -7.9232896008860961E-003
Approx. D= -2.5270044800000002E-004 Approx. Corr.=
-1.2892880000000001E-002

q relative to approx. equilib. value= 0.99178871200115581

Load statistics for gene model

Relative L= 0.99959998940624073 Relative B= 1.0003986389975743

Relative V x h = 0.48734244339747945

#### Results with lower mutation rate

**Mutation rate= 1.0000000000000000E-005**
**Selection coefficient= 1.0000000000000000E-002**
**Maximum number of generations= 5000**

#### Dominance coefficient=0

##### Sites model

Initial state (approximate equilibrium equation; no LD)
x = 3.0622776601683790E-002 y= 9.9999999999999980E-004 q=
3.1622776601683791E-002

Epistasis parameter= 0.0000000000000000

Iteration of haplotype frequencies

Population genetic statistics for sites model
Gen= 5000 x= 3.0622841358565613E-002 y= 9.9984855941616358E-004
q= 3.1622689917981776E-002

wx= 0.99968377310084422 wy= 0.99936754620168855 wz=
1.0000000000000000
wy*wz-wx**2= -9.9999451608212553E-008
wbar= 0.99998000010965016

D= -1.4595823266336239E-007 Correlation= -4.7663417852425698E-006
Approx. D= -1.4827199689142735E-007 Approx. Corr.=
-4.8418861169915802E-006

q relative to approx. equilib. value= 9.9999725882065613E-002

Load statistics for sites model

Relative L= 0.49999725882441315 Relative B= 0.98367210499679414

Relative V x h =   0.49949711835996863

Epistasis parameter=   8.0000000000000002E-002

Iteration of haplotype frequencies

Population genetic statistics for sites model
Gen=        5000  x=   3.0619349122048913E-002 y=   9.2823480896803937E-004
q=   3.1547583931016955E-002

wx=   0.99968378157517035       wy=   0.99931857219501552       wz=
1.0000000000000000
wy*wz-wx**2=   -4.9090949417407437E-005
wbar=   0.99998000267044418

D=   -6.7015242916516893E-005  Correlation=   -2.1934574026859889E-003
Approx. D=   -7.5020355743730586E-005  Approx. Corr.=
-2.4498221281347033E-003

q relative to approx. equilib. value=   9.9762219897341853E-002

Load statistics for sites model

Relative L=   0.49993324639134518       Relative B=   0.98360543258029642

Relative V x h =   0.50192888817563663

Epistasis parameter=   0.16000000000000000

Iteration of haplotype frequencies

Population genetic statistics for sites model
Gen=        5000  x=   3.0616698155335121E-002 y=   8.6550161614969048E-004
q=   3.1482199771484809E-002

wx=   0.99968379320746126       wy=   0.99926961299301109       wz=
1.0000000000000000
wy*wz-wx**2=   -9.8073408647092464E-005
wbar=   0.99998000543352072

D=   -1.2562728630198874E-004  Correlation=   -4.1201336234332258E-003
Approx. D=   -1.5004071148746117E-004  Approx. Corr.=
-4.8996442562694066E-003

q relative to approx. equilib. value=   9.9555457030324465E-002

Load statistics for sites model

Relative L= 0.49986418694718138 Relative B= 0.98354767467922855

Relative V x h = 0.50435641336476567

**Gene model**

Initial state (approximate equilibrium equation; no LD)
x = 3.0622776601683790E-002 y= 9.9999999999999980E-004 q=
3.1622776601683791E-002

Epistasis parameter= 0.0000000000000000

Iteration of haplotype frequencies

Population genetic statistics for gene model
Gen= 5000 x= 2.1949879681490113E-002 y= 6.8324656502870306E-004
q= 2.2633126246518816E-002
wx= 0.99955075194269050 wy= 0.99932783604687780 wz=
1.0000000000000000
wy*wz-wx**2= 2.2613033767981960E-004
wbar= 0.99997981878538755

D= 1.7098816133784278E-004 Correlation= 7.7297221136502876E-003
Approx. D= 2.3443861169915808E-004 Approx. Corr.=
7.6556941504209476E-003

q relative to approx. equilib. value= 0.71572229509137064

Load statistics for gene model

Relative L= 0.50452640080896138 Relative B= 0.69498760478879407

Relative V x h = 0.51498951110819879

Epistasis parameter= 8.0000000000000002E-002

Iteration of haplotype frequencies

Population genetic statistics for gene model
Gen= 5000 x= 2.1950146965920531E-002 y= 6.3152598659660429E-004
q= 2.2581672952517135E-002
wx= 0.99955076458680581 wy= 0.99927517146566225 wz=
1.0000000000000000
wy*wz-wx**2= 1.7344047959422237E-004
wbar= 0.99997982060653257

D= 1.2159403326215956E-004 Correlation= 5.5090367935480104E-003
Approx. D= 1.7817334489136013E-004 Approx. Corr.=
5.8183275543199206E-003

q relative to approx. equilib. value= 0.71409519906973473

Load statistics for gene model

Relative L= 0.50448090027837489 Relative B= 0.69497973599675966

Relative V x h = 0.51763580179508961

Epistasis parameter= 0.32000000000000001

Iteration of haplotype frequencies

Population genetic statistics for gene model
Gen= 5000 x= 2.1951858617975724E-002 y= 5.1355541614141577E-004
q= 2.2465414034117139E-002
wx= 0.99955079271369551 wy= 0.99911714518916428 wz=
1.0000000000000000
wy*wz-wx**2= 1.5357974587182355E-005
wbar= 0.99997982465836321

D= 8.8605884171061300E-006 Correlation= 4.0347441874885348E-004
Approx. D= 9.3775444679663317E-006 Approx. Corr.=
3.0622776601683818E-004

q relative to approx. equilib. value= 0.71041876926521819

Load statistics for gene model

Relative L= 0.50437968649030229 Relative B= 0.69497440080199202

Relative V x h = 0.52552265371076834

#### Dominance coefficient = 0.1

##### Sites model

Initial state (approximate equilibrium equation; no LD)
x = 9.2148613267964696E-003 y= 8.6515612975725985E-005 q=
9.3013769397721959E-003

Epistasis parameter=    0.0000000000000000

Iteration of haplotype frequencies

Population genetic statistics for sites model
Gen=         5000  x=    9.2202384530188884E-003 y=    8.6570594861339704E-005
q=    9.3068090478802285E-003

wx=   0.99890693190973201       wy=   0.99783247743751768       wz=
0.99998138638194645
wy*wz-wx**2=   -1.1544524127771183E-006
wbar=   0.99996138689678471

D=   -4.6099792365417169E-008  Correlation=   -4.9998731695966150E-006
Approx. D=   -4.6419820946084792E-008  Approx. Corr.=
-5.0374953349648026E-006

q relative to approx. equilib. value=    1.0005840111784745

Load statistics for sites model

Relative L=   0.96532758264950491       Relative B=   0.91660857314381849

Relative V x h =   0.13279635700260295

Epistasis parameter=    8.0000000000000002E-002

Iteration of haplotype frequencies

Population genetic statistics for sites model
Gen=         5000  x=    9.2085811019025510E-003 y=    7.9961047660841340E-005
q=    9.2885421495633932E-003

wx=   0.99890557083957221       wy=   0.99765943074016561       wz=
0.99998141012188724
wy*wz-wx**2=   -1.7145508138149967E-004
wbar=   0.99996141054736609

D=   -6.3159676033739537E-006  Correlation=   -6.8634923154221449E-004
Approx. D=   -6.8432928044585361E-006  Approx. Corr.=
-7.4263654782937406E-004

q relative to approx. equilib. value=   0.99862011933373840

Load statistics for sites model

Relative L=   0.96473631337225862       Relative B=   0.91528593011186110

Relative V x h = 0.13299885724183980

Epistasis parameter= 0.32000000000000001

Iteration of haplotype frequencies

Population genetic statistics for sites model
Gen= 5000 x= 9.1739362815391867E-003 y= 6.5004307574550020E-005
q= 9.2389405891137363E-003

wx= 0.99890151045814135 wy= 0.99714048276807488 wz=
0.99998148051576685
wy*wz-wx**2= -6.8221135493029550E-004
wbar= 0.99996148068536428

D= -2.0353715634623473E-005 Correlation= -2.2235791820393227E-003
Approx. D= -2.7258068184618205E-005 Approx. Corr.=
-2.9580551695718707E-003

q relative to approx. equilib. value= 0.99328740776094293

Load statistics for sites model

Relative L= 0.96298284979804261 Relative B= 0.91135917811778799

Relative V x h = 0.13373065299182088

**Gene model**

Initial state (approximate equilibrium equation; no LD)
x = 9.2148613267964696E-003 y= 8.6515612975725985E-005 q=
9.3013769397721959E-003

Epistasis parameter= 0.0000000000000000

Iteration of haplotype frequencies

Population genetic statistics for gene model
Gen= 5000 x= 8.6952504238411547E-003 y= 7.9373631859687308E-005
q= 8.7746240557008415E-003
wx= 0.99884237424047628 wy= 0.99777249473325358 wz=
0.99998245074953973
wy*wz-wx**2= 6.8895995489426376E-005
wbar= 0.99996244883313090

D= 2.3796045408031041E-006 Correlation= 2.7359229273239021E-004

Approx. D= 2.7465133397869526E-006 Approx. Corr.=
2.9805259595173667E-004

q relative to approx. equilib. value= 0.94336828971859132

Load statistics for gene model

Relative L= 0.93877903791338946 Relative B= 0.85717235477952525

Relative V x h = 0.15985467114072477

Epistasis parameter= 8.0000000000000002E-002

Iteration of haplotype frequencies

Population genetic statistics for gene model
Gen= 5000 x= 8.6952644277566212E-003 y= 7.3403079002350616E-005
q= 8.7686675067589724E-003
wx= 0.99884236949249205 wy= 0.99759440998632287 wz=
0.99998245091815063
wy*wz-wx**2= -1.0917597300652560E-004
wbar= 0.99996244900699283

D= -3.4864508417394025E-006 Correlation= -4.0112055910762081E-004
Approx. D= -3.9906750118138494E-006 Approx. Corr.=
-4.3306945924504760E-004

q relative to approx. equilib. value= 0.94272789540058455

Load statistics for gene model

Relative L= 0.93877469167697558 Relative B= 0.85716904766964741

Relative V x h = 0.16003909432932878

Epistasis parameter= 0.32000000000000001

Iteration of haplotype frequencies

Population genetic statistics for gene model
Gen= 5000 x= 8.6952965132143043E-003 y= 5.9889042630546041E-005
q= 8.7551855558448497E-003
wx= 0.99884235872782601 wy= 0.99706015442025320 wz=
0.99998245129756791
wy*wz-wx**2= -6.4340028067066957E-004
wbar= 0.99996244939545398

D= -1.6764231486729800E-005 Correlation= -1.9316897077201009E-003
Approx. D= -2.4202240066616254E-005 Approx. Corr.=
-2.6264356248354003E-003

q relative to approx. equilib. value= 0.94127843786312315

Load statistics for gene model

Relative L= 0.93876498070851888 Relative B= 0.85716162427868592

Relative V x h = 0.16059176311252452

#### Dominance coefficient = 0.2

##### Sites model

Initial state (approximate equilibrium equation; no LD)
x = 4.9025452748942709E-003 y= 2.4273543670968582E-005 q=
4.9268188185652394E-003

Epistasis parameter= 0.0000000000000000

Iteration of haplotype frequencies

Population genetic statistics for sites model
Gen= 5000 x= 4.9027977163079439E-003 y= 2.4251298000998524E-005
q= 4.9270490143089422E-003

wx= 0.99795072950985708 wy= 0.99592116721577140 wz=
0.99998029180394288
wy*wz-wx**2= -4.1191231054948929E-006
wbar= 0.99996029229814187

D= -2.4513988404299691E-008 Correlation= -5.0000249639738452E-006
Approx. D= -2.4629351608981303E-008 Approx. Corr.=
-5.0237887113673754E-006

q relative to approx. equilib. value= 1.0000467229975731

Load statistics for sites model

Relative L= 0.99269254645860938 Relative B= 0.97651275746374144

Relative V x h = 0.20531643088429652

Epistasis parameter=   8.0000000000000002E-002

Iteration of haplotype frequencies

Population genetic statistics for sites model
Gen=        5000  x=   4.8990865902532139E-003 y=   2.2421979263572746E-005
q=   4.9215085695167868E-003

wx=  0.99794919568149332      wy=  0.99559495633191453      wz=
0.99998030679068761
wy*wz-wx**2=  -3.2724728929034441E-004
wbar=  0.99996030728405261

D=  -1.7992673362540734E-006  Correlation=  -3.6740081143174177E-004
Approx. D=  -1.9307083387562750E-006  Approx. Corr.=
-3.9381754384673444E-004

q relative to approx. equilib. value=  0.99892217488728374

Load statistics for sites model

Relative L=  0.99231789863005526      Relative B=  0.97551955393181089

Relative V x h =  0.20556125104450090

Epistasis parameter=  0.32000000000000001

Iteration of haplotype frequencies

Population genetic statistics for sites model
Gen=        5000  x=   4.8880194013755651E-003 y=   1.8274508421387234E-005
q=   4.9062939097969521E-003

wx=  0.99794461022194469      wy=  0.99461637907260403      wz=
0.99998035143298603
wy*wz-wx**2=  -1.2966087850025954E-003
wbar=  0.99996035192395238

D=  -5.7972115079232486E-006  Correlation=  -1.1874124696648460E-003
Approx. D=  -7.7047162208200271E-006  Approx. Corr.=
-1.5715747206406349E-003

q relative to approx. equilib. value=  0.99583404433486666

Load statistics for sites model

Relative L= 0.99120190098669270 Relative B= 0.97255892498676511

Relative V x h = 0.20643857396423415

**Gene model**

Initial state (approximate equilibrium equation; no LD)
x = 4.9025452748942709E-003 y= 2.4273543670968582E-005 q=
4.9268188185652394E-003

Epistasis parameter= 0.0000000000000000

Iteration of haplotype frequencies

Population genetic statistics for gene model
Gen= 5000 x= 4.8348413961999126E-003 y= 2.3749094236641232E-005
q= 4.8585904904365537E-003
wx= 0.99792233379937489 wy= 0.99589325350369229 wz=
0.99998056563802307
wy*wz-wx**2= 2.4914658122376920E-005
wbar= 0.99996056611905337

D= 1.4319268288025960E-007 Correlation= 2.9615954352679078E-005
Approx. D= 1.4922696993392845E-007 Approx. Corr.=
3.0438672478582405E-005

q relative to approx. equilib. value= 0.98615164660174071

Load statistics for gene model

Relative L= 0.98584702288254256 Relative B= 0.95834062418748955

Relative V x h = 0.21220679289773736

Epistasis parameter= 8.0000000000000002E-002

Iteration of haplotype frequencies

Population genetic statistics for gene model
Gen= 5000 x= 4.8348420935517798E-003 y= 2.1974275682819245E-005
q= 4.8568163692345994E-003
wx= 0.99792233049172774 wy= 0.99556474443628329 wz=
0.99998056570273963
wy*wz-wx**2= -3.0358135894315019E-004
wbar= 0.99996056618377693

D= -1.6143895616455711E-006 Correlation= -3.3401893320953834E-004
Approx. D= -1.7875043189326979E-006 Approx. Corr.=
-3.6460740670492790E-004

q relative to approx. equilib. value= 0.98579155193065826

Load statistics for gene model

Relative L= 0.98584540479629168 Relative B= 0.95833912261242626

Relative V x h = 0.21230219236569164

Epistasis parameter= 0.32000000000000001

Iteration of haplotype frequencies

Population genetic statistics for gene model
Gen= 5000 x= 4.8348436750560841E-003 y= 1.7949962944696522E-005
q= 4.8527936380007809E-003
wx= 0.99792232299179795 wy= 0.99457921703634911 wz=
0.99998056584948025
wy*wz-wx**2= -1.2890744912048557E-003
wbar= 0.99996056633052921

D= -5.5996431483238893E-006 Correlation= -1.1595278911825436E-003
Approx. D= -7.5976981855325785E-006 Approx. Corr.=
-1.5497456442554590E-003

q relative to approx. equilib. value= 0.98497505524548279

Load statistics for gene model

Relative L= 0.98584173598916058 Relative B= 0.95833571787690375

Relative V x h = 0.21258811151634013

#### Dominance coefficient = 0.3

##### Sites model

Initial state (approximate equilibrium equation; no LD)
x = 3.3075871884426038E-003 y= 1.1013107927064852E-005 q=
3.3186002963696685E-003

Epistasis parameter= 0.0000000000000000

Iteration of haplotype frequencies

Population genetic statistics for sites model
Gen= 5000 x= 3.3075701058445143E-003 y= 1.0996345441068336E-005
q= 3.3185664512855825E-003

wx= 0.99696681433548717 wy= 0.99395354006968195 wz=
0.99998008860129228
wy*wz-wx**2= -9.0798218010945675E-006
wbar= 0.99996008909951817

D= -1.6537850529990683E-008 Correlation= -5.0000250003572315E-006
Approx. D= -1.6592453669061031E-008 Approx. Corr.=
-5.0164826272875015E-006

q relative to approx. equilib. value= 0.99998980139785953

Load statistics for sites model

Relative L= 0.99777251204399431 Relative B= 0.98814244085779468

Relative V x h = 0.30021054779243145

Epistasis parameter= 8.0000000000000002E-002

Iteration of haplotype frequencies

Population genetic statistics for sites model
Gen= 5000 x= 3.3058426893805156E-003 y= 1.0171513651556371E-005
q= 3.3160142030320720E-003

wx= 0.99696524247510454 wy= 0.99346986186525010 wz=
0.99998009903245522
wy*wz-wx**2= -4.8960384967156045E-004
wbar= 0.99996009953046960

D= -8.2443654315401960E-007 Correlation= -2.4944992132683080E-004
Approx. D= -8.7765450831498966E-007 Approx. Corr.=
-2.6534584224467212E-004

q relative to approx. equilib. value= 0.99922072768437176

Load statistics for sites model

Relative L= 0.99751173825658868 Relative B= 0.98723387267938389

Relative V x h = 0.30046142621225308

Epistasis parameter= 0.32000000000000001

Iteration of haplotype frequencies

Population genetic statistics for sites model
Gen= 5000 x= 3.3006819170353836E-003 y= 8.2999622303702387E-006
q= 3.3089818792657539E-003

wx= 0.99696053834408371 wy= 0.99201885001487111 wz=
0.99998013017279685
wy*wz-wx**2= -1.9311762435861857E-003
wbar= 0.99996013067018041

D= -2.6493988469389503E-006 Correlation= -8.0332708435337742E-004
Approx. D= -3.5050253760702280E-006 Approx. Corr.=
-1.0596925119064175E-003

q relative to approx. equilib. value= 0.99710166448353643

Load statistics for sites model

Relative L= 0.99673324548539333 Relative B= 0.98452005085248062

Relative V x h = 0.30136185723416770

#### Gene model

Initial state (approximate equilibrium equation; no LD)
x = 3.3075871884426038E-003 y= 1.1013107927064852E-005 q=
3.3186002963696685E-003

Epistasis parameter= 0.0000000000000000

Iteration of haplotype frequencies

Population genetic statistics for gene model
Gen= 5000 x= 3.2932600657748492E-003 y= 1.0924948836597375E-005
q= 3.3041850146114464E-003
wx= 0.99695376325969298 wy= 0.99394056836953215 wz=
0.99998017488991231
wy*wz-wx**2= 4.0573106796504277E-006
wbar= 0.99996017538639126

D= 7.3102258144400190E-009 Correlation= 2.2197486507510787E-006
Approx. D= 7.4867418190157522E-009 Approx. Corr.=

2.2635055079352050E-006

q relative to approx. equilib. value=   0.99565621633494350

Load statistics for gene model

Relative L=   0.99561534021730091       Relative B=   0.98061890532154361

Relative V x h =   0.30237635885229991

Epistasis parameter=    8.0000000000000002E-002

Iteration of haplotype frequencies

Population genetic statistics for gene model
Gen=         5000  x=    3.2932601493666424E-003 y=    1.0110757822184974E-005
q=    3.3033709071888274E-003
wx=   0.99695376089590548       wy=   0.99345582614803785       wz=
0.99998017492139302
wy*wz-wx**2=   -4.8067055629830158E-004
wbar=   0.99996017541787141

D=   -8.0150152827630078E-007  Correlation=   -2.4343556411939124E-004
Approx. D=   -8.7448604502025006E-007  Approx. Corr.=
-2.6438790429346377E-004

q relative to approx. equilib. value=   0.99541089983102182

Load statistics for gene model

Relative L=   0.99561455321317482       Relative B=   0.98061814962583027

Relative V x h =   0.30246067465798532

Epistasis parameter=   0.32000000000000001

Iteration of haplotype frequencies

Population genetic statistics for gene model
Gen=         5000  x=    3.2932603390451679E-003 y=    8.2632751582176426E-006
q=    3.3015236142033854E-003
wx=   0.99695375553223053       wy=   0.99200159942901911       wz=
0.99998017499282643
wy*wz-wx**2=   -1.9348576796243133E-003
wbar=   0.99996017548930338

D= -2.6367830169254751E-006 Correlation= -8.0130193321415713E-004
Approx. D= -3.5204044055380472E-006 Approx. Corr.=
-1.0643421336976606E-003

q relative to approx. equilib. value= 0.99485425159969887

Load statistics for gene model

Relative L= 0.99561276741418692 Relative B= 0.98061643486852068

Relative V x h = 0.30271343782567450

#### Dominance coefficient = 0.4

##### Sites model

Initial state (approximate equilibrium equation; no LD)
x = 2.4906406200928270E-003 y= 6.2343847900086618E-006 q=
2.4968750048828359E-003

Epistasis parameter= 0.0000000000000000

Iteration of haplotype frequencies

Population genetic statistics for sites model
Gen= 5000 x= 2.4905988478638240E-003 y= 6.2216596525145639E-006
q= 2.4968205075163387E-003

wx= 0.99597503179492486 wy= 0.99197003815390983 wz=
0.99998002543593989
wy*wz-wx**2= -1.6039974064518958E-005
wbar= 0.99996002593542910

D= -1.2452994239530585E-008 Correlation= -5.0000250002099071E-006
Approx. D= -1.2484336157104198E-008 Approx. Corr.=
-5.0125000196290465E-006

q relative to approx. equilib. value= 0.99997817377065712

Load statistics for sites model

Relative L= 0.99935161427121055 Relative B= 0.99001289829532024

Relative V x h = 0.39862097400729596

Epistasis parameter= 8.0000000000000002E-002

Iteration of haplotype frequencies

Population genetic statistics for sites model
Gen= 5000 x= 2.4896121102458606E-003 y= 5.7563830642388337E-006
q= 2.4953684933100993E-003

wx= 0.99597344651616726 wy= 0.99132766002432671 wz=
0.99998003336796837
wy*wz-wx**2= -6.5523961557645283E-004
wbar= 0.99996003386729904

D= -4.7048085316612337E-007 Correlation= -1.8901329181534446E-004
Approx. D= -4.9730760020515596E-007 Approx. Corr.=
-1.9967055712221584E-004

q relative to approx. equilib. value= 0.99939664117355076

Load statistics for sites model

Relative L= 0.99915331752535430 Relative B= 0.98882784409968194

Relative V x h = 0.39887326526907835

Epistasis parameter= 0.32000000000000001

Iteration of haplotype frequencies

Population genetic statistics for sites model
Gen= 5000 x= 2.4866612372073794E-003 y= 4.7002356076306571E-006
q= 2.4913614728150099E-003

wx= 0.99596869947544908 wy= 0.98940053683427065 wz=
0.99998005707561433
wy*wz-wx**2= -2.5728450406400372E-003
wbar= 0.99996005757447082

D= -1.5066463805966934E-006 Correlation= -6.0625861512118222E-004
Approx. D= -1.9868411777720502E-006 Approx. Corr.=
-7.9772294796107515E-004

q relative to approx. equilib. value= 0.99779182696088353

Load statistics for sites model

Relative L= 0.99856063823104779 Relative B= 0.98528423249264008

Relative V x h =   0.39977995589946996

#### Gene model

Initial state (approximate equilibrium equation; no LD)
x =   2.4906406200928270E-003  y=    6.2343847900086618E-006  q=
2.4968750048828359E-003

Epistasis parameter=    0.0000000000000000

Iteration of haplotype frequencies

Population genetic statistics for gene model
Gen=         5000  x=    2.4875166084561157E-003 y=    6.2100875614190697E-006
q=    2.4937266960175348E-003
wx=   0.99597008148973531       wy=   0.99196510024643092       wz=
0.99998005018643188
wy*wz-wx**2=   -1.1092495055042839E-005
wbar=   0.99996005068542615

D=   -8.5852730116048326E-009  Correlation=   -3.4513549028150712E-006
Approx. D=   -8.6178440690873603E-009  Approx. Corr.=
-3.4600913514235425E-006

q relative to approx. equilib. value=   0.99873910033175695

Load statistics for gene model

Relative L=   0.99873286434778652       Relative B=   0.98631180023520826

Relative V x h =   0.39924143927867412

Epistasis parameter=    8.0000000000000002E-002

Iteration of haplotype frequencies

Population genetic statistics for gene model
Gen=         5000  x=    2.4875166241138395E-003 y=    5.7479575626106352E-006
q=    2.4932645816764503E-003
wx=   0.99597007967542839       wy=   0.99132231425511341       wz=
0.99998005020465375
wy*wz-wx**2=   -6.5386203085737726E-004
wbar=   0.99996005070364757

D=   -4.6841071163136738E-007  Correlation=   -1.8834002055416346E-004
Approx. D=   -5.0829717458544057E-007  Approx. Corr.=
-2.0408290561265164E-004

q relative to approx. equilib. value=   0.99855402324933162

Load statistics for gene model

Relative L=   0.99873240880968295       Relative B=   0.98631136030929056

Relative V x h =   0.39932309543719191

Epistasis parameter=   0.32000000000000001

Iteration of haplotype frequencies

Population genetic statistics for gene model
Gen=         5000  x=    2.4875166596565845E-003 y=    4.6989304540692557E-006
q=    2.4922155901106538E-003
wx=   0.99597007555698291       wy=   0.98939395626107118       wz=
0.99998005024601710
wy*wz-wx**2=   -2.5821733099306021E-003
wbar=   0.99996005074501004

D=   -1.5122080935215353E-006  Correlation=   -6.0828857026814327E-004
Approx. D=   -2.0073351661345005E-006  Approx. Corr.=
-8.0595134839633604E-004

q relative to approx. equilib. value=   0.99813390147161141

Load statistics for gene model

Relative L=   0.99873137474526985       Relative B=   0.98631036168423269

Relative V x h =   0.39956792650275552

#### Dominance coefficient = 0.5

##### Sites model

Initial state (approximate equilibrium equation; no LD)
x =   2.4906406200928270E-003  y=    6.2343847900086618E-006  q=
2.4968750048828359E-003

Epistasis parameter=    0.0000000000000000

Iteration of haplotype frequencies

Population genetic statistics for gene model
Gen= 5000 x= 2.4875166084561157E-003 y= 6.2100875614190697E-006
q= 2.4937266960175348E-003
wx= 0.99597008148973531 wy= 0.99196510024643092 wz=
0.99998005018643188
wy*wz-wx**2= -1.1092495055042839E-005
wbar= 0.99996005068542615

D= -8.5852730116048326E-009 Correlation= -3.4513549028150712E-006
Approx. D= -8.6178440690873603E-009 Approx. Corr.=
-3.4600913514235425E-006

q relative to approx. equilib. value= 0.99873910033175695

Load statistics for gene model

Relative L= 0.99873286434778652 Relative B= 0.98631180023520826

Relative V x h = 0.39924143927867412

Epistasis parameter= 8.0000000000000002E-002

Iteration of haplotype frequencies

Population genetic statistics for gene model
Gen= 5000 x= 2.4875166241138395E-003 y= 5.7479575626106352E-006
q= 2.4932645816764503E-003
wx= 0.99597007967542839 wy= 0.99132231425511341 wz=
0.99998005020465375
wy*wz-wx**2= -6.5386203085737726E-004
wbar= 0.99996005070364757

D= -4.6841071163136738E-007 Correlation= -1.8834002055416346E-004
Approx. D= -5.0829717458544057E-007 Approx. Corr.=
-2.0408290561265164E-004

q relative to approx. equilib. value= 0.99855402324933162

Load statistics for gene model

Relative L= 0.99873240880968295 Relative B= 0.98631136030929056

Relative V x h = 0.39932309543719191

Epistasis parameter= 0.32000000000000001

Iteration of haplotype frequencies

Population genetic statistics for gene model
Gen= 5000 x= 2.4875166596565845E-003 y= 4.6989304540692557E-006 q= 2.4922155901106538E-003
wx= 0.99597007555698291 wy= 0.98939395626107118 wz= 0.99998005024601710
wy*wz-wx**2= -2.5821733099306021E-003
wbar= 0.99996005074501004

D= -1.5122080935215353E-006 Correlation= -6.0828857026814327E-004
Approx. D= -2.0073351661345005E-006 Approx. Corr.= -8.0595134839633604E-004

q relative to approx. equilib. value= 0.99813390147161141

Load statistics for gene model

Relative L= 0.99873137474526985 Relative B= 0.98631036168423269

Relative V x h = 0.39956792650275552

**Gene model**

Initial state (approximate equilibrium equation; no LD)
x = 1.9959999999999999E-003 y= 3.9999999999999998E-006 q= 2.0000000000000000E-003

Epistasis parameter= 0.0000000000000000

Iteration of haplotype frequencies

Population genetic statistics for sites model
Gen= 5000 x= 1.9959601809935645E-003 y= 3.9898202063947421E-006 q= 1.9999500011999593E-003

wx= 0.99498000049998803 wy= 0.98998000049998802 wz= 0.99998000049998803
wy*wz-wx**2= -2.4999999999941735E-005
wbar= 0.99996000099997606

D= -9.9798009049398284E-009 Correlation= -5.0000250001109753E-006
Approx. D= -1.0000040160642569E-008 Approx. Corr.= -5.0100401606425697E-006

q relative to approx. equilib. value= 0.99997500059997968

Load statistics for sites model

Relative L=   0.99997500059997968       Relative B=    1.0000039898202064

Relative V x h =   0.49799158603713684

Epistasis parameter=    8.0000000000000002E-002

Iteration of haplotype frequencies

Population genetic statistics for sites model
Gen=         5000  x=    1.9953244602772149E-003 y=    3.6920035693305395E-006
q=    1.9990164638465455E-003

wx=   0.99497840914538904       wy=   0.98917841062219047       wz=
0.99998000688175870
wy*wz-wx**2=   -8.2340080422405038E-004
wbar=   0.99996000738161894

D=   -3.0406325339906240E-007  Correlation=   -1.5241110008332040E-004
Approx. D=   -3.1925870949092663E-007  Approx. Corr.=
-1.5994925325196724E-004

q relative to approx. equilib. value=   0.99950823192327276

Load statistics for sites model

Relative L=   0.99981545952391826       Relative B=    1.0000046255536863

Relative V x h =   0.49824429238578616

Epistasis parameter=   0.32000000000000001

Iteration of haplotype frequencies

Population genetic statistics for sites model
Gen=         5000  x=    1.9934221436298670E-003 y=    3.0158125653722038E-006
q=    1.9964379561952391E-003

wx=   0.99497364219367812       wy=   0.98677364701897818       wz=
0.99998002596983782
wy*wz-wx**2=   -3.2186114877640337E-003
wbar=   0.99996002646931637

D=   -9.6995194756501618E-007  Correlation=   -4.8681315999158276E-004
Approx. D=   -1.2758055408310606E-006  Approx. Corr.=
-6.3918113268089205E-004

q relative to approx. equilib. value=   0.99821897809761950

Load statistics for sites model

Relative L=   0.99933826708892104       Relative B=    1.0000065279085097

Relative V x h =   0.49915327123525877
