## Supplementary File S3 for "A gene-based model of fitness and its implications for genetic variation: Linkage disequilibrium"

### FILE S3. SUPPLEMENTARY TABLES AND FIGURES

**Table S1**. Statistical summaries of linkage disequilibrium between selected alleles for the gene and site models, when sites are relatively far apart (800-1000 bases). Selection coefficients followed a gamma distribution with a shape parameter of 0.3 and mean $\bar{\gamma} = 2N\bar{s}$. 1000 selected sites were simulated, with $Nu$ = 0.005 and $Nr$ = 0.01, where $N$ is the population size ($N$ = 1000), and $u$ and $r$ are the mutation and recombination rate per site/generation respectively. The signs of $\bar{D}$and $\bar{\sigma}_d$ were assigned with respect to selected alleles. The means and standard errors (SEs) for 1000 replicate simulations are shown.

| | | ***h*=0.0** | | ***h*=0.2** | | ***h*=0.5** | |
|---|---|---|---|---|---|---|---|
| | | **gene** | **sites** | **gene** | **sites** | **gene** | **sites** |
| $\bar{\gamma} = 2$ | | | | | | | |
| $\bar{D}$ | mean | 0.00001 | 0.00004 | -0.00007 | 0.00000 | -0.00003 | -0.00003 |
| | SE | 0.00004 | 0.00004 | 0.00004 | 0.00004 | 0.00004 | 0.00004 |
| $\bar{\sigma}_d$ | mean | 0.00013 | 0.00036 | -0.00094 | -0.00014 | -0.00045 | -0.00032 |
| | SE | 0.00052 | 0.00043 | 0.00052 | 0.00049 | 0.00053 | 0.00052 |
| $\bar{\gamma} = 20$ | | | | | | | |
| $\bar{D}$ | mean | -0.00003 | -0.00004 | -0.00013 | 0.00000 | -0.00014 | -0.00002 |
| | SE | 0.00005 | 0.00004 | 0.00006 | 0.00005 | 0.00005 | 0.00005 |
| $\bar{\sigma}_d$ | mean | -0.00082 | -0.00020 | -0.00194 | -0.00004 | -0.00185 | -0.00053 |
| | SE | 0.00077 | 0.00050 | 0.00078 | 0.00065 | 0.00076 | 0.00076 |
| $\bar{\gamma} = 100$ | | | | | | | |
| $\bar{D}$ | mean | 0.00009 | -0.00001 | -0.00001 | 0.00010 | -0.00004 | 0.00009 |
| | SE | 0.00006 | 0.00004 | 0.00008 | 0.00006 | 0.00009 | 0.00009 |
| $\bar{\sigma}_d$ | mean | 0.00285 | -0.00023 | 0.00028 | 0.00117 | -0.00066 | 0.00102 |
| | SE | 0.00122 | 0.00060 | 0.00115 | 0.00097 | 0.00114 | 0.00119 |
| $\bar{\gamma} = 1000$ | | | | | | | |
| $\bar{D}$ | mean | 0.00024 | -0.00003 | 0.00031 | 0.00007 | 0.00020 | 0.00007 |
| | SE | 0.00007 | 0.00005 | 0.00010 | 0.00012 | 0.00018 | 0.00013 |
| $\bar{\sigma}_d$ | mean | 0.01217 | -0.00021 | 0.00685 | -0.00031 | 0.00155 | 0.00293 |
| | SE | 0.00346 | 0.00085 | 0.00270 | 0.00158 | 0.00208 | 0.00195 |

**Table S2**. Statistical summaries of linkage disequilibrium between selected alleles for the multiplicative fitness sites model, when sites are relatively close (1-100 bases) far apart (800-1000 bases). Selection coefficients followed a gamma distribution with a shape parameter of 0.3 and mean $\bar{\gamma} = 2N\bar{s}$. The other simulation parameters were the same as in Table S1. The signs of $\bar{D}$ and $\bar{\sigma}_d$ were assigned with respect to selected alleles. The means and standard errors (SEs) for 1000 replicate simulations are shown.

| | | **1-100 bp** | | | **800-1000 bp** | | |
|---|---|---|---|---|---|---|---|
| | | ***h*=0.0** | ***h*=0.2** | ***h*=0.5** | ***h*=0.0** | ***h*=0.2** | ***h*=0.5** |
| $\bar{\gamma} = \mathbf{2}$ | | | | | | | |
| $\bar{D}$ | mean | -0.00008 | -0.00005 | -0.00004 | -0.00004 | -0.00004 | -0.00005 |
| | SE | 0.00005 | 0.00005 | 0.00005 | 0.00004 | 0.00004 | 0.00005 |
| $\bar{\sigma}_d$ | mean | -0.00079 | -0.00055 | -0.00041 | -0.00019 | -0.00025 | -0.00044 |
| | SE | 0.00045 | 0.00050 | 0.00054 | 0.00044 | 0.00048 | 0.00057 |
| $\bar{\gamma} = \mathbf{20}$ | | | | | | | |
| $\bar{D}$ | mean | -0.00013 | -0.00006 | -0.00016 | -0.00005 | 0.00009 | -0.00006 |
| | SE | 0.00004 | 0.00005 | 0.00005 | 0.00004 | 0.00005 | 0.00005 |
| $\bar{\sigma}_d$ | mean | -0.00136 | -0.00062 | -0.00222 | -0.00085 | 0.00138 | -0.00068 |
| | SE | 0.00049 | 0.00064 | 0.00075 | 0.00050 | 0.00066 | 0.00075 |
| $\bar{\gamma} = \mathbf{100}$ | | | | | | | |
| $\bar{D}$ | mean | -0.00012 | -0.00007 | 0.00007 | -0.00001 | 0.00000 | -0.00004 |
| | SE | 0.00004 | 0.00006 | 0.00007 | 0.00004 | 0.00007 | 0.00007 |
| $\bar{\sigma}_d$ | mean | -0.00154 | -0.00055 | 0.00141 | -0.00042 | 0.00038 | 0.00003 |
| | SE | 0.00059 | 0.00086 | 0.00108 | 0.00060 | 0.00091 | 0.00107 |
| $\bar{\gamma} = \mathbf{1000}$ | | | | | | | |
| $\bar{D}$ | mean | 0.00001 | -0.00002 | 0.00004 | -0.00001 | -0.00021 | 0.00012 |
| | SE | 0.00003 | 0.00010 | 0.00015 | 0.00004 | 0.00010 | 0.00018 |
| $\bar{\sigma}_d$ | mean | 0.00051 | -0.00038 | 0.00010 | 0.00029 | -0.00161 | -0.00054 |
| | SE | 0.00071 | 0.00148 | 0.00196 | 0.00098 | 0.00148 | 0.00214 |

**Table S3:** Statistical summaries of linkage disequilibrium under the gene and sites models with a larger simulated population size ($N = 5000$). Scaled selection coefficients followed a gamma distribution with shape parameter 0.3 and a mean of $\bar{\gamma} = 2N\bar{s} = 100$. 1000 selected sites were simulated, with $Nu = 0.005$ and $Nr = 0.01$, where $N$ is the population size, $u$ is the mutation rate per site/generation, and $r$ is the rate of crossing over between adjacent sites. Statistics were calculated for alleles that were less than 100 basepairs apart. The sign of $\bar{\sigma}_d$ and $\bar{D}$ was assigned either by giving a positive sign to cases with excesses over random combinations of pairs of selectively deleterious alleles, or to cases with excesses of combinations of minor alleles (see the Methods section). The means and standard errors (SEs) for 1000 replicate simulations are shown

| | | ***h*=0.0** | | ***h*=0.2** | | ***h*=0.5** | |
|---|---|---|---|---|---|---|---|
| | | **gene** | **sites** | **gene** | **sites** | **gene** | **sites** |
| ***selected alleles*** | | | | | | | |
| ***N=5000*** | | | | | | | |
| $\bar{D}$ | mean | 0.00024 | 0.00001 | -0.00006 | -0.00012 | -0.00001 | -0.00013 |
| | SE | 0.00006 | 0.00004 | 0.00007 | 0.00006 | 0.00007 | 0.00007 |
| $\bar{\sigma}_d$ | mean | 0.00703 | 0.00017 | -0.00063 | -0.00166 | 0.00003 | -0.00138 |
| | SE | 0.00108 | 0.00060 | 0.00106 | 0.00081 | 0.00101 | 0.00103 |
| ***N=1000*** | | | | | | | |
| $\bar{D}$ | mean | 0.00014 | -0.00007 | 0.00015 | -0.00007 | 0.00004 | -0.00005 |
| | SE | 0.00005 | 0.00004 | 0.00007 | 0.00006 | 0.00007 | 0.00007 |
| $\bar{\sigma}_d$ | mean | 0.00540 | -0.00082 | 0.00241 | -0.00088 | 0.00089 | -0.00054 |
| | SE | 0.00100 | 0.00059 | 0.00103 | 0.00084 | 0.00103 | 0.00107 |
| ***minor alleles*** | | | | | | | |
| ***N=5000*** | | | | | | | |
| $\bar{D}$ | mean | 0.00123 | 0.00175 | 0.00173 | 0.00208 | 0.00206 | 0.00190 |
| | SE | 0.00008 | 0.00007 | 0.00010 | 0.00010 | 0.00011 | 0.00011 |
| $\bar{\sigma}_d$ | mean | 0.02218 | 0.02614 | 0.02559 | 0.03086 | 0.02992 | 0.02859 |
| | SE | 0.00124 | 0.00095 | 0.00139 | 0.00132 | 0.00149 | 0.00149 |
| ***N=1000*** | | | | | | | |
| $\bar{D}$ | mean | 0.00106 | 0.00175 | 0.00185 | 0.00206 | 0.00208 | 0.00194 |
| | SE | 0.00008 | 0.00007 | 0.00010 | 0.00010 | 0.00011 | 0.00011 |
| $\bar{\sigma}_d$ | mean | 0.01911 | 0.02636 | 0.02730 | 0.03011 | 0.03042 | 0.02872 |
| | SE | 0.00120 | 0.00094 | 0.00138 | 0.00129 | 0.00148 | 0.00145 |

**Table S4**. Statistical summaries of linkage disequilibrium between selected alleles less than 100 bp apart for the gene and sites models in human-like populations. 2000 selected sites were simulated, with *Nu* = 0.00025 and *Nr* = 0.0002, where *N* is the population size (*N* = 1000) and *u* and *r* are the mutation and recombination rate per site/generation respectively. Here $\bar{D}$, $\bar{\sigma}_d$, and $\bar{\sigma}_d^1$ were calculated for alleles that were less than 100 sites apart. The sign of the LD statisic $\bar{\sigma}_d$ and $\bar{\sigma}_d^1$ was assigned either by giving a positive sign to cases with excesses over random combinations of pairs of selectively deleterious alleles or to cases with excesses of combinations of minor alleles. The means and standard errors (SEs) for 1000 replicate simulations are shown.

| $\bar{\gamma} = 850$ | | ***h*=0.0** | | ***h*=0.2** | | ***h*=0.5** | |
|---|---|---|---|---|---|---|---|
| | | **gene** | **sites** | **gene** | **sites** | **gene** | **sites** |
| ***selected allele*** | | | | | | | |
| $\bar{D}$ | mean | 0.00071 | 0.00135 | 0.00193 | -0.00023 | -0.00204 | 0.00086 |
| | SE | 0.00106 | 0.00085 | 0.00177 | 0.00150 | 0.00184 | 0.00221 |
| $\bar{\sigma}_d$ | mean | 0.02485 | 0.01463 | 0.01709 | 0.00496 | -0.00066 | 0.01614 |
| | SE | 0.01230 | 0.00717 | 0.01387 | 0.01153 | 0.01259 | 0.01512 |
| $\bar{\sigma}_d^1$ | mean | 1.55314 | 0.39533 | 0.10215 | 0.52399 | 0.27492 | 0.37814 |
| | SE | 0.68573 | 0.28496 | 0.41651 | 0.36267 | 0.36166 | 0.38685 |
| ***minor allele*** | | | | | | | |
| $\bar{D}$ | mean | 0.00186 | 0.00119 | 0.00576 | 0.00184 | 0.00325 | 0.00596 |
| | SE | 0.00105 | 0.00085 | 0.00179 | 0.00149 | 0.00186 | 0.00217 |
| $\bar{\sigma}_d$ | mean | 0.03674 | 0.01776 | 0.05261 | 0.02340 | 0.02893 | 0.04956 |
| | SE | 0.01219 | 0.00712 | 0.01375 | 0.01141 | 0.01273 | 0.01479 |
| $\bar{\sigma}_d^1$ | mean | 1.70886 | 0.47674 | 0.98834 | 0.75829 | 0.46001 | 0.89475 |
| | SE | 0.68462 | 0.28473 | 0.41330 | 0.36086 | 0.36180 | 0.38415 |

**Table S5**. The numbers of segregating sites for the gene and sites models used for Table 2. 1000 selected sites were simulated, with $Nu$ = 0.005 and $Nr$ = 0.01, where $N$ is the population size ($N$ = 1000), $u$ is the mutation rate per site/generation, and $r$ is the rate of crossing over between adjacent sites. The means and standard errors (SEs) for 1000 replicate simulations are shown.

| | | **$h$=0.0** | | **$h$=0.2** | | **$h$=0.5** | |
|---|---|---|---|---|---|---|---|
| | | **Gene model** | **Sites model** | **Gene model** | **Sites model** | **Gene model** | **Sites model** |
| $\gamma = 2$ | mean | 81.67 | 118.40 | 82.55 | 99.23 | 82.47 | 82.91 |
| | SE | 0.36 | 0.51 | 0.37 | 0.44 | 0.36 | 0.38 |
| $\gamma = 20$ | mean | 35.34 | 85.16 | 35.33 | 51.24 | 36.13 | 35.88 |
| | SE | 0.19 | 0.42 | 0.20 | 0.26 | 0.21 | 0.20 |
| $\gamma = 100$ | mean | 20.82 | 56.37 | 17.92 | 24.75 | 14.03 | 14.01 |
| | SE | 0.14 | 0.29 | 0.13 | 0.16 | 0.12 | 0.12 |
| $\bar{\gamma} = 2$ | mean | 86.05 | 106.57 | 86.60 | 95.24 | 86.16 | 86.61 |
| | SE | 0.38 | 0.48 | 0.38 | 0.43 | 0.38 | 0.38 |
| $\bar{\gamma} = 20$ | mean | 60.31 | 89.81 | 59.67 | 70.51 | 60.18 | 59.93 |
| | SE | 0.30 | 0.42 | 0.29 | 0.33 | 0.31 | 0.30 |
| $\bar{\gamma} = 100$ | mean | 37.38 | 72.48 | 42.39 | 51.36 | 42.55 | 42.31 |
| | SE | 0.27 | 0.34 | 0.24 | 0.27 | 0.24 | 0.24 |
| $\bar{\gamma} = 1000$ | mean | 15.10 | 51.28 | 19.06 | 29.31 | 23.23 | 23.72 |
| | SE | 0.13 | 0.26 | 0.16 | 0.19 | 0.17 | 0.16 |

**Figure S1:** Statistical summaries of the coefficient of linkage disequilibrium (*D*) under the gene and sites models for a gamma distribution of scaled selection coefficients with shape parameter 0.3 and a mean of $\bar{\gamma} = 2N\bar{s}$. 1000 selected sites were simulated, with $Nu$ = 0.005 and $Nr$ = 0.01, where *N* is the population size (*N* = 1000), *u* is the mutation rate per site/generation, and *r* is the rate of crossing over between adjacent sites. $\bar{D}$ was calculated for alleles that were less than 100 basepairs apart. The sign of $\bar{D}$ was assigned either by giving a positive sign to cases with excesses over random combinations of pairs of selectively deleterious alleles, or to cases with excesses of combinations of minor alleles (see the Methods section). The means and standard errors (SEs) for 1000 replicate simulations are shown.

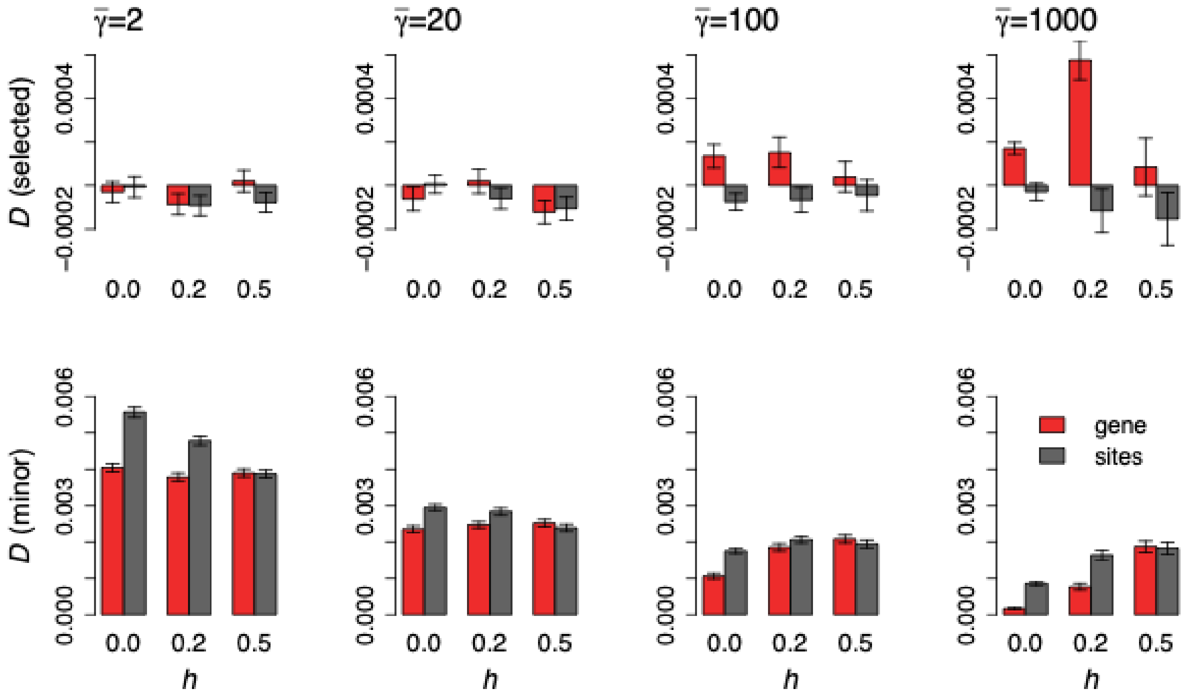

**Figure S2**: Statistical summaries of linkage disequilibrium (mean *D*) between tightly linked alleles (1-100 sites apart) for the gene and sites models with epistasis. Simulations were performed as in Figure S1.

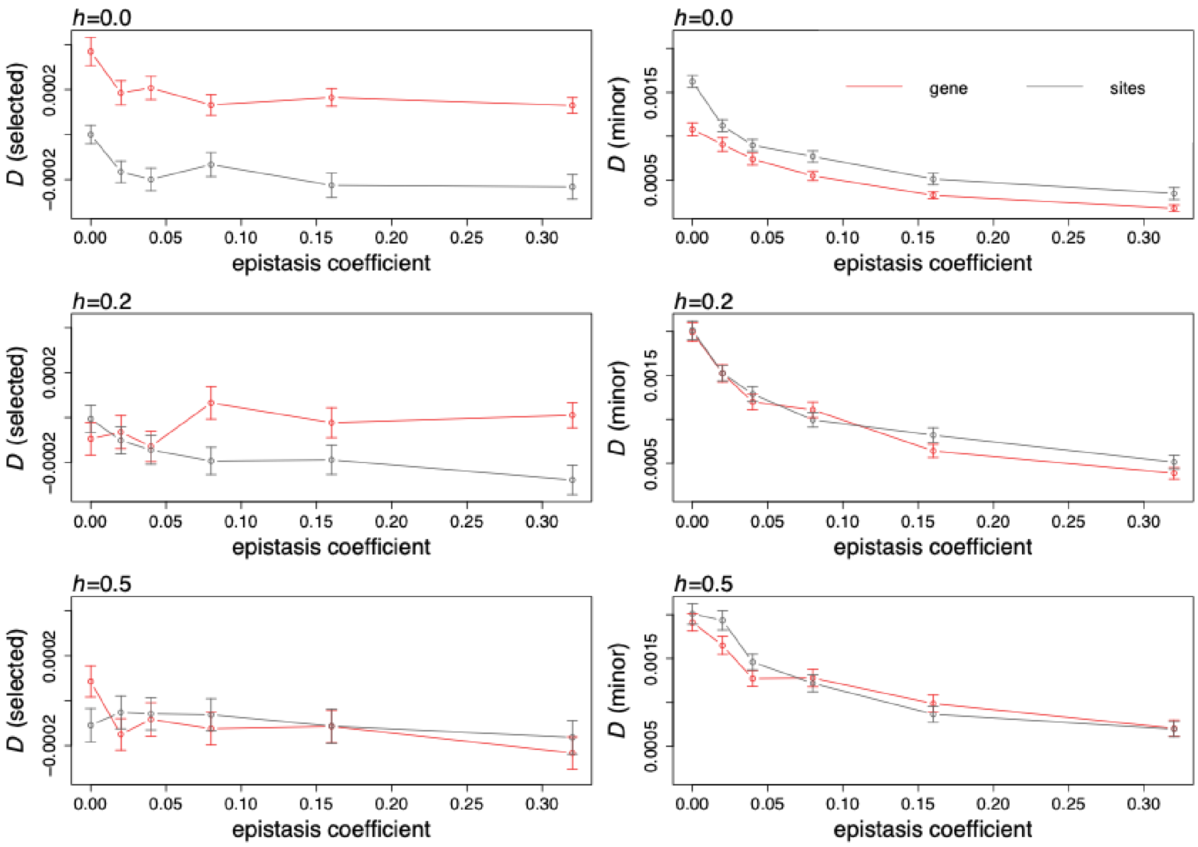

**Figure S3**: The distribution of $\bar{D}$ ("mean D") and $\bar{\sigma}_d$ ("mean sigmaD") across 1000 replicates is presented for the gene model, when $h = 0.2$ and when there is no epistasis. The distribution shown in black was obtained when simulations were run using the standard SLiM script, which did not model any epistasis. The distribution in red was obtained from simulations where the SLiM script included epistasis and $\epsilon = 0.0$. The two simulations yielded a nearly identical distribution of LD measures across replicates, even though the mean value appears to be different in Table 4 and Figure 5, suggesting that the differences in means mostly represent noise.

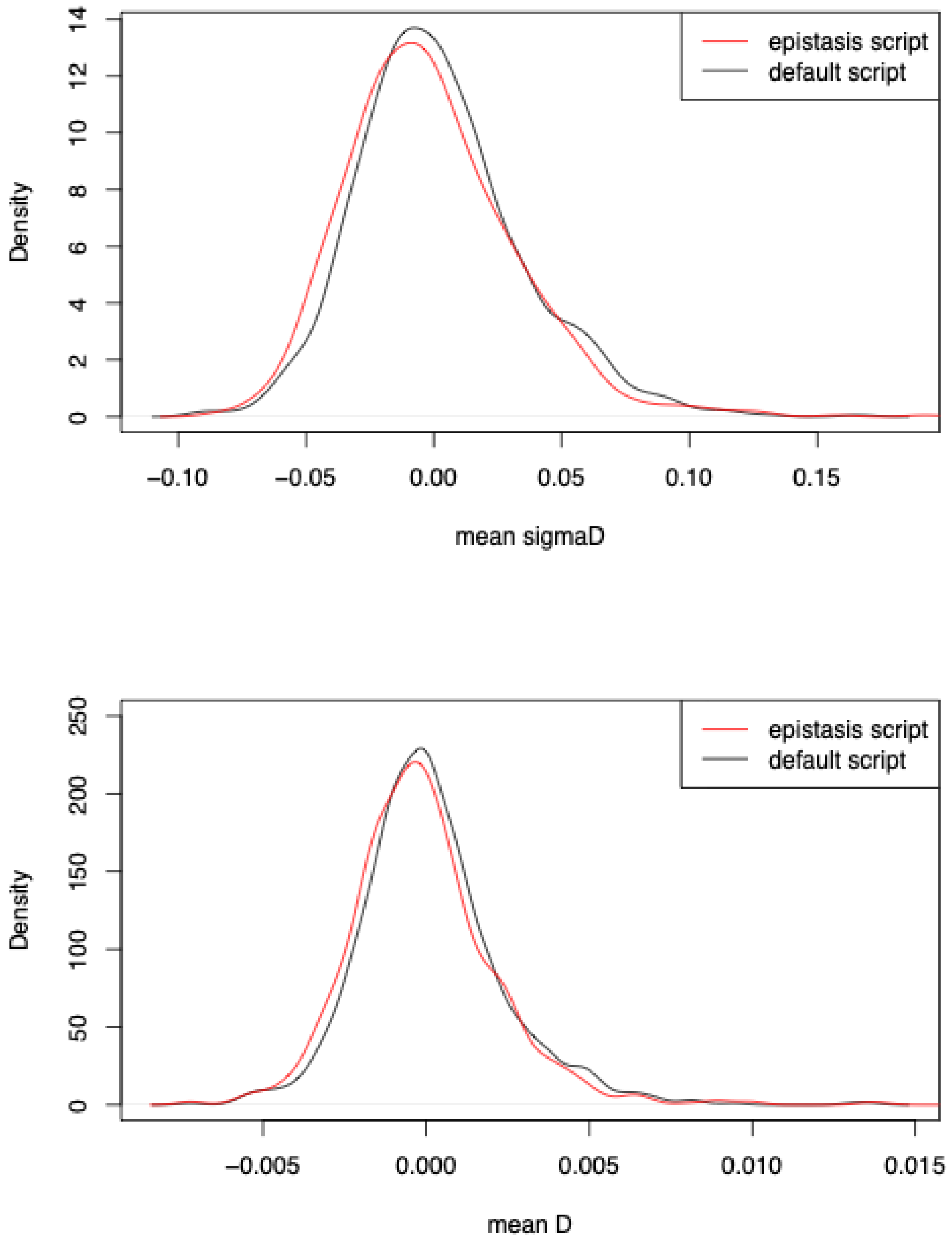

**Figure S4**: Statistical summaries of linkage disequilibrium ($\bar{\sigma}_d$) between tightly linked alleles (1-100 sites apart) for the gene and sites models with epistasis when the rate of crossing over is low. Simulations were performed as for Figure S2.

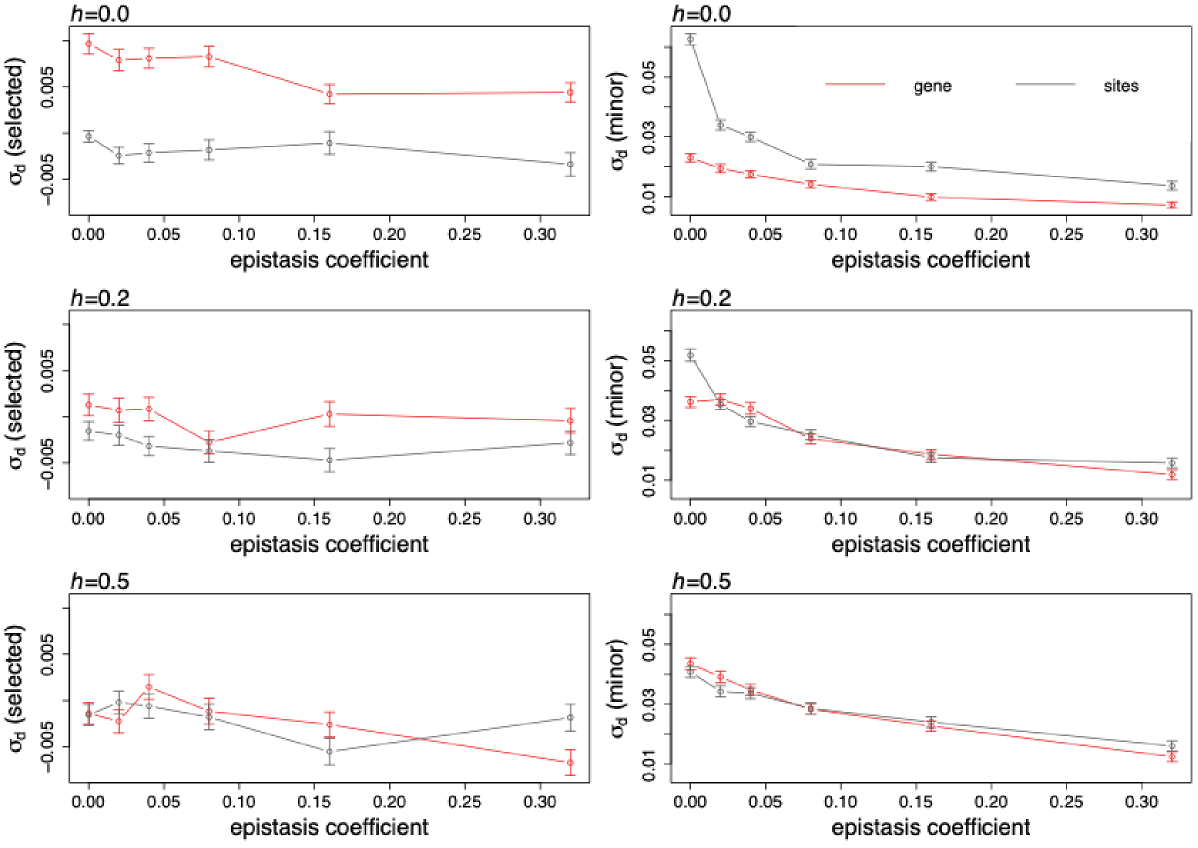

**Figure S5**: Statistical summaries of linkage disequilibrium (mean *D*) between tightly linked alleles (1-100 sites apart) for the gene and sites models with epistasis when the rate of crossing over is low. Simulations were performed as for Figure S4.

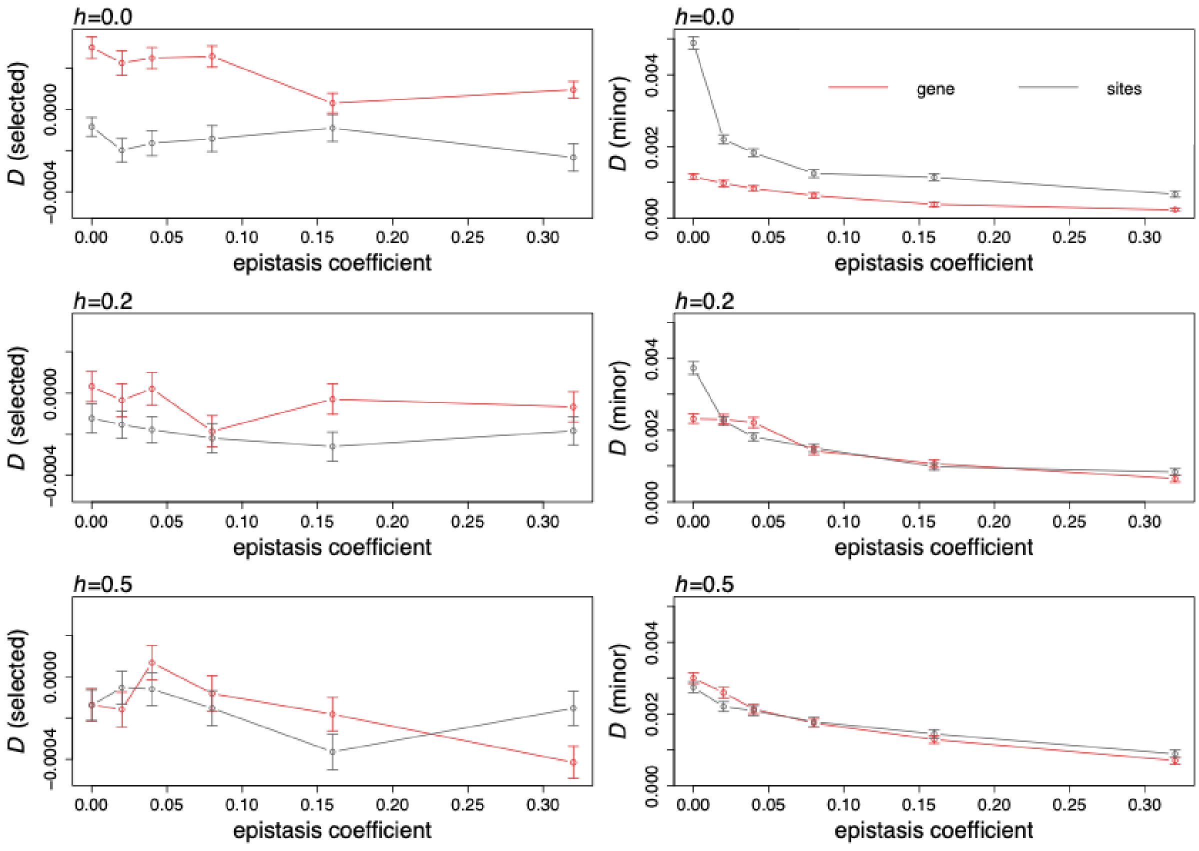
