## Supplementary File S4 for "A gene-based model of fitness and its implications for genetic variation: Linkage disequilibrium"

**Simulation Results for 2-locus LD with no Recombination**

**Finite sample size with Garcia-Lohmueller method of fixing derived allele frequency in sample**

**Size of haploid population= 1000**
**Sample size= 200**
**Sample frequency selection= 2**
**Number of replications= 1000000**

**Selection coefficient= 0.00000000**
**Scaled selection coefficient= 0.00000000**
**Epistasis coefficient= 0.0000000**

Summary statistics for whole population

ns1= 134401 ps1= 0.134400994
ns2= 865599 ps2= 0.865598977

Mean allele frequencies at 1st locus, conditioned on segregation
q1bar1= 0.458536416 q1bar2= 0.128767341 q1bar= 0.200937837
Mean allele frequencies at 2nd locus, conditioned on segregation
q2bar1= 9.32425335E-02 q2bar2= 2.61234380E-02 q2bar=
4.08125632E-02

Mean sojourn times
tbar1= 12.9221582 tbar2= 7.16150904 tbar= 7.93574619

D statistics
Mean D1= 3.66170555E-02 Mean D2= -1.04694292E-02 Mean D=
-1.64474826E-04
Mean D1^2= 5.41648269E-03 Mean D2^2= 1.33841485E-03 Mean D^2=
2.23090686E-03

Mean pr1= 2.07382930E-03 Mean pr2= 2.91529531E-03 Mean pr=
4.98912483E-03
sig1d1= 17.6567364 sig1d2= -3.59120703 sig1d=
-3.29666696E-02
sigd1= 0.804075658 sigd2= -0.193901747 sigd= -2.32855906E-03
sig2d1= 2.61182666 sig2d2= 0.459100962 sig2d= 0.447153956
Theoretical value of unweighted sigma2d= 0.454544991

Sums of generation times for samples which match chosen allele frequency

tsamp1= 5261 tsamp2= 31687 tsamp= 36948
Mean allele frequencies at 1st locus, conditioned on segregation
q1bar1= 9.99946333E-03 q1bar2= 9.99992993E-03 q1bar=
9.99986287E-03
Mean allele frequencies at 2nd locus, conditioned on segregation
q2bar1= 9.99946333E-03 q2bar2= 9.99993552E-03 q2bar=
9.99986846E-03

D statistics
Mean D1= 9.89942811E-03 Mean D2= -9.99690310E-05 Mean D=
1.32383825E-03
Mean D1^2= 9.80131517E-05 Mean D2^2= 6.02369923E-08 Mean D^2=
1.39646017E-05

Mean pr1= 9.80131517E-05 Mean pr2= 9.79938704E-05 Mean pr=
9.79966208E-05
sig1d1= 101.001015 sig1d2= -1.02015591 sig1d= 13.5090199
sigd1= 0.999926150 sigd2= -1.00987135E-02 sigd= 0.133730173
sig2d1= 1.00000000 sig2d2= 6.14701654E-04 sig2d= 0.142500848

**Selection coefficient= 4.99999989E-03**
**Scaled selection coefficient= 10.0000000**
**Epistasis coefficient= 0.00000000**

Summary statistics for whole population

ns1= 23675 ps1= 2.36750003E-02
ns2= 976325 ps2= 0.976324975

Mean allele frequencies at 1st locus, conditioned on segregation
q1bar1= 0.101496592 q1bar2= 2.94015482E-02 q1bar= 3.22520956E-02
Mean allele frequencies at 2nd locus, conditioned on segregation
q2bar1= 1.32668875E-02 q2bar2= 5.46150433E-04 q2bar= 1.04911264E-03

Mean sojourn times
tbar1= 8.42432976 tbar2= 4.96235895 tbar= 5.04432106

D statistics
Mean D1= 1.15246158E-02 Mean D2= -4.93895495E-04 Mean D=
-1.86988982E-05
Mean D1^2= 5.10663434E-04 Mean D2^2= 7.70242332E-06 Mean D^2=
2.75888578E-05

Mean pr1= 5.03871415E-05 Mean pr2= 3.72321578E-04 Mean pr=
4.22708690E-04
sig1d1= 228.721359 sig1d2= -1.32652938 sig1d= -4.42358963E-02

sigd1= 1.62355340 sigd2= -2.55962275E-02 sigd= -9.09484748E-04
sig2d1= 10.1347971 sig2d2= 2.06875559E-02 sig2d= 6.52668327E-02
Theoretical value of unweighted sigma2d= 0.454544991

Summary statistics for sample

Sums of generation times for samples which match chosen allele frequency
tsamp1= 3540 tsamp2= 42817 tsamp= 46357
Mean allele frequencies at 1st locus, conditioned on segregation
q1bar1= 1.00000184E-02 q1bar2= 1.00024864E-02 q1bar= 1.00022973E-02
Mean allele frequencies at 2nd locus, conditioned on segregation
q2bar1= 1.00000184E-02 q2bar2= 1.00024911E-02 q2bar= 1.00023020E-02

D statistics
Mean D1= 9.89956874E-03 Mean D2= -9.99659896E-05 Mean D=
6.63637184E-04
Mean D1^2= 9.80112454E-05 Mean D2^2= 1.20999218E-07 Mean D^2=
7.49375840E-06

Mean pr1= 9.80112454E-05 Mean pr2= 9.80039986E-05 Mean pr=
9.80045443E-05
sig1d1= 101.004417 sig1d2= -1.02001953 sig1d= 6.77149391
sigd1= 0.999950051 sigd2= -1.00978836E-02 sigd= 6.70359284E-02
sig2d1= 1.00000000 sig2d2= 1.23463548E-03 sig2d= 7.64633790E-02

**Selection coefficient= 4.99999989E-03**
**Scaled selection coefficient= 10.0000000**
**Epistasis coefficient= 1.99999996E-02**

Summary statistics for whole population

ns1= 23733 ps1= 2.37329993E-02
ns2= 976267 ps2= 0.976266980

Mean allele frequencies at 1st locus, conditioned on segregation
q1bar1= 9.79016200E-02 q1bar2= 2.97417268E-02 q1bar= 3.24252434E-02
Mean allele frequencies at 2nd locus, conditioned on segregation
q2bar1= 1.28182275E-02 q2bar2= 5.25348936E-04 q2bar= 1.00933097E-03

Mean sojourn times
tbar1= 8.38777256 tbar2= 4.97520447 tbar= 5.05619478

D statistics
Mean D1= 1.12251323E-02 Mean D2= -4.88664373E-04 Mean D=
-2.74815156E-05
Mean D1^2= 4.87699668E-04 Mean D2^2= 5.34894616E-06 Mean D^2=
2.43395316E-05

Mean pr1= 4.50242806E-05 Mean pr2= 3.76034819E-04 Mean pr=

4.21059114E-04
sig1d1= 249.312866 sig1d2= -1.29951894 sig1d= -6.52675927E-02
sigd1= 1.67289269 sigd2= -2.51997747E-02 sigd= -1.33927318E-03
sig2d1= 10.8319254 sig2d2= 1.42246038E-02 sig2d= 5.78054972E-02
Theoretical value of unweighted sigma2d= 0.454544991

Summary statistics for sample

Sums of generation times for samples which match chosen allele frequency
tsamp1= 3792 tsamp2= 42850 tsamp= 46642
Mean allele frequencies at 1st locus, conditioned on segregation
q1bar1= 9.99990571E-03 q1bar2= 1.00024920E-02 q1bar= 1.00022815E-02
Mean allele frequencies at 2nd locus, conditioned on segregation
q2bar1= 9.99990571E-03 q2bar2= 1.00024967E-02 q2bar= 1.00022862E-02

D statistics
Mean D1= 9.89953987E-03 Mean D2= -9.99661206E-05 Mean D=
7.12994894E-04
Mean D1^2= 9.80118130E-05 Mean D2^2= 1.13045253E-07 Mean D^2=
7.97756275E-06

Mean pr1= 9.80118130E-05 Mean pr2= 9.80041659E-05 Mean pr=
9.80047917E-05
sig1d1= 101.003540 sig1d2= -1.02001905 sig1d= 7.27510262
sigd1= 0.999944329 sigd2= -1.00978883E-02 sigd= 7.20215961E-02
sig2d1= 1.00000000 sig2d2= 1.15347398E-03 sig2d= 8.13997239E-02

**Selection coefficient= 4.99999989E-03**
**Scaled selection coefficient= 10.0000000**
**Epistasis coefficient= 3.99999991E-02**

Summary statistics for whole population

ns1= 23651 ps1= 2.36510001E-02
ns2= 976349 ps2= 0.976348996

Mean allele frequencies at 1st locus, conditioned on segregation
q1bar1= 9.89011303E-02 q1bar2= 2.98025366E-02 q1bar=
3.24925072E-02
Mean allele frequencies at 2nd locus, conditioned on segregation
q2bar1= 1.35665676E-02 q2bar2= 5.49532298E-04 q2bar=
1.05627859E-03

Mean sojourn times
tbar1= 8.27487183 tbar2= 4.94860363 tbar= 5.02727318

D statistics

Mean D1= 1.17372554E-02 Mean D2= -5.32956794E-04 Mean D=
-5.52841411E-05
Mean D1^2= 5.63709415E-04 Mean D2^2= 1.21710846E-05 Mean D^2=
3.36421726E-05

Mean pr1= 5.09751735E-05 Mean pr2= 3.86260246E-04 Mean pr=
4.37235431E-04
sig1d1= 230.254349 sig1d2= -1.37978685 sig1d= -0.126440212
sigd1= 1.64394462 sigd2= -2.71176472E-02 sigd= -2.64388695E-03
sig2d1= 11.0585089 sig2d2= 3.15100625E-02 sig2d=
7.69429207E-02
Theoretical value of unweighted sigma2d= 0.454544991

Summary statistics for sample

Sums of generation times for samples which match chosen allele frequency
tsamp1= 3561 tsamp2= 42650 tsamp= 46211
Mean allele frequencies at 1st locus, conditioned on segregation
q1bar1= 1.00000091E-02 q1bar2= 1.00024575E-02 q1bar=
1.00022685E-02
Mean allele frequencies at 2nd locus, conditioned on segregation
q2bar1= 1.00000091E-02 q2bar2= 1.00024622E-02 q2bar=
1.00022731E-02

D statistics
Mean D1= 9.89956595E-03 Mean D2= -9.99653275E-05 Mean D=
6.70594280E-04
Mean D1^2= 9.80112964E-05 Mean D2^2= 1.19816136E-07 Mean D^2=
7.56194186E-06

Mean pr1= 9.80112964E-05 Mean pr2= 9.80031182E-05 Mean pr=
9.80037439E-05
sig1d1= 101.004333 sig1d2= -1.02002192 sig1d= 6.84253740
sigd1= 0.999949574 sigd2= -1.00978622E-02 sigd= 6.77389577E-02
sig2d1= 1.00000000 sig2d2= 1.22257473E-03 sig2d=
7.71597251E-02

**Selection coefficient= 4.99999989E-03**
**Scaled selection coefficient= 10.0000000**
**Epistasis coefficient= 7.99999982E-02**

Summary statistics for whole population

ns1= 23585 ps1= 2.35849991E-02
ns2= 976415 ps2= 0.976414979

Mean allele frequencies at 1st locus, conditioned on segregation
q1bar1= 0.103102289 q1bar2= 2.95502543E-02 q1bar= 3.24335657E-02
Mean allele frequencies at 2nd locus, conditioned on segregation
q2bar1= 1.25726713E-02 q2bar2= 5.12970088E-04 q2bar= 9.85722290E-04

Mean sojourn times
tbar1= 8.40898895 tbar2= 4.97829819 tbar= 5.05921078

D statistics
Mean D1= 1.07557606E-02 Mean D2= -4.74531698E-04 Mean D=
-3.42933054E-05
Mean D1^2= 4.33419598E-04 Mean D2^2= 5.82593566E-06 Mean D^2=
2.25880249E-05

Mean pr1= 4.86448553E-05 Mean pr2= 3.63263855E-04 Mean pr=
4.11908724E-04
sig1d1= 221.107880 sig1d2= -1.30630040 sig1d= -8.32546204E-02
sigd1= 1.54213595 sigd2= -2.48974096E-02 sigd= -1.68969715E-03
sig2d1= 8.90987492 sig2d2= 1.60377529E-02 sig2d= 5.48374504E-02
Theoretical value of unweighted sigma2d= 0.454544991

Summary statistics for sample

Sums of generation times for samples which match chosen allele frequency
tsamp1= 3825 tsamp2= 43021 tsamp= 46846
Mean allele frequencies at 1st locus, conditioned on segregation
q1bar1= 9.99989267E-03 q1bar2= 1.00025209E-02 q1bar= 1.00023067E-02
Mean allele frequencies at 2nd locus, conditioned on segregation
q2bar1= 9.99989267E-03 q2bar2= 1.00025255E-02 q2bar= 1.00023104E-02

D statistics
Mean D1= 9.89953615E-03 Mean D2= -9.99667973E-05 Mean D=
7.16497772E-04
Mean D1^2= 9.80118857E-05 Mean D2^2= 1.12517547E-07 Mean D^2=
8.01190799E-06

Mean pr1= 9.80118857E-05 Mean pr2= 9.80050609E-05 Mean pr=
9.80056211E-05
sig1d1= 101.003426 sig1d2= -1.02001667 sig1d= 7.31078243
sigd1= 0.999943554 sigd2= -1.00979107E-02 sigd= 7.23751262E-02
sig2d1= 1.00000000 sig2d2= 1.14807894E-03 sig2d= 8.17494765E-02

**Selection coefficient= 4.99999989E-03**
**Scaled selection coefficient= 10.0000000**
**Epistasis coefficient= 0.159999996**

Summary statistics for whole population

ns1= 24040 ps1= 2.40400005E-02
ns2= 975960 ps2= 0.975960016

Mean allele frequencies at 1st locus, conditioned on segregation
q1bar1= 0.100298353 q1bar2= 2.91881543E-02 q1bar= 3.19123492E-02
Mean allele frequencies at 2nd locus, conditioned on segregation
q2bar1= 1.20213954E-02 q2bar2= 4.78879141E-04 q2bar= 9.21067083E-04

Mean sojourn times
tbar1= 7.96530771 tbar2= 4.92531681 tbar= 4.99839783

D statistics
Mean D1= 1.02526639E-02 Mean D2= -4.46878927E-04 Mean D=
-3.69850677E-05
Mean D1^2= 4.34124202E-04 Mean D2^2= 4.71828798E-06 Mean D^2=
2.11686038E-05

Mean pr1= 4.60499577E-05 Mean pr2= 3.45606444E-04 Mean pr=
3.91656387E-04
sig1d1= 222.642197 sig1d2= -1.29302835 sig1d= -9.44324359E-02
sigd1= 1.51085258 sigd2= -2.40380354E-02 sigd= -1.86884729E-03
sig2d1= 9.42724419 sig2d2= 1.36521989E-02 sig2d= 5.40489182E-02
Theoretical value of unweighted sigma2d= 0.454544991

Summary statistics for sample

Sums of generation times for samples which match chosen allele frequency
tsamp1= 3134 tsamp2= 42498 tsamp= 45632
Mean allele frequencies at 1st locus, conditioned on segregation
q1bar1= 1.00001981E-02 q1bar2= 1.00024315E-02 q1bar= 1.00022778E-02
Mean allele frequencies at 2nd locus, conditioned on segregation
q2bar1= 1.00001981E-02 q2bar2= 1.00024361E-02 q2bar= 1.00022824E-02

D statistics
Mean D1= 9.89962369E-03 Mean D2= -9.99647164E-05 Mean D=
5.86805749E-04
Mean D1^2= 9.80101322E-05 Mean D2^2= 1.35655227E-07 Mean D^2=
6.74063995E-06

Mean pr1= 9.80101322E-05 Mean pr2= 9.80023106E-05 Mean pr=
9.80028490E-05
sig1d1= 101.006126 sig1d2= -1.02002406 sig1d= 5.98763990
sigd1= 0.999961317 sigd2= -1.00978427E-02 sigd= 5.92754707E-02
sig2d1= 1.00000000 sig2d2= 1.38420437E-03 sig2d= 6.87800422E-02

**Selection coefficient= 4.99999989E-03**
**Scaled selection coefficient= 10.0000000**

**Epistasis coefficient= 0.319999993**

Summary statistics for whole population

ns1= 23830 ps1= 2.38300003E-02
ns2= 976170 ps2= 0.976170003

Mean allele frequencies at 1st locus, conditioned on segregation
q1bar1= 9.96260270E-02 q1bar2= 2.98742130E-02 q1bar= 3.24128196E-02
Mean allele frequencies at 2nd locus, conditioned on segregation
q2bar1= 1.02070021E-02 q2bar2= 3.85513325E-04 q2bar= 7.42965261E-04

Mean sojourn times
tbar1= 7.71716309 tbar2= 4.98787212 tbar= 5.05291080

D statistics
Mean D1= 8.82092584E-03 Mean D2= -5.14713989E-04 Mean D=
-1.74944667E-04
Mean D1^2= 2.93892750E-04 Mean D2^2= 7.92670107E-06 Mean D^2=
1.83343964E-05

Mean pr1= 3.56786259E-05 Mean pr2= 3.86949367E-04 Mean pr=
4.22628014E-04
sig1d1= 247.232773 sig1d2= -1.33018434 sig1d= -0.413944811
sigd1= 1.47676063 sigd2= -2.61660945E-02 sigd= -8.50984361E-03
sig2d1= 8.23722172 sig2d2= 2.04851124E-02 sig2d= 4.33818772E-02
Theoretical value of unweighted sigma2d= 0.454544991

Summary statistics for sample

Sums of generation times for samples which match chosen allele frequency
tsamp1= 3157 tsamp2= 43103 tsamp= 46260
Mean allele frequencies at 1st locus, conditioned on segregation
q1bar1= 1.00001991E-02 q1bar2= 1.00025348E-02 q1bar= 1.00023746E-02
Mean allele frequencies at 2nd locus, conditioned on segregation
q2bar1= 1.00001991E-02 q2bar2= 1.00025386E-02 q2bar= 1.00023793E-02

D statistics
Mean D1= 9.89961997E-03 Mean D2= -9.99671174E-05 Mean D=
5.82451758E-04
Mean D1^2= 9.80101977E-05 Mean D2^2= 1.36585555E-07 Mean D^2=
6.69799829E-06

Mean pr1= 9.80101977E-05 Mean pr2= 9.80054829E-05 Mean pr=
9.80058103E-05
sig1d1= 101.006020 sig1d2= -1.02001560 sig1d= 5.94303274
sigd1= 0.999960601 sigd2= -1.00979218E-02 sigd= 5.88347688E-02
sig2d1= 1.00000000 sig2d2= 1.39365217E-03 sig2d= 6.83428720E-02

**Selection coefficient= 4.99999989E-03**
**Scaled selection coefficient= 10.0000000**
**Epistasis coefficient= 0.639999986**

Summary statistics for whole population

ns1= 23827 ps1= 2.38269996E-02
ns2= 976173 ps2= 0.976172984

Mean allele frequencies at 1st locus, conditioned on segregation
q1bar1= 9.38053653E-02 q1bar2= 2.93177180E-02 q1bar= 3.15042622E-02
Mean allele frequencies at 2nd locus, conditioned on segregation
q2bar1= 8.07556044E-03 q2bar2= 2.83423171E-04 q2bar= 5.47626638E-04

Mean sojourn times
tbar1= 7.12704086 tbar2= 4.95666361 tbar= 5.00837708

D statistics
Mean D1= 7.15769455E-03 Mean D2= -4.78636241E-04 Mean D=
-2.19715803E-04
Mean D1^2= 1.91703366E-04 Mean D2^2= 7.14821499E-06 Mean D^2=
1.34058146E-05

Mean pr1= 2.38641023E-05 Mean pr2= 3.64738109E-04 Mean pr=
3.88602202E-04
sig1d1= 299.935638 sig1d2= -1.31227374 sig1d= -0.565400302
sigd1= 1.46521258 sigd2= -2.50619594E-02 sigd= -1.11457342E-02
sig2d1= 8.03312683 sig2d2= 1.95982121E-02 sig2d= 3.44975255E-02
Theoretical value of unweighted sigma2d= 0.454544991

Summary statistics for sample

Sums of generation times for samples which match chosen allele frequency
tsamp1= 2849 tsamp2= 42843 tsamp= 45692
Mean allele frequencies at 1st locus, conditioned on segregation
q1bar1= 1.00001954E-02 q1bar2= 1.00024911E-02 q1bar= 1.00023476E-02
Mean allele frequencies at 2nd locus, conditioned on segregation
q2bar1= 1.00001954E-02 q2bar2= 1.00024948E-02 q2bar= 1.00023523E-02

D statistics
Mean D1= 9.89967212E-03 Mean D2= -9.99660915E-05 Mean D=
5.23534080E-04
Mean D1^2= 9.80091572E-05 Mean D2^2= 1.50437884E-07 Mean D^2=
6.12047370E-06

Mean pr1= 9.80091572E-05 Mean pr2= 9.80041295E-05 Mean pr=
9.80044497E-05
sig1d1= 101.007622 sig1d2= -1.02001917 sig1d= 5.34194183

sigd1= 0.999971211 sigd2= -1.00978874E-02 sigd= 5.28837293E-02
sig2d1= 1.00000000 sig2d2= 1.53501576E-03 sig2d= 6.24509789E-02

**Selection coefficient= 4.99999989E-03**
**Scaled selection coefficient= 10.0000000**
**Epistasis coefficient= 1.27999997**

Summary statistics for whole population

ns1= 23782 ps1= 2.37819999E-02
ns2= 976218 ps2= 0.976217985

Mean allele frequencies at 1st locus, conditioned on segregation
q1bar1= 9.69558880E-02 q1bar2= 2.98722275E-02 q1bar= 3.19410115E-02
Mean allele frequencies at 2nd locus, conditioned on segregation
q2bar1= 6.63331198E-03 q2bar2= 2.11073129E-04 q2bar= 4.09127766E-04

Mean sojourn times
tbar1= 6.47519112 tbar2= 4.95737839 tbar= 4.99347496

D statistics
Mean D1= 5.83804678E-03 Mean D2= -5.07823424E-04 Mean D=
-3.12124117E-04
Mean D1^2= 1.30138680E-04 Mean D2^2= 8.15052681E-06 Mean D^2=
1.19124998E-05

Mean pr1= 1.81595060E-05 Mean pr2= 3.83686478E-04 Mean pr=
4.01845988E-04
sig1d1= 321.487091 sig1d2= -1.32353747 sig1d= -0.776725709
sigd1= 1.36998415 sigd2= -2.59253401E-02 sigd= -1.55703193E-02
sig2d1= 7.16642189 sig2d2= 2.12426744E-02 sig2d= 2.96444409E-02
Theoretical value of unweighted sigma2d= 0.454544991

Summary statistics for sample

Sums of generation times for samples which match chosen allele frequency
tsamp1= 2293 tsamp2= 42615 tsamp= 44908
Mean allele frequencies at 1st locus, conditioned on segregation
q1bar1= 1.00001870E-02 q1bar2= 1.00024519E-02 q1bar= 1.00023365E-02
Mean allele frequencies at 2nd locus, conditioned on segregation
q2bar1= 1.00001870E-02 q2bar2= 1.00024566E-02 q2bar= 1.00023402E-02

D statistics
Mean D1= 9.89980157E-03 Mean D2= -9.99651893E-05 Mean D=
4.10622335E-04
Mean D1^2= 9.80082477E-05 Mean D2^2= 1.85920129E-07 Mean D^2=
5.01378872E-06

Mean pr1= 9.80082477E-05 Mean pr2= 9.80029290E-05 Mean pr=

9.80032055E-05
sig1d1= 101.009880 sig1d2= -1.02002251 sig1d= 4.18988657
sigd1= 0.999988914 sigd2= -1.00978576E-02 sigd= 4.14784402E-02
sig2d1= 1.00000000 sig2d2= 1.89708744E-03 sig2d= 5.11594377E-02

**Selection coefficient= 2.50000004E-02**
**Scaled selection coefficient= 50.0000000**
**Epistasis coefficient= 0.00000000**

Summary statistics for whole population

ns1= 6608 ps1= 6.60800003E-03
ns2= 993392 ps2= 0.993391991

Mean allele frequencies at 1st locus, conditioned on segregation
q1bar1= 1.64681338E-02 q1bar2= 7.16272276E-03 q1bar= 7.25588575E-03
Mean allele frequencies at 2nd locus, conditioned on segregation
q2bar1= 3.66476877E-03 q2bar2= 3.70626440E-05 q2bar= 7.33831694E-05

Mean sojourn times
tbar1= 5.21262121 tbar2= 3.42867255 tbar= 3.44046092

D statistics
Mean D1= 3.57463397E-03 Mean D2= -3.69108784E-05 Mean D=
-7.53033419E-07
Mean D1^2= 3.59722326E-05 Mean D2^2= 1.58047868E-08 Mean D^2=
3.75791160E-07

Mean pr1= 8.55122437E-07 Mean pr2= 3.50347946E-05 Mean pr=
3.58899160E-05
sig1d1= 4180.25977 sig1d2= -1.05354917 sig1d= -2.09817551E-02
sigd1= 3.86560464 sigd2= -6.23597810E-03 sigd= -1.25697901E-04
sig2d1= 42.0667610 sig2d2= 4.51116881E-04 sig2d= 1.04706613E-02
Theoretical value of unweighted sigma2d= 0.454544991

Summary statistics for sample

Sums of generation times for samples which match chosen allele frequency
tsamp1= 1357 tsamp2= 33570 tsamp= 34927
Mean allele frequencies at 1st locus, conditioned on segregation
q1bar1= 1.00001590E-02 q1bar2= 1.00004813E-02 q1bar= 1.00004692E-02
Mean allele frequencies at 2nd locus, conditioned on segregation
q2bar1= 1.00001590E-02 q2bar2= 1.00004869E-02 q2bar= 1.00004738E-02

D statistics
Mean D1= 9.90007631E-03 Mean D2= -9.99650147E-05 Mean D=
2.88561219E-04
Mean D1^2= 9.80105251E-05 Mean D2^2= 2.47427323E-07 Mean D^2=
3.81756354E-06

Mean pr1=   9.80105251E-05  Mean pr2=   9.79936594E-05  Mean pr=
9.79943143E-05
sig1d1=   101.010338     sig1d2=  -1.02011716     sig1d=   2.94467306
sigd1=   1.00000501     sigd2=  -1.00983176E-02  sigd=   2.91499309E-02
sig2d1=   1.00000000     sig2d2=   2.52493192E-03  sig2d=   3.89569886E-02

**Selection coefficient=   2.50000004E-02**
**Scaled selection coefficient=   50.0000000**
**Epistasis coefficient=   1.99999996E-02**

Summary statistics for whole population

ns1=        6573  ps1=   6.57299999E-03
ns2=      993427  ps2=   0.993426979

Mean allele frequencies at 1st locus, conditioned on segregation
q1bar1=   1.58239678E-02  q1bar2=   7.23575195E-03  q1bar=   7.31948297E-03
Mean allele frequencies at 2nd locus, conditioned on segregation
q2bar1=   3.63679929E-03  q2bar2=   3.58064499E-05  q2bar=   7.09147062E-05

Mean sojourn times
tbar1=   5.12414408     tbar2=   3.44358373     tbar=   3.45462990

D statistics
Mean D1=   3.55104613E-03 Mean D2=   -3.80188067E-05  Mean D=
-3.02712920E-06
Mean D1^2=   3.62250685E-05 Mean D2^2=   1.83402769E-08  Mean D^2=
3.71338672E-07

Mean pr1=   7.95491758E-07  Mean pr2=   3.60339909E-05  Mean pr=
3.68294823E-05
sig1d1=   4463.96338     sig1d2=  -1.05508173     sig1d=  -8.21930990E-02
sigd1=   3.98142433     sigd2=  -6.33347873E-03  sigd=  -4.98807698E-04
sig2d1=   45.5379562     sig2d2=   5.08971571E-04  sig2d=   1.00826472E-02
Theoretical value of unweighted sigma2d=   0.454544991

Summary statistics for sample

Sums of generation times for samples which match chosen allele frequency
tsamp1=        1325  tsamp2=       34179  tsamp=       35504
Mean allele frequencies at 1st locus, conditioned on segregation
q1bar1=   1.00001572E-02  q1bar2=   1.00006470E-02  q1bar=   1.00006284E-02
Mean allele frequencies at 2nd locus, conditioned on segregation
q2bar1=   1.00001572E-02  q2bar2=   1.00006517E-02  q2bar=   1.00006340E-02

D statistics
Mean D1=   9.90007538E-03 Mean D2=   -9.99638069E-05  Mean D=
2.73235055E-04

Mean D1^2= 9.80106634E-05 Mean D2^2= 2.58004548E-07 Mean D^2=
3.66736117E-06

Mean pr1= 9.80106634E-05 Mean pr2= 9.79935940E-05 Mean pr=
9.79942342E-05
sig1d1= 101.010185 sig1d2= -1.02010548 sig1d= 2.78827691
sigd1= 1.00000429 sigd2= -1.00981994E-02 sigd= 2.76017208E-02
sig2d1= 1.00000000 sig2d2= 2.63287150E-03 sig2d= 3.74242552E-02

**Selection coefficient= 2.50000004E-02**
**Scaled selection coefficient= 50.0000000**
**Epistasis coefficient= 3.99999991E-02**

Summary statistics for whole population

ns1= 6449 ps1= 6.44899998E-03
ns2= 993551 ps2= 0.993551016

Mean allele frequencies at 1st locus, conditioned on segregation
q1bar1= 1.65999383E-02 q1bar2= 7.13088037E-03 q1bar= 7.22099748E-03
Mean allele frequencies at 2nd locus, conditioned on segregation
q2bar1= 3.47225834E-03 q2bar2= 3.33634162E-05 q2bar= 6.60917940E-05

Mean sojourn times
tbar1= 5.06621170 tbar2= 3.42240095 tbar= 3.43300200

D statistics
Mean D1= 3.38773034E-03 Mean D2= -3.66580061E-05 Mean D=
-4.06798517E-06
Mean D1^2= 3.15698453E-05 Mean D2^2= 1.67481886E-08 Mean D^2=
3.17040133E-07

Mean pr1= 7.64445133E-07 Mean pr2= 3.47888927E-05 Mean pr=
3.55533411E-05
sig1d1= 4431.62012 sig1d2= -1.05372727 sig1d= -0.114419207
sigd1= 3.87467861 sigd2= -6.21510576E-03 sigd= -6.82243088E-04
sig2d1= 41.2977257 sig2d2= 4.81423456E-04 sig2d= 8.91730934E-03
Theoretical value of unweighted sigma2d= 0.454544991

Summary statistics for sample

Sums of generation times for samples which match chosen allele frequency
tsamp1= 1189 tsamp2= 33457 tsamp= 34646
Mean allele frequencies at 1st locus, conditioned on segregation
q1bar1= 1.00001488E-02 q1bar2= 1.00004496E-02 q1bar= 1.00004394E-02
Mean allele frequencies at 2nd locus, conditioned on segregation
q2bar1= 1.00001488E-02 q2bar2= 1.00004552E-02 q2bar= 1.00004449E-02

D statistics

Mean D1= 9.90007352E-03 Mean D2= -9.99652402E-05 Mean D=
2.43221468E-04
Mean D1^2= 9.80107725E-05 Mean D2^2= 2.81436115E-07 Mean D^2=
3.37324468E-06

Mean pr1= 9.80107725E-05 Mean pr2= 9.79936667E-05 Mean pr=
9.79942561E-05
sig1d1= 101.010056 sig1d2= -1.02011943 sig1d= 2.48199725
sigd1= 1.00000346 sigd2= -1.00983409E-02 sigd= 2.45697983E-02
sig2d1= 1.00000000 sig2d2= 2.87198275E-03 sig2d= 3.44228819E-02

Selection coefficient= 2.50000004E-02
Scaled selection coefficient= 50.0000000
Epistasis coefficient= 7.99999982E-02

**Selection coefficient= 2.50000004E-02**
**Scaled selection coefficient= 50.0000000**
**Epistasis coefficient= 7.99999982E-02**

Summary statistics for whole population

ns1= 6548 ps1= 6.54800003E-03
ns2= 993452 ps2= 0.993452013

Mean allele frequencies at 1st locus, conditioned on segregation
q1bar1= 1.60273649E-02 q1bar2= 7.16395583E-03 q1bar= 7.24997325E-03
Mean allele frequencies at 2nd locus, conditioned on segregation
q2bar1= 3.58739914E-03 q2bar2= 3.51567178E-05 q2bar= 6.96310599E-05

Mean sojourn times
tbar1= 5.09651804 tbar2= 3.42778826 tbar= 3.43871498

D statistics
Mean D1= 3.50264460E-03 Mean D2= -3.72022296E-05 Mean D=
-2.84876660E-06
Mean D1^2= 3.48998474E-05 Mean D2^2= 1.63566458E-08 Mean D^2=
3.54893530E-07

Mean pr1= 7.83666621E-07 Mean pr2= 3.53021605E-05 Mean pr=
3.60858285E-05
sig1d1= 4469.55957 sig1d2= -1.05382299 sig1d= -7.89441913E-02
sigd1= 3.95667529 sigd2= -6.26135478E-03 sigd= -4.74229484E-04
sig2d1= 44.5340500 sig2d2= 4.63332719E-04 sig2d= 9.83470678E-03
Theoretical value of unweighted sigma2d= 0.454544991

Summary statistics for sample

Sums of generation times for samples which match chosen allele frequency
tsamp1= 1221 tsamp2= 33500 tsamp= 34721
Mean allele frequencies at 1st locus, conditioned on segregation

q1bar1= 1.00001507E-02 q1bar2= 1.00004617E-02 q1bar= 1.00004505E-02
Mean allele frequencies at 2nd locus, conditioned on segregation
q2bar1= 1.00001507E-02 q2bar2= 1.00004673E-02 q2bar= 1.00004561E-02

D statistics
Mean D1= 9.90007445E-03 Mean D2= -9.99651529E-05 Mean D=
2.51696620E-04
Mean D1^2= 9.80108161E-05 Mean D2^2= 2.74412827E-07 Mean D^2=
3.45630201E-06

Mean pr1= 9.80108161E-05 Mean pr2= 9.79936667E-05 Mean pr=
9.79942633E-05
sig1d1= 101.010017 sig1d2= -1.02011847 sig1d= 2.56848311
sigd1= 1.00000334 sigd2= -1.00983316E-02 sigd= 2.54259408E-02
sig2d1= 1.00000000 sig2d2= 2.80031189E-03 sig2d= 3.52704525E-02

**Selection coefficient= 2.50000004E-02**
**Scaled selection coefficient= 50.0000000**
**Epistasis coefficient= 0.159999996**

Summary statistics for whole population

ns1= 6501 ps1= 6.50099991E-03
ns2= 993499 ps2= 0.993498981

Mean allele frequencies at 1st locus, conditioned on segregation
q1bar1= 1.53444186E-02 q1bar2= 7.19115557E-03 q1bar= 7.26750074E-03
Mean allele frequencies at 2nd locus, conditioned on segregation
q2bar1= 3.31858383E-03 q2bar2= 3.13683595E-05 q2bar= 6.21492654E-05

Mean sojourn times
tbar1= 4.96092892 tbar2= 3.43432045 tbar= 3.44424510

D statistics
Mean D1= 3.24140466E-03 Mean D2= -3.70935450E-05 Mean D=
-6.39455493E-06
Mean D1^2= 2.94886868E-05 Mean D2^2= 1.57036144E-08 Mean D^2=
2.91680806E-07

Mean pr1= 6.86147757E-07 Mean pr2= 3.52252428E-05 Mean pr=
3.59113874E-05
sig1d1= 4724.06201 sig1d2= -1.05303872 sig1d= -0.178064823
sigd1= 3.91313148 sigd2= -6.24987483E-03 sigd= -1.06707332E-03
sig2d1= 42.9771652 sig2d2= 4.45805723E-04 sig2d= 8.12223740E-03
Theoretical value of unweighted sigma2d= 0.454544991

Summary statistics for sample

Sums of generation times for samples which match chosen allele frequency

tsamp1= 1168 tsamp2= 33753 tsamp= 34921
Mean allele frequencies at 1st locus, conditioned on segregation
q1bar1= 1.00001469E-02 q1bar2= 1.00005316E-02 q1bar= 1.00005185E-02
Mean allele frequencies at 2nd locus, conditioned on segregation
q2bar1= 1.00001469E-02 q2bar2= 1.00005371E-02 q2bar= 1.00005241E-02

D statistics
Mean D1= 9.90007352E-03 Mean D2= -9.99646436E-05 Mean D=
2.34505875E-04
Mean D1^2= 9.80107434E-05 Mean D2^2= 2.89033409E-07 Mean D^2=
3.28782517E-06

Mean pr1= 9.80107434E-05 Mean pr2= 9.79936376E-05 Mean pr=
9.79942051E-05
sig1d1= 101.010086 sig1d2= -1.02011359 sig1d= 2.39305854
sigd1= 1.00000370 sigd2= -1.00982813E-02 sigd= 2.36893725E-02
sig2d1= 1.00000000 sig2d2= 2.94951210E-03 sig2d= 3.35512199E-02

**Selection coefficient= 2.50000004E-02**
**Scaled selection coefficient= 50.0000000**
**Epistasis coefficient= 0.319999993**

Summary statistics for whole population

ns1= 6611 ps1= 6.61099982E-03
ns2= 993389 ps2= 0.993389010

Mean allele frequencies at 1st locus, conditioned on segregation
q1bar1= 1.63054056E-02 q1bar2= 7.18021439E-03 q1bar= 7.26521062E-03
Mean allele frequencies at 2nd locus, conditioned on segregation
q2bar1= 3.13652493E-03 q2bar2= 2.94901110E-05 q2bar= 5.84308509E-05

Mean sojourn times
tbar1= 4.84677076 tbar2= 3.43065405 tbar= 3.44001603

D statistics
Mean D1= 3.06284195E-03 Mean D2= -3.73913535E-05 Mean D=
-8.51425830E-06
Mean D1^2= 2.55784253E-05 Mean D2^2= 1.76269399E-08 Mean D^2=
2.55712763E-07

Mean pr1= 6.53648328E-07 Mean pr2= 3.54726071E-05 Mean pr=
3.61262537E-05
sig1d1= 4685.76416 sig1d2= -1.05409098 sig1d= -0.235680625
sigd1= 3.78837109 sigd2= -6.27804780E-03 sigd= -1.41656131E-03
sig2d1= 39.1317825 sig2d2= 4.96916939E-04 sig2d= 7.07830815E-03
Theoretical value of unweighted sigma2d= 0.454544991

Summary statistics for sample

Sums of generation times for samples which match chosen allele frequency
tsamp1= 1044 tsamp2= 33240 tsamp= 34284
Mean allele frequencies at 1st locus, conditioned on segregation
q1bar1= 1.00001376E-02 q1bar2= 1.00003891E-02 q1bar= 1.00003816E-02
Mean allele frequencies at 2nd locus, conditioned on segregation
q2bar1= 1.00001376E-02 q2bar2= 1.00003947E-02 q2bar= 1.00003872E-02

D statistics
Mean D1= 9.90007073E-03 Mean D2= -9.99656841E-05 Mean D=
2.04550670E-04
Mean D1^2= 9.80105469E-05 Mean D2^2= 3.18443483E-07 Mean D^2=
2.99426733E-06

Mean pr1= 9.80105469E-05 Mean pr2= 9.79936958E-05 Mean pr=
9.79942051E-05
sig1d1= 101.010262 sig1d2= -1.02012360 sig1d= 2.08737516
sigd1= 1.00000429 sigd2= -1.00983838E-02 sigd= 2.06633490E-02
sig2d1= 1.00000000 sig2d2= 3.24963243E-03 sig2d= 3.05555556E-02

Selection coefficient= 2.50000004E-02
Scaled selection coefficient= 50.0000000
Epistasis coefficient= 0.639999986

Summary statistics for whole population

ns1= 6451 ps1= 6.45099999E-03
ns2= 993549 ps2= 0.993548989

Mean allele frequencies at 1st locus, conditioned on segregation
q1bar1= 1.52360871E-02 q1bar2= 7.23737525E-03 q1bar= 7.30269821E-03
Mean allele frequencies at 2nd locus, conditioned on segregation
q2bar1= 2.57623359E-03 q2bar2= 2.12134310E-05 q2bar= 4.20803735E-05

Mean sojourn times
tbar1= 4.36490488 tbar2= 3.44195199 tbar= 3.44790602

D statistics
Mean D1= 2.52082082E-03 Mean D2= -3.78193363E-05 Mean D=
-1.69236991E-05
Mean D1^2= 1.59701995E-05 Mean D2^2= 1.69285954E-08 Mean D^2=
1.47214109E-07

Mean pr1= 4.33190024E-07 Mean pr2= 3.59411169E-05 Mean pr=
3.63743056E-05
sig1d1= 5819.20312 sig1d2= -1.05225825 sig1d= -0.465265214
sigd1= 3.83003521 sigd2= -6.30838377E-03 sigd= -2.80606654E-03
sig2d1= 36.8664970 sig2d2= 4.71009174E-04 sig2d= 4.04720055E-03
Theoretical value of unweighted sigma2d= 0.454544991

Summary statistics for sample

Sums of generation times for samples which match chosen allele frequency
tsamp1= 858 tsamp2= 34192 tsamp= 35050
Mean allele frequencies at 1st locus, conditioned on segregation
q1bar1= 1.00001181E-02 q1bar2= 1.00006498E-02 q1bar= 1.00006377E-02
Mean allele frequencies at 2nd locus, conditioned on segregation
q2bar1= 1.00001181E-02 q2bar2= 1.00006554E-02 q2bar= 1.00006424E-02

D statistics
Mean D1= 9.90006607E-03 Mean D2= -9.99637778E-05 Mean D=
1.44830105E-04
Mean D1^2= 9.80101468E-05 Mean D2^2= 3.98585257E-07 Mean D^2=
2.40897839E-06

Mean pr1= 9.80101468E-05 Mean pr2= 9.79935940E-05 Mean pr=
9.79939941E-05
sig1d1= 101.010620 sig1d2= -1.02010524 sig1d= 1.47794878
sigd1= 1.00000596 sigd2= -1.00981966E-02 sigd= 1.46304984E-02
sig2d1= 1.00000000 sig2d2= 4.06746240E-03 sig2d= 2.45829187E-02

**Selection coefficient= 2.50000004E-02**
**Scaled selection coefficient= 50.0000000**
**Epistasis coefficient= 1.27999997**

Summary statistics for whole population

ns1= 6576 ps1= 6.57599978E-03
ns2= 993424 ps2= 0.993423998

Mean allele frequencies at 1st locus, conditioned on segregation
q1bar1= 1.43991429E-02 q1bar2= 7.15646334E-03 q1bar= 7.20906677E-03
Mean allele frequencies at 2nd locus, conditioned on segregation
q2bar1= 2.03886465E-03 q2bar2= 1.49168955E-05 q2bar= 2.96171074E-05

Mean sojourn times
tbar1= 3.79273105 tbar2= 3.43160820 tbar= 3.43398309

D statistics
Mean D1= 1.99643336E-03 Mean D2= -3.73208859E-05 Mean D=
-2.25497388E-05
Mean D1^2= 9.61851129E-06 Mean D2^2= 1.78582980E-08 Mean D^2=
8.75877930E-08

Mean pr1= 2.95095077E-07 Mean pr2= 3.54745462E-05 Mean pr=
3.57696408E-05
sig1d1= 6765.39014 sig1d2= -1.05204690 sig1d= -0.630415559
sigd1= 3.67513943 sigd2= -6.26604492E-03 sigd= -3.77037236E-03

sig2d1= 32.5946198 sig2d2= 5.03411575E-04 sig2d= 2.44866288E-03
Theoretical value of unweighted sigma2d= 0.454544991

Summary statistics for sample

Sums of generation times for samples which match chosen allele frequency
tsamp1= 581 tsamp2= 33450 tsamp= 34031
Mean allele frequencies at 1st locus, conditioned on segregation
q1bar1= 1.00000650E-02 q1bar2= 1.00004477E-02 q1bar= 1.00004412E-02
Mean allele frequencies at 2nd locus, conditioned on segregation
q2bar1= 1.00000650E-02 q2bar2= 1.00004533E-02 q2bar= 1.00004468E-02

D statistics
Mean D1= 9.90005396E-03 Mean D2= -9.99652548E-05 Mean D=
7.07617583E-05
Mean D1^2= 9.80094483E-05 Mean D2^2= 5.75830427E-07 Mean D^2=
1.68311374E-06

Mean pr1= 9.80094483E-05 Mean pr2= 9.79936740E-05 Mean pr=
9.79939359E-05
sig1d1= 101.011223 sig1d2= -1.02011943 sig1d= 0.722103417
sigd1= 1.00000823 sigd2= -1.00983409E-02 sigd= 7.14823790E-03
sig2d1= 1.00000000 sig2d2= 5.87619981E-03 sig2d= 1.71756931E-02
