## Supplementary File S5 for "A gene-based model of fitness and its implications for genetic variation: Linkage disequilibrium"

### Simulation Results for 2-locus LD with no Recombination

### Population statistics with Good's weighting towards low derived allele frequencies

**Size of haploid population= 1000**
**Number of replications= 1000000**
**Selection coefficient= 0.00000000**
**Scaled selection coefficient= 0.00000000**
**Epistasis coefficient= 0.00000000**

**Allele frequency weight= 10.0000000**

Summary statistics for whole population

ns1= 134844 ps1= 0.134844005
ns2= 865156 ps2= 0.865155995

Mean allele frequencies at 1st locus, conditioned on segregation
q1bar1= 0.464014411 q1bar2= 0.131491452 q1bar= 0.204718426
Mean allele frequencies at 2nd locus, conditioned on segregation
q2bar1= 9.56748724E-02 q2bar2= 2.70192418E-02 q2bar=
4.21383306E-02

Mean sojourn times
tbar1= 12.9488592 tbar2= 7.14650059 tbar= 7.92891407

D statistics
Mean D1= 1.19403121E-03 Mean D2= -1.76531263E-04 Mean D=
1.25288905E-04
Mean D1^2= 1.17407426E-05 Mean D2^2= 1.12109845E-07 Mean D^2=
2.67292421E-06

Mean pr1= 2.37475015E-05 Mean pr2= 1.13226262E-04 Mean pr=
1.36973773E-04
sig1d1= 50.2802887 sig1d2= -1.55910170 sig1d= 0.914692640
sigd1= 0.245022923 sigd2= -1.65900625E-02 sigd= 1.07051777E-02
sig2d1= 0.494399071 sig2d2= 9.90139903E-04 sig2d=
1.95141323E-02
Theoretical value of unweighted sigma2d= 0.454544991

**Allele frequency weight= 100.000000**

Summary statistics for whole population

ns1= 134649 ps1= 0.134648994
ns2= 865351 ps2= 0.865351021

Mean allele frequencies at 1st locus, conditioned on segregation
q1bar1= 0.463013232 q1bar2= 0.130399674 q1bar= 0.204821959
Mean allele frequencies at 2nd locus, conditioned on segregation
q2bar1= 9.94325355E-02 q2bar2= 2.86609046E-02 q2bar= 4.44960557E-02

Mean sojourn times
tbar1= 13.2036333 tbar2= 7.12759352 tbar= 7.94572592

D statistics
Mean D1= 1.91951549E-05 Mean D2= -1.78142955E-06 Mean D=
2.91208084E-06
Mean D1^2= 2.15825775E-08 Mean D2^2= 1.28844236E-11 Mean D^2=
4.83910245E-09

Mean pr1= 4.68440078E-08 Mean pr2= 1.35315452E-06 Mean pr=
1.39999850E-06
sig1d1= 409.767548 sig1d2= -1.31650126 sig1d= 2.08006001
sigd1= 8.86879489E-02 sigd2= -1.53142225E-03 sigd= 2.46115890E-03
sig2d1= 0.460732937 sig2d2= 9.52176833E-06 sig2d= 3.45650548E-03
Theoretical value of unweighted sigma2d= 0.454544991

**Selection coefficient= 4.99999989E-03**
**Scaled selection coefficient= 10.0000000**
**Epistasis coefficient= 0.00000000**

**Allele frequency weight= 10.0000000**

Summary statistics for whole population

ns1= 23664 ps1= 2.36639995E-02
ns2= 976336 ps2= 0.976336002

Mean allele frequencies at 1st locus, conditioned on segregation
q1bar1= 9.95222628E-02 q1bar2= 2.89978720E-02 q1bar= 3.16699371E-02
Mean allele frequencies at 2nd locus, conditioned on segregation
q2bar1= 1.20215379E-02 q2bar2= 4.73415654E-04 q2bar= 9.10957227E-04

Mean sojourn times
tbar1= 7.99277401 tbar2= 4.91930771 tbar= 4.99203777

D statistics
Mean D1= 3.24830483E-03 Mean D2= -1.07170337E-04 Mean D=
1.99636997E-05
Mean D1^2= 3.02259868E-05 Mean D2^2= 5.51295649E-08 Mean D^2=

1.19825904E-06

Mean pr1= 6.90550405E-06 Mean pr2= 9.29223170E-05 Mean pr=
9.98278192E-05
sig1d1= 470.393585 sig1d2= -1.15333259 sig1d= 0.199981332
sigd1= 1.23611557 sigd2= -1.11176902E-02 sigd= 1.99809088E-03
sig2d1= 4.37708616 sig2d2= 5.93286590E-04 sig2d= 1.20032579E-02
Theoretical value of unweighted sigma2d= 0.454544991

**Allele frequency weight= 100.000000**

Summary statistics for whole population

ns1= 23695 ps1= 2.36949995E-02
ns2= 976305 ps2= 0.976305008

Mean allele frequencies at 1st locus, conditioned on segregation
q1bar1= 0.101071864 q1bar2= 2.94975266E-02 q1bar= 3.23193520E-02
Mean allele frequencies at 2nd locus, conditioned on segregation
q2bar1= 1.26309050E-02 q2bar2= 5.18413552E-04 q2bar= 9.95950075E-04

Mean sojourn times
tbar1= 8.40413570 tbar2= 4.96961308 tbar= 5.05099392

D statistics
Mean D1= 1.65037360E-04 Mean D2= -3.00740021E-06 Mean D=
3.61778325E-06
Mean D1^2= 1.86129071E-07 Mean D2^2= 2.18649526E-11 Mean D^2=
7.35916217E-09

Mean pr1= 6.64270416E-08 Mean pr2= 2.83072177E-06 Mean pr=
2.89714876E-06
sig1d1= 2484.49048 sig1d2= -1.06241465 sig1d= 1.24873924
sigd1= 0.640338778 sigd2= -1.78748602E-03 sigd= 2.12548045E-03
sig2d1= 2.80200744 sig2d2= 7.72416206E-06 sig2d= 2.54013948E-03
Theoretical value of unweighted sigma2d= 0.454544991

**Selection coefficient= 4.99999989E-03**
**Scaled selection coefficient= 10.0000000**
**Epistasis coefficient= 1.99999996E-02**

**Allele frequency weight= 10.0000000**

Summary statistics for whole population

ns1= 23652 ps1= 2.36520004E-02
ns2= 976348 ps2= 0.976347983

Mean allele frequencies at 1st locus, conditioned on segregation

q1bar1= 0.104158565 q1bar2= 3.00233699E-02 q1bar= 3.28975581E-02
Mean allele frequencies at 2nd locus, conditioned on segregation
q2bar1= 1.30397351E-02 q2bar2= 5.25935786E-04 q2bar= 1.01109094E-03

Mean sojourn times
tbar1= 8.29308319 tbar2= 4.98099327 tbar= 5.05933094

D statistics
Mean D1= 3.24212224E-03 Mean D2= -1.11958514E-04 Mean D=
1.80776988E-05
Mean D1^2= 2.97450315E-05 Mean D2^2= 5.95286451E-08 Mean D^2=
1.21042228E-06

Mean pr1= 7.26174176E-06 Mean pr2= 9.65604777E-05 Mean pr=
1.03822225E-04
sig1d1= 446.466187 sig1d2= -1.15946519 sig1d= 0.174121663
sigd1= 1.20312011 sigd2= -1.13935061E-02 sigd= 1.77418126E-03
sig2d1= 4.09612894 sig2d2= 6.16490783E-04 sig2d= 1.16586043E-02
Theoretical value of unweighted sigma2d= 0.454544991

**Allele frequency weight= 100.000000**

Summary statistics for whole population

ns1= 23888 ps1= 2.38879994E-02
ns2= 976112 ps2= 0.976112008

Mean allele frequencies at 1st locus, conditioned on segregation
q1bar1= 0.101535350 q1bar2= 2.94477604E-02 q1bar= 3.22479457E-02
Mean allele frequencies at 2nd locus, conditioned on segregation
q2bar1= 1.20246895E-02 q2bar2= 4.85966593E-04 q2bar= 9.34179232E-04

Mean sojourn times
tbar1= 8.19491005 tbar2= 4.96240187 tbar= 5.03961992

D statistics
Mean D1= 1.59555682E-04 Mean D2= -3.02199578E-06 Mean D=
3.29320392E-06
Mean D1^2= 1.79578919E-07 Mean D2^2= 2.19961688E-11 Mean D^2=
6.99674096E-09

Mean pr1= 6.27796197E-08 Mean pr2= 2.84631983E-06 Mean pr=
2.90909929E-06
sig1d1= 2541.52026 sig1d2= -1.06172037 sig1d= 1.13203561
sigd1= 0.636799872 sigd2= -1.79123261E-03 sigd= 1.93080911E-03
sig2d1= 2.86046529 sig2d2= 7.72793283E-06 sig2d= 2.40512285E-03
Theoretical value of unweighted sigma2d= 0.454544991

**Selection coefficient=    4.99999989E-03**
**Scaled selection coefficient=    10.0000000**
**Epistasis coefficient=    3.99999991E-02**

**Allele frequency weight=    10.0000000**

Summary statistics for whole population

ns1=        24088  ps1=    2.40880009E-02
ns2=       975912  ps2=    0.975911975

Mean allele frequencies at 1st locus, conditioned on segregation
q1bar1=    9.82538909E-02  q1bar2=    2.95222159E-02  q1bar=    3.22005861E-02
Mean allele frequencies at 2nd locus, conditioned on segregation
q2bar1=    1.18926791E-02  q2bar2=    4.82232106E-04  q2bar=    9.26880457E-04

Mean sojourn times
tbar1=    8.16265392      tbar2=    4.96872759      tbar=    5.04566288

D statistics
Mean D1=    3.37105291E-03 Mean D2=   -1.10332767E-04  Mean D=
2.53316684E-05
Mean D1^2=    3.15007528E-05 Mean D2^2=    5.78219854E-08  Mean D^2=
1.28310626E-06

Mean pr1=    7.14427870E-06  Mean pr2=    9.53488197E-05  Mean pr=
1.02493097E-04
sig1d1=    471.853485      sig1d2=   -1.15714872      sig1d=   0.247154877
sigd1=    1.26120698      sigd2=   -1.12991780E-02  sigd=    2.50216830E-03
sig2d1=    4.40922785      sig2d2=    6.06425805E-04  sig2d=    1.25189526E-02
Theoretical value of unweighted sigma2d=   0.454544991

**Allele frequency weight=    100.000000**

Summary statistics for whole population

ns1=        23659  ps1=    2.36590002E-02
ns2=       976341  ps2=    0.976341009

Mean allele frequencies at 1st locus, conditioned on segregation
q1bar1=   0.101617515      q1bar2=    2.95971148E-02  q1bar=    3.24469618E-02
Mean allele frequencies at 2nd locus, conditioned on segregation
q2bar1=    1.19358953E-02  q2bar2=    4.91762825E-04  q2bar=    9.44607542E-04

Mean sojourn times
tbar1=    8.43957901      tbar2=    4.96381187      tbar=    5.04604483

D statistics
Mean D1=    1.59364252E-04 Mean D2=   -3.02291664E-06  Mean D=
3.40274391E-06

Mean D1^2= 1.78621931E-07 Mean D2^2= 2.20339910E-11 Mean D^2=
7.08923231E-09

Mean pr1= 6.45812150E-08 Mean pr2= 2.84503426E-06 Mean pr=
2.90961543E-06
sig1d1= 2467.65649 sig1d2= -1.06252384 sig1d= 1.16948235
sigd1= 0.627101481 sigd2= -1.79218326E-03 sigd= 1.99485570E-03
sig2d1= 2.76584959 sig2d2= 7.74471937E-06 sig2d= 2.43648421E-03
Theoretical value of unweighted sigma2d= 0.454544991

**Selection coefficient= 4.99999989E-03**
**Scaled selection coefficient= 10.0000000**
**Epistasis coefficient= 7.99999982E-02**

**Allele frequency weight= 10.0000000**

Summary statistics for whole population

ns1= 23711 ps1= 2.37109996E-02
ns2= 976289 ps2= 0.976288974

Mean allele frequencies at 1st locus, conditioned on segregation
q1bar1= 0.101355776 q1bar2= 2.98216641E-02 q1bar= 3.25034633E-02
Mean allele frequencies at 2nd locus, conditioned on segregation
q2bar1= 1.10096922E-02 q2bar2= 4.28827916E-04 q2bar= 8.25502502E-04

Mean sojourn times
tbar1= 7.97933435 tbar2= 4.97542858 tbar= 5.04665422

D statistics
Mean D1= 3.11333244E-03 Mean D2= -1.11139736E-04 Mean D=
9.74504474E-06
Mean D1^2= 2.81649136E-05 Mean D2^2= 5.86194737E-08 Mean D^2=
1.11231850E-06

Mean pr1= 6.62739831E-06 Mean pr2= 9.60838515E-05 Mean pr=
1.02711252E-04
sig1d1= 469.766907 sig1d2= -1.15669525 sig1d= 9.48780626E-02
sigd1= 1.20935547 sigd2= -1.13382014E-02 sigd= 9.61556565E-04
sig2d1= 4.24976921 sig2d2= 6.10086659E-04 sig2d= 1.08295679E-02
Theoretical value of unweighted sigma2d= 0.454544991

**Allele frequency weight= 100.000000**

Summary statistics for whole population

ns1= 23965 ps1= 2.39649992E-02
ns2= 976035 ps2= 0.976034999

Mean allele frequencies at 1st locus, conditioned on segregation

q1bar1= 9.95988399E-02 q1bar2= 2.94967704E-02 q1bar= 3.22322585E-02
Mean allele frequencies at 2nd locus, conditioned on segregation
q2bar1= 1.27304085E-02 q2bar2= 5.16931119E-04 q2bar= 9.93519323E-04

Mean sojourn times
tbar1= 8.20580006 tbar2= 4.96184063 tbar= 5.03958178

D statistics
Mean D1= 1.51745990E-04 Mean D2= -3.02416834E-06 Mean D=
3.01519367E-06
Mean D1^2= 1.67619689E-07 Mean D2^2= 2.20172058E-11 Mean D^2=
6.56192833E-09

Mean pr1= 6.21745784E-08 Mean pr2= 2.84792191E-06 Mean pr=
2.91009633E-06
sig1d1= 2440.64355 sig1d2= -1.06188595 sig1d= 1.03611469
sigd1= 0.608570337 sigd2= -1.79201609E-03 sigd= 1.76750857E-03
sig2d1= 2.69595218 sig2d2= 7.73097236E-06 sig2d= 2.25488353E-03
Theoretical value of unweighted sigma2d= 0.454544991

**Selection coefficient= 4.99999989E-03**
**Scaled selection coefficient= 10.0000000**
**Epistasis coefficient= 0.159999996**

**Allele frequency weight= 10.0000000**

Summary statistics for whole population

ns1= 23981 ps1= 2.39809994E-02
ns2= 976019 ps2= 0.976019025

Mean allele frequencies at 1st locus, conditioned on segregation
q1bar1= 0.100944795 q1bar2= 2.95819137E-02 q1bar= 3.22856866E-02
Mean allele frequencies at 2nd locus, conditioned on segregation
q2bar1= 1.09279472E-02 q2bar2= 4.30339307E-04 q2bar= 8.28069518E-04

Mean sojourn times
tbar1= 7.94345522 tbar2= 4.95617294 tbar= 5.02781105

D statistics
Mean D1= 3.04534938E-03 Mean D2= -1.09376757E-04 Mean D=
1.01484347E-05
Mean D1^2= 2.66726402E-05 Mean D2^2= 5.70312046E-08 Mean D^2=
1.06543439E-06

Mean pr1= 6.52314066E-06 Mean pr2= 9.46472646E-05 Mean pr=
1.01170408E-04
sig1d1= 466.853241 sig1d2= -1.15562510 sig1d= 0.100310311
sigd1= 1.19236374 sigd2= -1.12427101E-02 sigd= 1.00895623E-03
sig2d1= 4.08892632 sig2d2= 6.02565822E-04 sig2d= 1.05310874E-02

Theoretical value of unweighted sigma2d= 0.454544991

**Allele frequency weight= 100.000000**

Summary statistics for whole population

ns1= 23641 ps1= 2.36409996E-02
ns2= 976359 ps2= 0.976359010

Mean allele frequencies at 1st locus, conditioned on segregation
q1bar1= 0.102974437 q1bar2= 2.95178276E-02 q1bar= 3.23398151E-02
Mean allele frequencies at 2nd locus, conditioned on segregation
q2bar1= 1.10472981E-02 q2bar2= 4.41360899E-04 q2bar= 8.48810188E-04

Mean sojourn times
tbar1= 8.20519447 tbar2= 4.97287560 tbar= 5.04929113

D statistics
Mean D1= 1.58646988E-04 Mean D2= -3.02014405E-06 Mean D=
3.19063429E-06
Mean D1^2= 1.78974659E-07 Mean D2^2= 2.19956432E-11 Mean D^2=
6.89683377E-09

Mean pr1= 6.19916278E-08 Mean pr2= 2.84566727E-06 Mean pr=
2.90765888E-06
sig1d1= 2559.16797 sig1d2= -1.06131315 sig1d= 1.09732068
sigd1= 0.637184680 sigd2= -1.79034041E-03 sigd= 1.87113578E-03
sig2d1= 2.88707781 sig2d2= 7.72952080E-06 sig2d= 2.37195427E-03
Theoretical value of unweighted sigma2d= 0.454544991

**Selection coefficient= 4.99999989E-03**
**Scaled selection coefficient= 10.0000000**
**Epistasis coefficient= 0.319999993**

**Allele frequency weight= 10.0000000**

Summary statistics for whole population

ns1= 23794 ps1= 2.37939991E-02
ns2= 976206 ps2= 0.976206005

Mean allele frequencies at 1st locus, conditioned on segregation
q1bar1= 9.97561142E-02 q1bar2= 2.92188264E-02 q1bar= 3.18362825E-02
Mean allele frequencies at 2nd locus, conditioned on segregation
q2bar1= 1.10843917E-02 q2bar2= 4.27163759E-04 q2bar= 8.22625705E-04

Mean sojourn times
tbar1= 7.81465912 tbar2= 4.94257975 tbar= 5.01091814

D statistics

Mean D1= 3.01068812E-03 Mean D2= -1.08951041E-04 Mean D=
6.81056781E-06
Mean D1^2= 2.66156549E-05 Mean D2^2= 5.68105456E-08 Mean D^2=
1.04233936E-06

Mean pr1= 6.29483293E-06 Mean pr2= 9.44529165E-05 Mean pr=
1.00747755E-04
sig1d1= 478.279266 sig1d2= -1.15349579 sig1d= 6.76001906E-02
sigd1= 1.19997907 sigd2= -1.12104667E-02 sigd= 6.78524666E-04
sig2d1= 4.22817516 sig2d2= 6.01469481E-04 sig2d= 1.03460308E-02
Theoretical value of unweighted sigma2d= 0.454544991

**Allele frequency weight= 100.000000**

Summary statistics for whole population

ns1= 23793 ps1= 2.37930007E-02
ns2= 976207 ps2= 0.976207018

Mean allele frequencies at 1st locus, conditioned on segregation
q1bar1= 9.84127373E-02 q1bar2= 2.98482049E-02 q1bar= 3.22690159E-02
Mean allele frequencies at 2nd locus, conditioned on segregation
q2bar1= 9.16109513E-03 q2bar2= 3.35289136E-04 q2bar= 6.46902190E-04

Mean sojourn times
tbar1= 7.46160650 tbar2= 4.96898603 tbar= 5.02829313

D statistics
Mean D1= 1.55864895E-04 Mean D2= -3.01464320E-06 Mean D=
2.59491867E-06
Mean D1^2= 1.72832742E-07 Mean D2^2= 2.19738880E-11 Mean D^2=
6.12340534E-09

Mean pr1= 5.58657725E-08 Mean pr2= 2.84984503E-06 Mean pr=
2.90571074E-06
sig1d1= 2789.98901 sig1d2= -1.05782712 sig1d= 0.893040955
sigd1= 0.659440219 sigd2= -1.78576913E-03 sigd= 1.52229064E-03
sig2d1= 3.09371424 sig2d2= 7.71055511E-06 sig2d= 2.10736925E-03
Theoretical value of unweighted sigma2d= 0.454544991

**Selection coefficient= 4.99999989E-03**
**Scaled selection coefficient= 10.0000000**
**Epistasis coefficient= 0.639999986**

**Allele frequency weight= 10.0000000**

Summary statistics for whole population

ns1= 23791 ps1= 2.37910002E-02
ns2= 976209 ps2= 0.976208985

Mean allele frequencies at 1st locus, conditioned on segregation
q1bar1= 9.79156271E-02 q1bar2= 2.96486393E-02 q1bar= 3.20755243E-02
Mean allele frequencies at 2nd locus, conditioned on segregation
q2bar1= 9.02181957E-03 q2bar2= 3.32546711E-04 q2bar= 6.41449413E-04

Mean sojourn times
tbar1= 7.48320770 tbar2= 4.94765568 tbar= 5.00797892

D statistics
Mean D1= 2.79971259E-03 Mean D2= -1.09477252E-04 Mean D=
-6.05593641E-06
Mean D1^2= 2.31533977E-05 Mean D2^2= 5.72193883E-08 Mean D^2=
8.78285562E-07

Mean pr1= 5.50056257E-06 Mean pr2= 9.49538226E-05 Mean pr=
1.00454381E-04
sig1d1= 508.986603 sig1d2= -1.15295255 sig1d= -6.02854379E-02
sigd1= 1.19374049 sigd2= -1.12348599E-02 sigd= -6.04222470E-04
sig2d1= 4.20927811 sig2d2= 6.02602260E-04 sig2d= 8.74312874E-03
Theoretical value of unweighted sigma2d= 0.454544991

**Allele frequency weight= 100.000000**

Summary statistics for whole population

ns1= 23984 ps1= 2.39840001E-02
ns2= 976016 ps2= 0.976015985

Mean allele frequencies at 1st locus, conditioned on segregation
q1bar1= 9.88415331E-02 q1bar2= 2.96751130E-02 q1bar= 3.20875570E-02
Mean allele frequencies at 2nd locus, conditioned on segregation
q2bar1= 8.30864161E-03 q2bar2= 3.00268643E-04 q2bar= 5.79591258E-04

Mean sojourn times
tbar1= 7.32563353 tbar2= 4.98115730 tbar= 5.03738689

D statistics
Mean D1= 1.58234441E-04 Mean D2= -3.01975592E-06 Mean D=
2.60459660E-06
Mean D1^2= 1.74201830E-07 Mean D2^2= 2.20108532E-11 Mean D^2=
6.09719342E-09

Mean pr1= 5.66719471E-08 Mean pr2= 2.85588590E-06 Mean pr=
2.91255787E-06
sig1d1= 2792.11230 sig1d2= -1.05737972 sig1d= 0.894264340
sigd1= 0.664686620 sigd2= -1.78690488E-03 sigd= 1.52617099E-03
sig2d1= 3.07386351 sig2d2= 7.70718907E-06 sig2d= 2.09341547E-03
Theoretical value of unweighted sigma2d= 0.454544991

**Selection coefficient= 4.99999989E-03**
**Scaled selection coefficient= 10.0000000**
**Epistasis coefficient= 1.27999997**

**Allele frequency weight= 10.0000000**

Summary statistics for whole population

ns1= 23962 ps1= 2.39620004E-02
ns2= 976038 ps2= 0.976037979

Mean allele frequencies at 1st locus, conditioned on segregation
q1bar1= 9.46139693E-02 q1bar2= 2.94000581E-02 q1bar= 3.14721353E-02
Mean allele frequencies at 2nd locus, conditioned on segregation
q2bar1= 6.84449123E-03 q2bar2= 2.24610572E-04 q2bar= 4.34947811E-04

Mean sojourn times
tbar1= 6.63108253 tbar2= 4.96080494 tbar= 5.00082779

D statistics
Mean D1= 2.39733979E-03 Mean D2= -1.10846879E-04 Mean D=
-3.11529111E-05
Mean D1^2= 1.82950826E-05 Mean D2^2= 5.86722102E-08 Mean D^2=
6.38107508E-07

Mean pr1= 3.96703217E-06 Mean pr2= 9.65294894E-05 Mean pr=
1.00496516E-04
sig1d1= 604.315674 sig1d2= -1.14832139 sig1d= -0.309989959
sigd1= 1.20364034 sigd2= -1.12821916E-02 sigd= -3.10758571E-03
sig2d1= 4.61178064 sig2d2= 6.07816444E-04 sig2d= 6.34954870E-03
Theoretical value of unweighted sigma2d= 0.454544991

**Allele frequency weight= 100.000000**

Summary statistics for whole population

ns1= 23711 ps1= 2.37109996E-02
ns2= 976289 ps2= 0.976288974

Mean allele frequencies at 1st locus, conditioned on segregation
q1bar1= 9.44105759E-02 q1bar2= 2.94521078E-02 q1bar= 3.15152481E-02
Mean allele frequencies at 2nd locus, conditioned on segregation
q2bar1= 6.61525456E-03 q2bar2= 2.16998873E-04 q2bar= 4.20213561E-04

Mean sojourn times
tbar1= 6.68525171 tbar2= 4.94969845 tbar= 4.99084997

D statistics
Mean D1= 1.53136876E-04 Mean D2= -3.01824002E-06 Mean D=

1.94139034E-06
Mean D1^2= 1.68993637E-07 Mean D2^2= 2.19843605E-11 Mean D^2=
5.38867972E-09

Mean pr1= 4.89813843E-08 Mean pr2= 2.86370323E-06 Mean pr=
2.91268475E-06
sig1d1= 3126.43018 sig1d2= -1.05396402 sig1d= 0.666529536
sigd1= 0.691933334 sigd2= -1.78356841E-03 sigd= 1.13753858E-03
sig2d1= 3.45016050 sig2d2= 7.67689926E-06 sig2d= 1.85007311E-03
Theoretical value of unweighted sigma2d= 0.454544991

**Selection coefficient= 2.50000004E-02**
**Scaled selection coefficient= 50.0000000**
**Epistasis coefficient= 0.00000000**

**Allele frequency weight= 10.0000000**

Summary statistics for whole population

ns1= 6542 ps1= 6.54199999E-03
ns2= 993458 ps2= 0.993457973

Mean allele frequencies at 1st locus, conditioned on segregation
q1bar1= 1.55718485E-02 q1bar2= 7.15696812E-03 q1bar= 7.24124350E-03
Mean allele frequencies at 2nd locus, conditioned on segregation
q2bar1= 3.82935279E-03 q2bar2= 3.87393375E-05 q2bar= 7.67027232E-05

Mean sojourn times
tbar1= 5.28431654 tbar2= 3.43974590 tbar= 3.45181298

D statistics
Mean D1= 2.70975218E-03 Mean D2= -2.52264126E-05 Mean D=
2.16447165E-06
Mean D1^2= 1.71401534E-05 Mean D2^2= 4.25073399E-09 Mean D^2=
1.75867243E-07

Mean pr1= 5.25691974E-07 Mean pr2= 2.41172420E-05 Mean pr=
2.46429336E-05
sig1d1= 5154.63867 sig1d2= -1.04599082 sig1d= 8.78333598E-02
sigd1= 3.73735094 sigd2= -5.13678836E-03 sigd= 4.36019298E-04
sig2d1= 32.6049347 sig2d2= 1.76252899E-04 sig2d= 7.13661965E-03
Theoretical value of unweighted sigma2d= 0.454544991

**Allele frequency weight= 100.000000**

Summary statistics for whole population

ns1= 6569 ps1= 6.56899996E-03
ns2= 993431 ps2= 0.993430972

Mean allele frequencies at 1st locus, conditioned on segregation
q1bar1= 1.63250193E-02 q1bar2= 7.11567886E-03 q1bar= 7.20944349E-03
Mean allele frequencies at 2nd locus, conditioned on segregation
q2bar1= 3.93934874E-03 q2bar2= 4.05213941E-05 q2bar= 8.02176583E-05

Mean sojourn times
tbar1= 5.32288027 tbar2= 3.42179990 tbar= 3.43428802

D statistics
Mean D1= 5.14567073E-04 Mean D2= -3.43534498E-06 Mean D=
1.83866564E-06
Mean D1^2= 6.05013895E-07 Mean D2^2= 2.45253401E-11 Mean D^2=
6.18418916E-09

Mean pr1= 4.14593195E-08 Mean pr2= 3.34873948E-06 Mean pr=
3.39019903E-06
sig1d1= 12411.3730 sig1d2= -1.02586210 sig1d= 0.542347431
sigd1= 2.52714920 sigd2= -1.87728263E-03 sigd= 9.98596777E-04
sig2d1= 14.5929527 sig2d2= 7.32375293E-06 sig2d= 1.82413752E-03
Theoretical value of unweighted sigma2d= 0.454544991

**Selection coefficient= 2.50000004E-02**
**Scaled selection coefficient= 50.0000000**
**Epistasis coefficient= 1.99999996E-02**

**Allele frequency weight= 10.0000000**

Summary statistics for whole population

ns1= 6563 ps1= 6.56299992E-03
ns2= 993437 ps2= 0.993436992

Mean allele frequencies at 1st locus, conditioned on segregation
q1bar1= 1.54879736E-02 q1bar2= 7.18807662E-03 q1bar= 7.27025792E-03
Mean allele frequencies at 2nd locus, conditioned on segregation
q2bar1= 3.74656636E-03 q2bar2= 3.74676601E-05 q2bar= 7.41933545E-05

Mean sojourn times
tbar1= 5.19762325 tbar2= 3.43357968 tbar= 3.44515705

D statistics
Mean D1= 2.70359125E-03 Mean D2= -2.53159615E-05 Mean D=
1.70413864E-06
Mean D1^2= 1.69618452E-05 Mean D2^2= 4.36392344E-09 Mean D^2=
1.72267335E-07

Mean pr1= 4.98448117E-07 Mean pr2= 2.41911257E-05 Mean pr=
2.46895725E-05
sig1d1= 5424.01758 sig1d2= -1.04649782 sig1d= 6.90226033E-02
sigd1= 3.82940292 sigd2= -5.14714466E-03 sigd= 3.42963700E-04

sig2d1= 34.0293083 sig2d2= 1.80393574E-04 sig2d= 6.97733182E-03
Theoretical value of unweighted sigma2d= 0.454544991

**Allele frequency weight= 100.000000**

Summary statistics for whole population

ns1= 6588 ps1= 6.58799987E-03
ns2= 993412 ps2= 0.993412018

Mean allele frequencies at 1st locus, conditioned on segregation
q1bar1= 1.59108657E-02 q1bar2= 7.18599791E-03 q1bar= 7.27480836E-03
Mean allele frequencies at 2nd locus, conditioned on segregation
q2bar1= 3.84972780E-03 q2bar2= 3.95896859E-05 q2bar= 7.83734067E-05

Mean sojourn times
tbar1= 5.31967211 tbar2= 3.43052220 tbar= 3.44296789

D statistics
Mean D1= 5.17541601E-04 Mean D2= -3.43330089E-06 Mean D=
1.86970681E-06
Mean D1^2= 5.97282963E-07 Mean D2^2= 2.44763567E-11 Mean D^2=
6.10397510E-09

Mean pr1= 4.22198774E-08 Mean pr2= 3.34656329E-06 Mean pr=
3.38878317E-06
sig1d1= 12258.2451 sig1d2= -1.02591836 sig1d= 0.551733971
sigd1= 2.51875997 sigd2= -1.87677564E-03 sigd= 1.01566769E-03
sig2d1= 14.1469612 sig2d2= 7.31387854E-06 sig2d= 1.80122920E-03
Theoretical value of unweighted sigma2d= 0.454544991

**Scaled selection coefficient= 50.0000000**
**Epistasis coefficient= 3.99999991E-02**

**Allele frequency weight= 10.0000000**

Summary statistics for whole population

ns1= 6633 ps1= 6.63299998E-03
ns2= 993367 ps2= 0.993367016

Mean allele frequencies at 1st locus, conditioned on segregation
q1bar1= 1.66430231E-02 q1bar2= 7.21562468E-03 q1bar= 7.30589591E-03
Mean allele frequencies at 2nd locus, conditioned on segregation
q2bar1= 3.47209279E-03 q2bar2= 3.35684927E-05 q2bar= 6.64941253E-05

Mean sojourn times
tbar1= 4.97180748 tbar2= 3.43382263 tbar= 3.44402409

D statistics

Mean D1= 2.49487162E-03 Mean D2= -2.55113337E-05 Mean D=
-1.37759014E-06
Mean D1^2= 1.44610995E-05 Mean D2^2= 4.39678383E-09 Mean D^2=
1.42825883E-07

Mean pr1= 4.69955665E-07 Mean pr2= 2.43814320E-05 Mean pr=
2.48513879E-05
sig1d1= 5308.73828 sig1d2= -1.04634273 sig1d= -5.54331280E-02
sigd1= 3.63931608 sigd2= -5.16658463E-03 sigd= -2.76340608E-04
sig2d1= 30.7712002 sig2d2= 1.80333285E-04 sig2d= 5.74719952E-03
Theoretical value of unweighted sigma2d= 0.454544991

**Allele frequency weight= 100.000000**

Summary statistics for whole population

ns1= 6582 ps1= 6.58199983E-03
ns2= 993418 ps2= 0.993417978

Mean allele frequencies at 1st locus, conditioned on segregation
q1bar1= 1.60845723E-02 q1bar2= 7.17225298E-03 q1bar= 7.26220198E-03
Mean allele frequencies at 2nd locus, conditioned on segregation
q2bar1= 3.75833549E-03 q2bar2= 3.83188817E-05 q2bar= 7.58642782E-05

Mean sojourn times
tbar1= 5.27544832 tbar2= 3.42824078 tbar= 3.44039893

D statistics
Mean D1= 5.13599254E-04 Mean D2= -3.42951785E-06 Mean D=
1.78871085E-06
Mean D1^2= 6.00163787E-07 Mean D2^2= 2.44335819E-11 Mean D^2=
6.08147532E-09

Mean pr1= 4.04176781E-08 Mean pr2= 3.34322522E-06 Mean pr=
3.38364293E-06
sig1d1= 12707.2920 sig1d2= -1.02581120 sig1d= 0.528634608
sigd1= 2.55469275 sigd2= -1.87564327E-03 sigd= 9.72406531E-04
sig2d1= 14.8490419 sig2d2= 7.30838656E-06 sig2d= 1.79731590E-03
Theoretical value of unweighted sigma2d= 0.454544991

**Scaled selection coefficient= 50.0000000**
**Epistasis coefficient= 7.99999982E-02**

**Allele frequency weight= 10.0000000**

Summary statistics for whole population

ns1= 6561 ps1= 6.56099990E-03
ns2= 993439 ps2= 0.993439019

Mean allele frequencies at 1st locus, conditioned on segregation
q1bar1= 1.57506932E-02 q1bar2= 7.15647498E-03 q1bar= 7.23739108E-03
Mean allele frequencies at 2nd locus, conditioned on segregation
q2bar1= 3.41009744E-03 q2bar2= 3.24126195E-05 q2bar= 6.42148952E-05

Mean sojourn times
tbar1= 4.93644285 tbar2= 3.43007374 tbar= 3.43995690

D statistics
Mean D1= 2.45672301E-03 Mean D2= -2.50855865E-05 Mean D=
-1.71877764E-06
Mean D1^2= 1.40968696E-05 Mean D2^2= 4.16619494E-09 Mean D^2=
1.36852293E-07

Mean pr1= 4.49217566E-07 Mean pr2= 2.40020472E-05 Mean pr=
2.44512648E-05
sig1d1= 5468.89355 sig1d2= -1.04514360 sig1d= -7.02940151E-02
sigd1= 3.66545439 sigd2= -5.12035564E-03 sigd= -3.47591413E-04
sig2d1= 31.3809395 sig2d2= 1.73576656E-04 sig2d= 5.59694134E-03
Theoretical value of unweighted sigma2d= 0.454544991

**Allele frequency weight= 100.000000**

Summary statistics for whole population

ns1= 6638 ps1= 6.63799979E-03
ns2= 993362 ps2= 0.993362010

Mean allele frequencies at 1st locus, conditioned on segregation
q1bar1= 1.57254245E-02 q1bar2= 7.13476492E-03 q1bar= 7.21826730E-03
Mean allele frequencies at 2nd locus, conditioned on segregation
q2bar1= 3.57000786E-03 q2bar2= 3.50418304E-05 q2bar= 6.94024420E-05

Mean sojourn times
tbar1= 5.03570366 tbar2= 3.42827702 tbar= 3.43894696

D statistics
Mean D1= 5.17186709E-04 Mean D2= -3.43826196E-06 Mean D=
1.62227855E-06
Mean D1^2= 6.05572211E-07 Mean D2^2= 2.45350198E-11 Mean D^2=
5.91053517E-09

Mean pr1= 3.85859131E-08 Mean pr2= 3.35287132E-06 Mean pr=
3.39145731E-06
sig1d1= 13403.5107 sig1d2= -1.02546787 sig1d= 0.478342623
sigd1= 2.63289165 sigd2= -1.87771861E-03 sigd= 8.80911422E-04
sig2d1= 15.6941271 sig2d2= 7.31761475E-06 sig2d= 1.74277148E-03
Theoretical value of unweighted sigma2d= 0.454544991

**Selection coefficient= 2.50000004E-02**

**Scaled selection coefficient= 50.0000000**
**Epistasis coefficient= 0.159999996**

**Allele frequency weight= 10.0000000**

Summary statistics for whole population

ns1= 6580 ps1= 6.57999981E-03
ns2= 993420 ps2= 0.993420005

Mean allele frequencies at 1st locus, conditioned on segregation
q1bar1= 1.60006508E-02 q1bar2= 7.22421287E-03 q1bar= 7.30839418E-03
Mean allele frequencies at 2nd locus, conditioned on segregation
q2bar1= 3.60897533E-03 q2bar2= 3.49520051E-05 q2bar= 6.92335088E-05

Mean sojourn times
tbar1= 5.02401209 tbar2= 3.43604708 tbar= 3.44649601

D statistics
Mean D1= 2.58925138E-03 Mean D2= -2.52861209E-05 Mean D=
-2.08075846E-07
Mean D1^2= 1.55961607E-05 Mean D2^2= 4.20499058E-09 Mean D^2=
1.53759444E-07

Mean pr1= 4.76473076E-07 Mean pr2= 2.41835314E-05 Mean pr=
2.46600048E-05
sig1d1= 5434.20312 sig1d2= -1.04559255 sig1d= -8.43778625E-03
sigd1= 3.75106883 sigd2= -5.14188455E-03 sigd= -4.19010685E-05
sig2d1= 32.7325134 sig2d2= 1.73878274E-04 sig2d= 6.23517483E-03
Theoretical value of unweighted sigma2d= 0.454544991

**Allele frequency weight= 100.000000**

Summary statistics for whole population

ns1= 6538 ps1= 6.53799996E-03
ns2= 993462 ps2= 0.993462026

Mean allele frequencies at 1st locus, conditioned on segregation
q1bar1= 1.59683786E-02 q1bar2= 7.15202233E-03 q1bar= 7.23564904E-03
Mean allele frequencies at 2nd locus, conditioned on segregation
q2bar1= 3.38314893E-03 q2bar2= 3.23981621E-05 q2bar= 6.41817096E-05

Mean sojourn times
tbar1= 4.99219942 tbar2= 3.43075919 tbar= 3.44096804

D statistics
Mean D1= 4.92492225E-04 Mean D2= -3.43968054E-06 Mean D=
1.26443786E-06
Mean D1^2= 5.60235662E-07 Mean D2^2= 2.45523237E-11 Mean D^2=

5.33838573E-09

Mean pr1= 3.70208575E-08 Mean pr2= 3.35501591E-06 Mean pr=
3.39203689E-06
sig1d1= 13303.1016 sig1d2= -1.02523530 sig1d= 0.372766554
sigd1= 2.55962396 sigd2= -1.87789288E-03 sigd= 6.86542131E-04
sig2d1= 15.1329737 sig2d2= 7.31809450E-06 sig2d= 1.57379941E-03
Theoretical value of unweighted sigma2d= 0.454544991

**Selection coefficient= 2.50000004E-02**
**Scaled selection coefficient= 50.0000000**
**Epistasis coefficient= 0.319999993**

**Allele frequency weight= 10.0000000**

Summary statistics for whole population

ns1= 6625 ps1= 6.62499992E-03
ns2= 993375 ps2= 0.993375003

Mean allele frequencies at 1st locus, conditioned on segregation
q1bar1= 1.58472769E-02 q1bar2= 7.15644052E-03 q1bar= 7.23260501E-03
Mean allele frequencies at 2nd locus, conditioned on segregation
q2bar1= 2.95396894E-03 q2bar2= 2.61170462E-05 q2bar= 5.17763256E-05

Mean sojourn times
tbar1= 4.53418875 tbar2= 3.42025924 tbar= 3.42763901

D statistics
Mean D1= 2.16614990E-03 Mean D2= -2.50019020E-05 Mean D=
-5.79917742E-06
Mean D1^2= 1.13512760E-05 Mean D2^2= 4.14527612E-09 Mean D^2=
1.03588782E-07

Mean pr1= 3.58636697E-07 Mean pr2= 2.39380588E-05 Mean pr=
2.42966962E-05
sig1d1= 6039.95605 sig1d2= -1.04444146 sig1d= -0.238681734
sigd1= 3.61710525 sigd2= -5.11009013E-03 sigd= -1.17650232E-03
sig2d1= 31.6511841 sig2d2= 1.73166758E-04 sig2d= 4.26349230E-03
Theoretical value of unweighted sigma2d= 0.454544991

**Allele frequency weight= 100.000000**

Summary statistics for whole population

ns1= 6429 ps1= 6.42899983E-03
ns2= 993571 ps2= 0.993570983

Mean allele frequencies at 1st locus, conditioned on segregation

q1bar1= 1.52916806E-02 q1bar2= 7.16422591E-03 q1bar= 7.23643648E-03
Mean allele frequencies at 2nd locus, conditioned on segregation
q2bar1= 3.05123068E-03 q2bar2= 2.73532951E-05 q2bar= 5.42205344E-05

Mean sojourn times
tbar1= 4.75066090 tbar2= 3.42908263 tbar= 3.43757892

D statistics
Mean D1= 4.99626971E-04 Mean D2= -3.43950364E-06 Mean D=
1.03011189E-06
Mean D1^2= 5.76806656E-07 Mean D2^2= 2.45725697E-11 Mean D^2=
5.14913223E-09

Mean pr1= 3.37379689E-08 Mean pr2= 3.35701179E-06 Mean pr=
3.39074995E-06
sig1d1= 14809.0410 sig1d2= -1.02457297 sig1d= 0.303800613
sigd1= 2.72010946 sigd2= -1.87723804E-03 sigd= 5.59418113E-04
sig2d1= 17.0966625 sig2d2= 7.31977480E-06 sig2d= 1.51858211E-03
Theoretical value of unweighted sigma2d= 0.454544991

**Selection coefficient= 2.50000004E-02**
**Scaled selection coefficient= 50.0000000**
**Epistasis coefficient= 0.639999986**

**Allele frequency weight= 10.0000000**

Summary statistics for whole population

ns1= 6372 ps1= 6.37200009E-03
ns2= 993628 ps2= 0.993628025

Mean allele frequencies at 1st locus, conditioned on segregation
q1bar1= 1.52966846E-02 q1bar2= 7.17075961E-03 q1bar= 7.23434752E-03
Mean allele frequencies at 2nd locus, conditioned on segregation
q2bar1= 2.58833636E-03 q2bar2= 2.04176831E-05 q2bar= 4.05158171E-05

Mean sojourn times
tbar1= 4.23053980 tbar2= 3.43982768 tbar= 3.44486594

D statistics
Mean D1= 1.95530849E-03 Mean D2= -2.53295329E-05 Mean D=
-9.83050813E-06
Mean D1^2= 9.19222930E-06 Mean D2^2= 4.24322799E-09 Mean D^2=
7.61416814E-08

Mean pr1= 2.69761301E-07 Mean pr2= 2.42724782E-05 Mean pr=
2.45422380E-05
sig1d1= 7248.29150 sig1d2= -1.04354954 sig1d= -0.400554687
sigd1= 3.76465750 sigd2= -5.14126662E-03 sigd= -1.98435294E-03
sig2d1= 34.0754204 sig2d2= 1.74816436E-04 sig2d= 3.10247508E-03

Theoretical value of unweighted sigma2d= 0.454544991

**Allele frequency weight= 100.000000**

Summary statistics for whole population

ns1= 6645 ps1= 6.64500007E-03
ns2= 993355 ps2= 0.993354976

Mean allele frequencies at 1st locus, conditioned on segregation
q1bar1= 1.62277650E-02 q1bar2= 7.19113648E-03 q1bar= 7.26769771E-03
Mean allele frequencies at 2nd locus, conditioned on segregation
q2bar1= 2.61344621E-03 q2bar2= 2.23316019E-05 q2bar= 4.42847995E-05

Mean sojourn times
tbar1= 4.37908220 tbar2= 3.42825365 tbar= 3.43457198

D statistics
Mean D1= 4.37907700E-04 Mean D2= -3.43475995E-06 Mean D=
3.04461054E-07
Mean D1^2= 4.80223434E-07 Mean D2^2= 2.45098421E-11 Mean D^2=
4.09293754E-09

Mean pr1= 2.88844859E-08 Mean pr2= 3.35366440E-06 Mean pr=
3.38254904E-06
sig1d1= 15160.6543 sig1d2= -1.02418113 sig1d= 9.00093541E-02
sigd1= 2.57661939 sigd2= -1.87558436E-03 sigd= 1.65542573E-04
sig2d1= 16.6256523 sig2d2= 7.30837655E-06 sig2d= 1.21001573E-03
Theoretical value of unweighted sigma2d= 0.454544991

**Selection coefficient= 2.50000004E-02**
**Scaled selection coefficient= 50.0000000**
**Epistasis coefficient= 1.27999997**

**Allele frequency weight= 10.0000000**

Summary statistics for whole population

ns1= 6441 ps1= 6.44099992E-03
ns2= 993559 ps2= 0.993559003

Mean allele frequencies at 1st locus, conditioned on segregation
q1bar1= 1.51753444E-02 q1bar2= 7.20200175E-03 q1bar= 7.25957099E-03
Mean allele frequencies at 2nd locus, conditioned on segregation
q2bar1= 2.07508914E-03 q2bar2= 1.50917995E-05 q2bar= 2.99656676E-05

Mean sojourn times
tbar1= 3.83993173 tbar2= 3.42284560 tbar= 3.42553210

D statistics

Mean D1= 1.58269552E-03 Mean D2= -2.52581631E-05 Mean D=
-1.36484268E-05
Mean D1^2= 5.69049416E-06 Mean D2^2= 4.30531522E-09 Mean D^2=
4.53606894E-08

Mean pr1= 2.06486476E-07 Mean pr2= 2.42047900E-05 Mean pr=
2.44112780E-05
sig1d1= 7664.88721 sig1d2= -1.04351926 sig1d= -0.559103310
sigd1= 3.48298478 sigd2= -5.13394363E-03 sigd= -2.76240497E-03
sig2d1= 27.5586777 sig2d2= 1.77870388E-04 sig2d= 1.85818574E-03
Theoretical value of unweighted sigma2d= 0.454544991

**Allele frequency weight= 100.000000**

Summary statistics for whole population

ns1= 6529 ps1= 6.52900012E-03
ns2= 993471 ps2= 0.993471026

Mean allele frequencies at 1st locus, conditioned on segregation
q1bar1= 1.44868745E-02 q1bar2= 7.20852986E-03 q1bar= 7.25965900E-03
Mean allele frequencies at 2nd locus, conditioned on segregation
q2bar1= 1.94312271E-03 q2bar2= 1.37474408E-05 q2bar= 2.73017376E-05

Mean sojourn times
tbar1= 3.69382763 tbar2= 3.43142271 tbar= 3.43313599

D statistics
Mean D1= 3.96940857E-04 Mean D2= -3.43619695E-06 Mean D=
-6.23639153E-07
Mean D1^2= 4.17366323E-07 Mean D2^2= 2.45280636E-11 Mean D^2=
2.95625946E-09

Mean pr1= 2.05193000E-08 Mean pr2= 3.35987988E-06 Mean pr=
3.38039922E-06
sig1d1= 19344.7559 sig1d2= -1.02271426 sig1d= -0.184486836
sigd1= 2.77105117 sigd2= -1.87463255E-03 sigd= -3.39194958E-04
sig2d1= 20.3401833 sig2d2= 7.30027978E-06 sig2d= 8.74529709E-04
Theoretical value of unweighted sigma2d= 0.454544991
